# Structure-based learning to model complex protein-DNA interactions and transcription-factor co-operativity in *cis*-regulatory elements

**DOI:** 10.1101/2022.04.17.488557

**Authors:** O Fornes, A Meseguer, J Aguirre-Plans, P Gohl, PM Bota, R Molina-Fernández, J Bonet, AC Hernandez, F Pegenaute, O Gallego, N Fernandez-Fuentes, B Oliva

## Abstract

Transcription factor (TF) binding is a key component of genomic regulation. There are numerous high-throughput experimental methods to characterize TF-DNA binding specificities. Their application, however, is both laborious and expensive, which makes profiling all TFs challenging. For instance, the binding preferences of ~25% human TFs remain unknown; they neither have been determined experimentally nor inferred computationally. We introduce a structure-based learning approach to predict the binding preferences of TFs and the automated modelling of TF regulatory complexes. We show the advantage of using our approach over the state-of-art nearest-neighbor prediction in the limits of remote homology. Starting from a TF sequence or structure, we predict binding preferences in the form of motifs that are then used to scan a DNA sequence for occurrences. The best matches are either profiled with a binding score or collected for their subsequent modeling into a higher-order regulatory complex with DNA. Cooperativity is modelled by: i) the co-localization of TFs; and ii) the structural modeling of protein-protein interactions between TFs and with co-factors. As case examples, we apply our approach to automatically model the interferon-β enhanceosome and the pioneering complex of OCT4, SOX2 and SOX11 with a nucleosome, which are compared with the experimentally known structures.

## Introduction

Transcriptional regulatory elements are key players of the genome during development, cell and tissue homeostasis, responses to external stimuli, and disease^1^. Unravelling the mechanisms that regulate gene expression has consequently become one of the major challenges in Biology. With this objective the increase in the scale of experimental data, across multiple data types, has provided a plethora of activating regulatory elements of the genome^2^. Classical definitions of activating regulatory elements are focused in two classes: promoters (where transcription is initiated) and enhancers (elements that amplify such transcription initiation in cis, i.e. located within less than 1M bases distance of the initiation).

However, this distinction is becoming increasingly unclear, suggesting an updated model based on DNA accessibility of binding sites and enhancer/promoter potential ^1^. The sequence preferences of transcription factors (TFs) for these binding sites can be assessed by a wide variety of experimental techniques, both in vitro (such as SELEX^3, 4^, SMiLE-SEQ ^5^, protein-binding microarrays (PBM)^6–8^ and MPRA^9, 10^) and in vivo (such as bacterial and yeast one hybrid assays^11, 12^, ChIP-Seq^13^ and other high-throughput techniques^14, 15^). Recent models show similar potential of enhancers and promoters to promote the transcription machinery. Andersson et al.^1^ have pointed towards the TF and RNA polymerase II-centric cooperative model, in which regulatory elements work together to increase or maintain the local concentrations of transcription factors (TFs), RNA polymerase II (RNAPII), and other co-factors, thereby increasing the probability to target gene transcription start sites. Besides, it appears that very few proteins in humans occupy most of their motif matches under physiological conditions^16^, which highlights the importance of the balance between the co-operativity of TFs and their strength upon binding. Cooperative recognition of DNA by multiple TFs defines unique genomic positions on the genome and confers a systemic stability of regulation. Co-operative binding is most easily understood when it is mediated by protein-protein interactions that confer additional stability when two (or more) interacting proteins bind DNA^16^. Most eukaryotic TFs recruit cofactors as ‘‘coactivators” or ‘‘corepressors” forming large protein complexes to regulate transcription^17^. They commonly contain domains involved in chromatin binding, nucleosome remodelling, and/or covalent modification of histones or other proteins^16^. In the absence of direct protein–protein contacts between TFs, co-operativity can be mediated through DNA. Using CAP-SELEX^18^ Jolma et al.^19^ unveiled *in vitro* the co-operation of pairs of TFs through protein-protein and protein-DNA interactions. However, experimental protocols are both laborious and difficult to apply, and consequently most high-throughput efforts have been focused on a limited number of organisms.

Here, we have developed a structure-based learning approach to predict TF binding features and model the regulatory complex(es) in cis-regulatory modules (i.e. enhancers and promoters). Our objective is to characterize the role of structural elements, taking advantage of the recent developments on protein structure prediction (i.e. AlphaFold^20^) to reinforce both its modelling and prediction. Our approach integrates the experimental knowledge of structures of TF-DNA complexes and the large amount of high-throughput TF-DNA interactions to develop statistical knowledge-based potentials with which to score the binding capability of TFs in cis-regulatory elements. We have developed a server to characterize and model the binding specificity of a TF sequence or its structure. The server can automatically produce structural models of TF-DNA interactions and their complexes with co-factors. The approach is applied to the examples of interferon-β enhanceosome^21^ and the recent complex of “pioneer factors” SOX11/SOX2 and OCT4 with the nucleosome^22^. The model of interferon-β enhanceosome highlights the co-operativity of TFs with a more holistic view of domain-domain interactions that were missed in the experimental structures. The model of the pioneering regulatory complex locates OCT4, which is missing in the experimental structure, suggesting a potential role for nucleosome opening.

## Results

### A structure-based learning approach to score TF-DNA interactions

There are many methods to score the quality of protein folding^23–25^ and proteinprotein interactions^26, 27^, such as knowledge-based potentials, also known as statistical potentials^28–32^. In previous works we developed a set of statistical potentials^33^ to analyse protein structures and their interactions^34–36^. Ours is a structure-based learning approach that considers the frequency of contacts between pairs of residues and includes their structural environment, such as solvent accessibility and type of secondary structure, to evaluate the interaction between transcription factors (TFs) and nucleic-acids. It is optimized by grid searching to get the best parameters for each TF family. However, a limitation of this approach is the scarcity of known structures and in particular the scarcity of structures of protein-DNA interactions. To overcome this limitation, recently we developed a method for the C2H2 zinc-fingers (C2H2-ZF) family of TFs ^37^ that incorporated non-structural experimental information from systematic yeast-one-hybrid (Y1H) experiments^38^. Here, we have integrated experimental interactions from protein-binding microarrays (PBMs) for 37 TF families (plus some of their combinations) using the dataset of CisBP^39, 40^, notably increasing the landscape of protein-DNA contacts (see Figure 1; Supplementary Materials). The integration of experimental data from PBMs increased both the number and coverage for different types of contacts over many interval distances. For example, the use of PBMs data substantially increased the number of contacts for the AP2 and the homeodomain families; however, for the bHLH family the increase was more subtle (see Figure 1). We have named our approach ModCRE (*Mo*delling of *C*is-*R*egulatory *E*lements) and we offer a webservice for its practical use (https://sbi.upf.edu/modcre).

**Figure 1:**
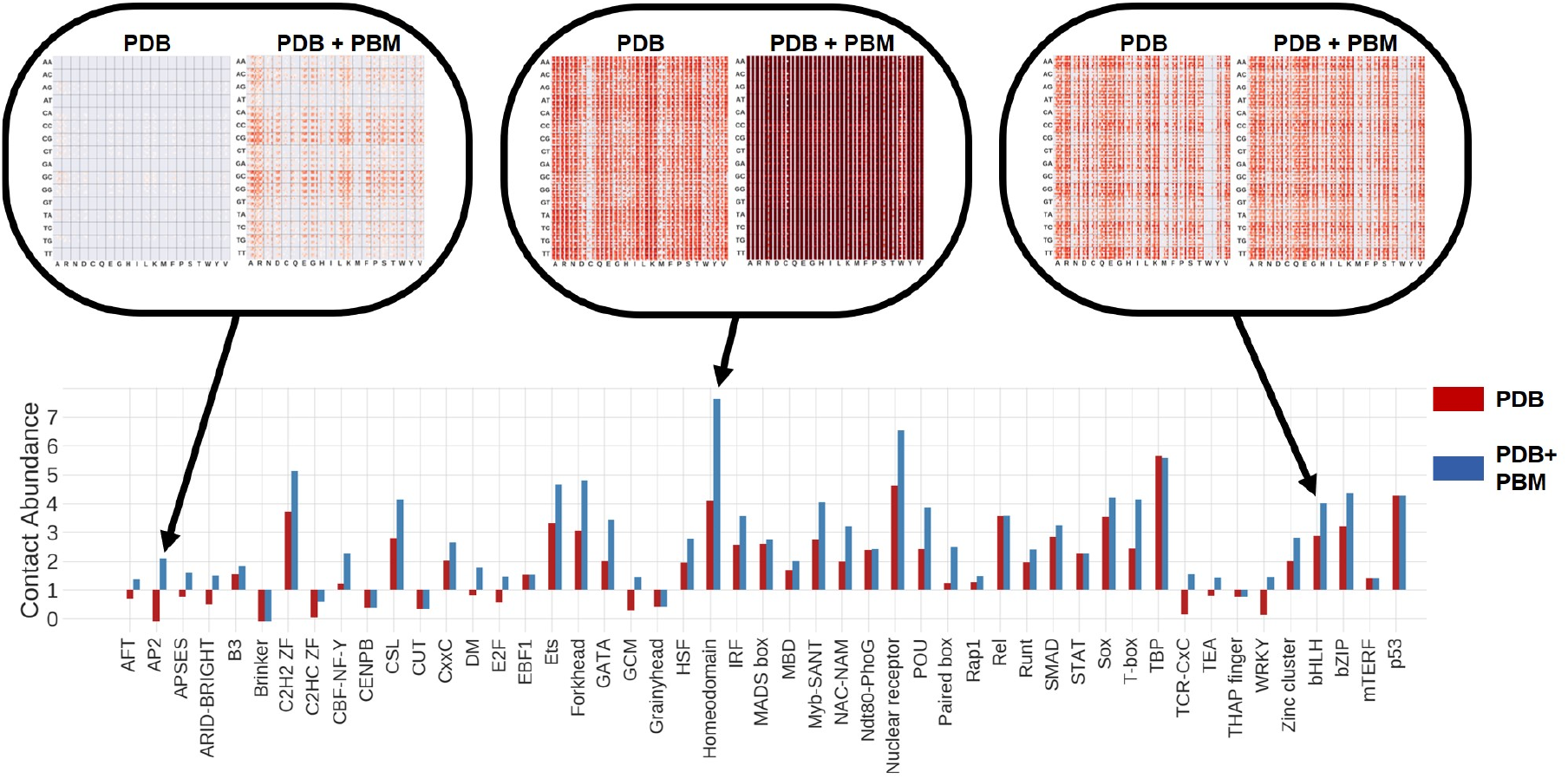
We defined the contact abundance score to capture the increase of coverage in the number and diversity of contacts thanks to the use of PBM experiments. This is defined as the logarithm of the ratio between the total number of potential accessible contacts and the number of contacts at less than 30Å. For the interaction between an amino-acid and two nucleotides there are 48 different types of contacts (considering the combination of all features, see supplementary methods), therefore the total number of potential contacts is 15360 (i.e. 4^2^ × 20 × 48), while the number of real contacts depends on the number occurrences in the known structures of a TF family and in the PBMs experiments that can be used to increase them. The figure shows the contact abundance score of each family calculated using only the known structures (PDB ^42^) or including the potential (modelled) contacts of other TFs derived from PBMs experiments (PDB+PBM). The figure shows in the top the heatmaps of the distribution of contacts, calculated with and without experimental PBMs data, for three types of families (AP2, for which the use of PBMs significantly increases the coverage; bHLH, for which the increase is not relevant; and the homeodomain family for which the coverage was already very large with only data from PDB). The labels and details of the heatmaps are shown as example in Figure S1. All heatmaps can be downloaded from http://sbi.upf.edu/modcre/##faq.

### ModCRE predicts TF binding preferences

We propose two tests to evaluate the predictive power of ModCRE: 1) evaluate the capacity to classify DNA 8-mers as bound/unbound for all TFs of the PBMs experiments and specifically for the TFs of each family; and 2) evaluate the capacity to predict the DNA binding motifs of the TFs in the JASPAR dataset^41^ (which are described by means of position weight matrices, i.e. PWM), analyzing the results by TF families. Our objective is to select the binding region of a TF with the purpose of automatically modelling the structure of protein-DNA complexes; therefore, our tests are addressed to check the capacity of recognizing the binding site rather than identifying the critical nucleotides that may be affected by mutations (i.e., characterizing the relevance of each position in the binding site). This goal is achieved by selecting the correct binding or predicting a PWM sufficiently similar to the experimental PWM of a TF target. Further analysis of the binding region and the role of each nucleotide and its position can be addressed in the webserver using the predicted PWMs to scan a DNA sequence uploaded by the user (see in “Characterizing/Identifying the binding sites of TFs”).

First, we tested the structure-based potential (i.e. named in short *ZES3DC_dd_* in methods and supplementary material) to discern positive (binding) from negative (non-binding) 8-mers as described in the PBMs experiments of the CisBP database (version 2.0) ^40^ (see methods for details). We avoided redundancies within the TFs of each family by filtering out all other TFs of the same family with more than 70% identical contacts in the interface, and 40% for the general potential with independence of the TF family. We applied a 5-fold cross validation to train the potentials and test. Taking this into account, each TF was not tested with the statistical potential of itself or a too close homolog (family-specific potentials are defined per structure and fold, not by sequence). The purpose of this test was to understand the role of the features used to describe the knowledge-based potential. Despite this filtering, a relevant number of TFs in the training set had similar interfaces in the testing set of the family-specific potentials. However, the number of TFs in the test of some families was already too small and we could not use a more stringent cut-off without a dramatic loss of applicability. We scored the interaction of each TF with all their positive and negative 8-mers by modelling the TF-DNA complexes. We obtained precision-recall curves and calculated the area under the curve (AUROC and AUPRC) to compare TF families and the features characterizing the statistical potential (Figure 2). Interestingly, some features were better than others depending on the family. Consequently, we designed a grid-search protocol to optimize the best features for each family of TFs to predict their PWMs (see methods).

**Figure 2.**
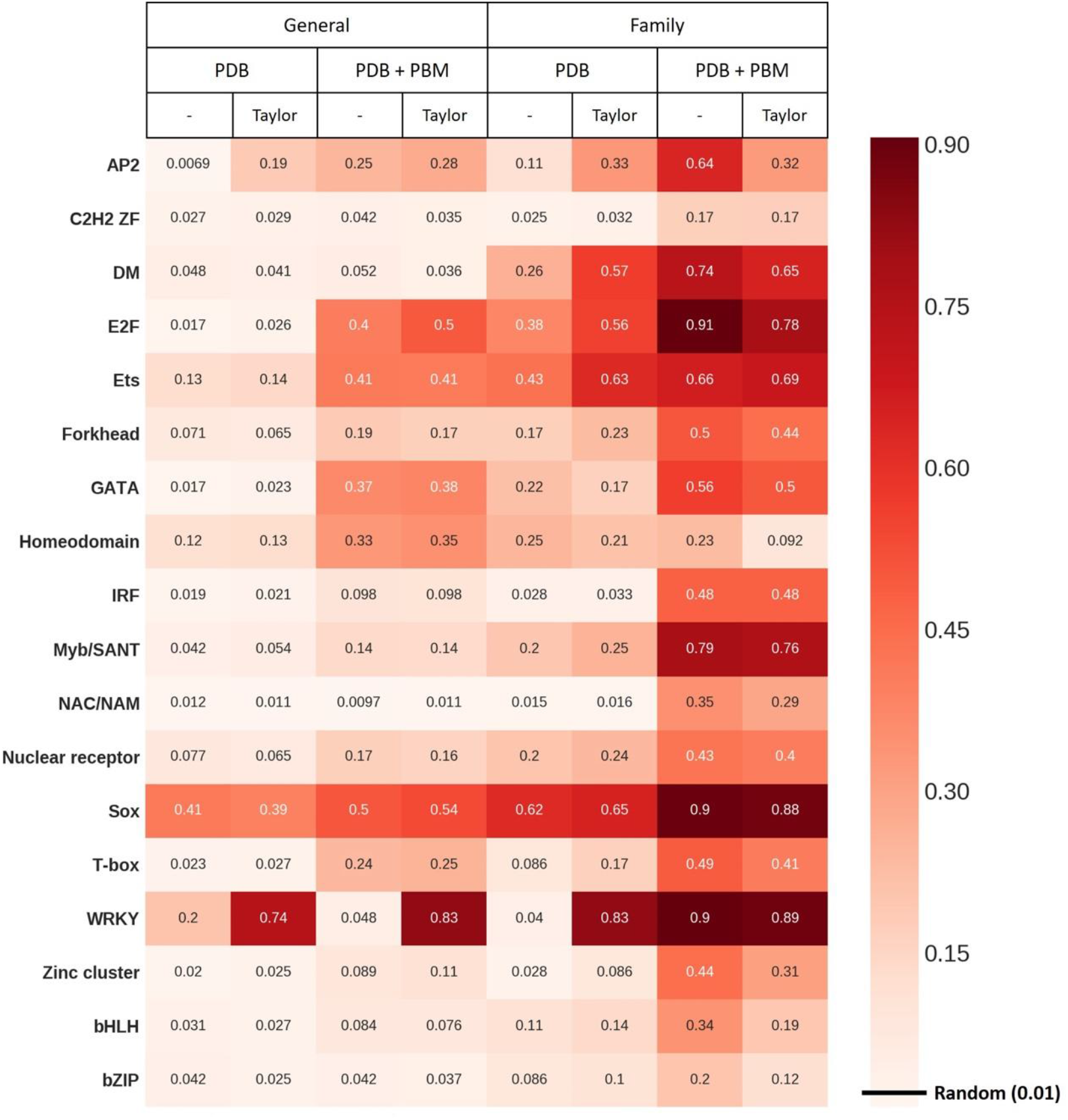
Area under the curve of precision-recall (AUPRC) on the prediction of positive and negative 8-mers of the PBMs experiments. We analyzed TFs of all families from Cis-BP database with PBMs experiments. We used the PWMs predicted with structural models of TFs using different features to calculate the ZES3DC_dd_ statistical potential (i.e. using contacts extracted from PDB or from PDB plus those derived from PBMs experiments, using only contacts obtained with TFs of the same family, or using a Taylor polynomial approach to complement the missing contacts in the experimental data). Not all negative 8-mers were used, forcing the ratio of positive/negatives to be 1 to 100 (negative 8-mers were selected randomly). We restricted the study to those families with at least 10 different TFs to sufficiently support the results (the test of the rest of families is in supplementary figure S2). The results of the area under the curve of true and false positive rates (AUROC) for all TF families are shown in Figure S2.

Second, we used these optimized parameters to predict the motifs of TFs in JASPAR dataset^41^ which structure could be modelled (most PWMs in this set were obtained by SELEX instead of PBMs, thus representing an independent testing set). The structural models of many TFs were obtained with different templates (i.e., using several known structures of close homologs interacting with a double-strand DNA helix). Therefore, to equilibrate the number of models, we limited to 100 the number of models for each TF, obtained by using all available templates from PDB^42^ and/or generating several conformations with MODELLER^43^. We used the version of JASPAR 2020 consisting of 1934 PWMs. After discarding TFs with more than one PWM to avoid misinterpretation of predictions, the final dataset contained 1210 TFs (i.e. approximately 62% of the original dataset). We predicted 100 PWMs per TF and compared them with the experimental motif from JASPAR using TOMTOM^44^. We obtained the *P*-*value* provided by TOMTOM from each comparison and transformed it into a measure of similarity (*similarity score*) defined as −*log*_10_(*Pvalue*). Figure 3 shows the results of the prediction for several families of TFs by plotting the average (in Figure 3A) and the best scores (in Figure 3B) out of 100 models of each TF. For most TFs (around 75%) we predicted at least one motif significantly similar to the experimental (P-value < 0.05). However, we must note that the criterion of P-value from the comparison of two PWMs may have some intrinsic dependencies (for example on the size of the two PWMs or in the variance of DNA binding sequences for different TF families); thus, it is not the best criterion to compare results from different TF families. Besides, we notice that having a good match out of 100 doesn’t imply the other 99 are either good or bad. This only ensures that the method can find at least one good match. Therefore, in the next section (“ModCRE predicts well in the twilight zone”) we will use a different criterion, independent of the TF families and the size of their PWMs, to score these comparisons. Then, we will use the score-distributions to avoid the problem of the selection of a single motif per TF (see further).

**Figure 3.**
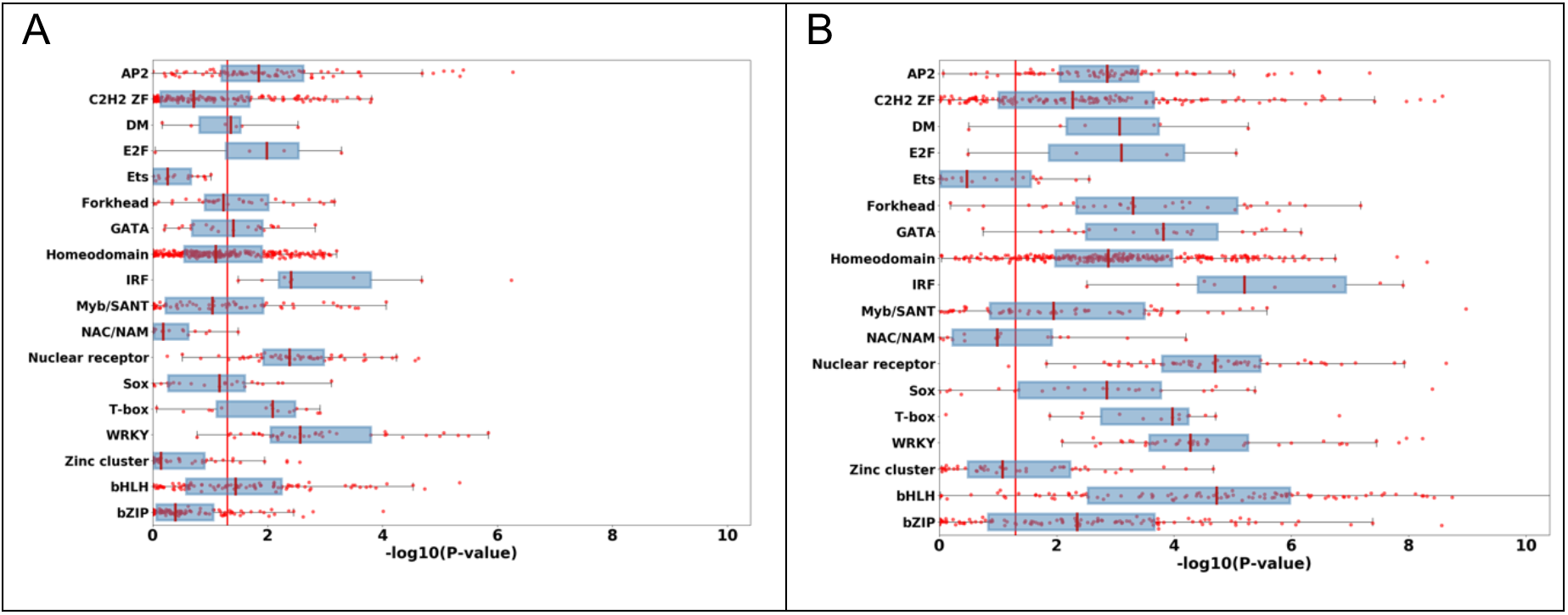
Distribution of “similarity scores” to compare predicted and experimental PWMs of TFs. The score of similarity is defined as −log_10_(Pvalue), where the Pvalue is obtained with TOMTOM and it shows if the alignment of the two PWMs is significant (i.e. Pvalue is the probability that a random motif of the same width as the experimental PWM would have an optimal alignment as good or better than the PWM predicted with the structure of the TF). Each red dot in the plot shows either the average of scores (in A) or the highest score (in B) of the comparison of 100 predicted PWMs obtained with the structural models of each TF. Boxes in blue show the best quartiles of the distribution for each TF highlighting in red the mean of the distribution for all TFs of a family. A red line indicates the threshold at which the predicted PWM is significantly similar to the experimental (i. e. Pvalue<0.05). For TFs of the C2H2-ZF family we used the parameters and statistical potentials derived from a previous work^37^. As in Figure 2, we restricted the study to those families with at least 10 different TF sequences (the comparison with the rest of families of TFs is in supplementary figure S3).

We studied more deeply the prediction of 177 TFs that have motifs in JASPAR and in CisBP (as obtained with PBMs). These TFs have 295 motifs in CisBP and 213 in JASPAR, which is about 20% of the original data with representation for most TF families. Considering that some TFs have more than one motif in JASPAR and in CisBP, the total number of comparisons between motifs was 369. All families except for TCR/CxC had at least one TF for which one model produced a PWM like the experimental (in JASPAR and CisBP). Table 1 shows the predicted motifs of a selected set of TFs compared with the experimental ones from JASPAR and CisBP. Despite our approach having learned the parameters of the predictive model using data from CisBP, the flexibility introduced with the variety of structural models helped to achieve good predictions of JASPAR motifs. In supplementary Table S2 is shown a summary of the predictions for these TFs. From Table S2, considering successful the prediction for a TF if more than 50% of the predicted PWMs are significatively like the experimental motif, we correctly predicted almost 48% TFs’ motifs compared with JASPAR and 57% with CisBP. Then, if we consider successful the prediction for a TF if at least one of the predicted PWMs is significatively similar to the experimental motif, we predict 82% motifs compared with JASPAR and 89% with CisBP. In the next section we will apply these ideas to develop a new algorithm, like a jury-vote approach, to improve the accuracy and help in the selection of a single PWM and the detection of a binding site. Two facets of this validation with JASPAR must be noted: 1) We test the sequence of a TF blindly; therefore, although we have avoided TFs with more than one motif in JASPAR, the structural model can be produced by partial structures combining more than one domain (e.g., for sequences of the C2H2-Zf family with several domains). Then, an issue for this test is that for some TFs a portion of their models might correspond to regions of the protein addressed to a different binding site than the motif under test and consequently the comparison will fail. In addition, some structures contain more than one binding pose and some correspond to incomplete binding sites caused by the crystal (e.g. for the homeodomain family, the crystal structures of 2HOT, 2HOS, 1HDD, 2HDD, 3HDD, and 1DUO from PDB show two different poses of the homeodomain binding, one of them incomplete), yielding the same problem. We assumed these problems as failures of the method because the region of the protein cannot be selected before the comparison with the real motif is done (we could compare the predicted PWMs and cluster them; nevertheless, selecting one or another cluster would still be random). 2) The approach searches in the database of structures of TF-DNA complexes to obtain the structural model(s) of a TF. Therefore, finding a sufficiently similar TF sequence automatically imposes a specific conformation and sequence of the DNA. We can further stress this point by simply searching on a database of TF sequences with known DNA binding sites or motifs. This is known as nearest-neighbor approach. In our validation, we neglected the DNA sequence of the template, but we assumed the same conformation in the model as in the template. A relatively large number of TFs preserve a B-DNA conformation (where subtle differences are unnoticed by the coarse-grained potential of our approach), but for some others, such as the TATA-box, the conformation in the template may be determinant of the sequence of the binding site. Therefore, it is more appropriate to validate ModCRE by comparison with the nearest-neighbor (see next section). Finally, if we plan to use our approach on novel complexes without previous knowledge of the structure of the TF-DNA interaction, then the role of the DNA conformation must be considered, as this affects the quality of the prediction.

**Table 1.**
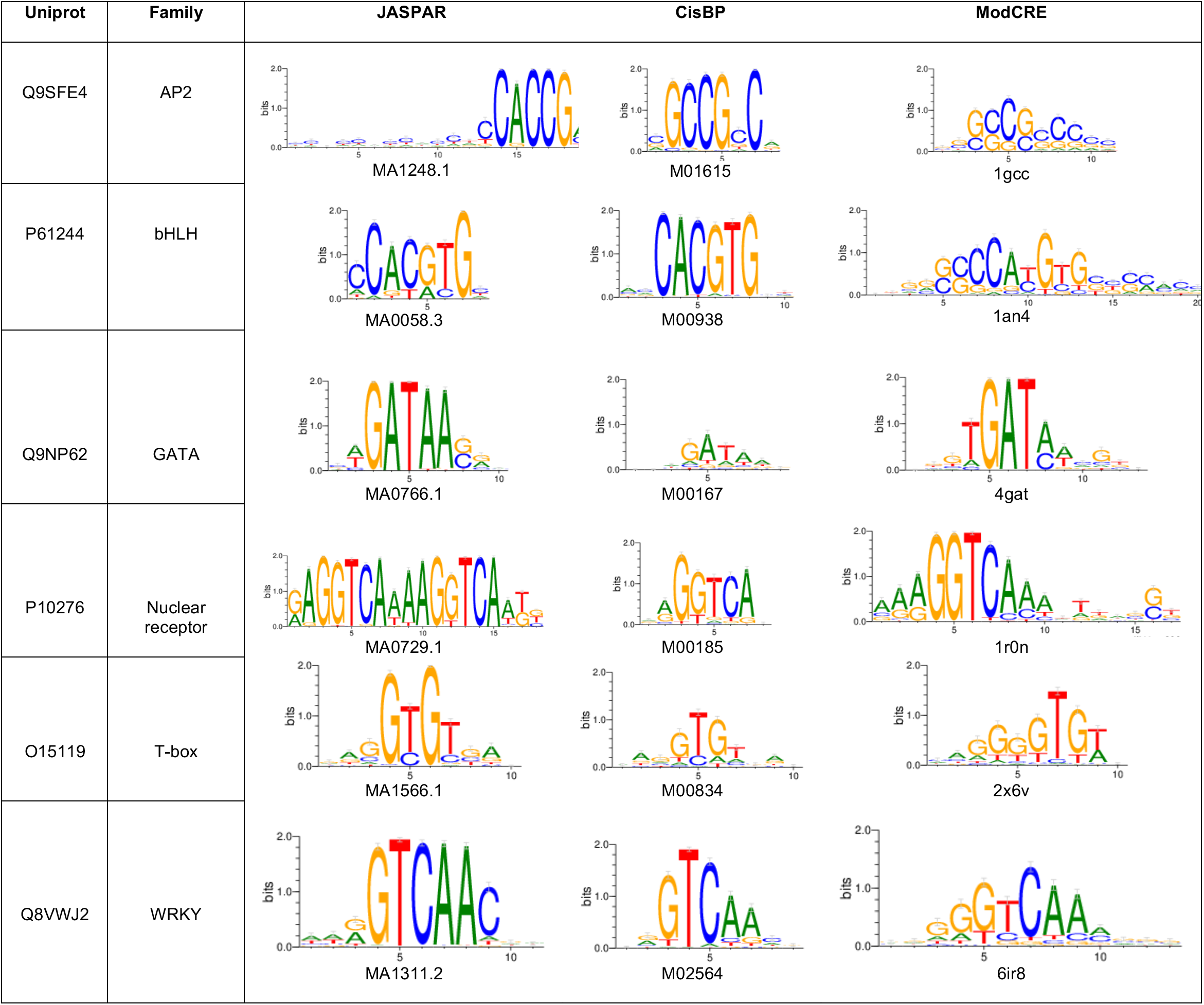
Logos of experimental PWMs (from JASPAR and CisBP) and of their best predictions. The table shows the logos of only some TFs for which the PWMs were also obtained by PBMs in CisBP. The TF is identified by the UniProt code. The code of the PWMs corresponding to the logos in JASPAR and CisBP are shown at the bottom. For the logos of the predicted PWMs the PDB code of the protein used as template is shown at the bottom. The logos of TFs from other families is shown in supplementary Table S1.

| Uniprot | Family | JASPAR | CisBP | ModCRE |
| --- | --- | --- | --- | --- |
| Q9SFE4 | AP2 | MA1248.1 | M01615 | 1gcc |
| P61244 | bHLH | MA0058.3 | M00938 | 1an4 |
| Q9NP62 | GATA | MA0766.1 | M00167 | 4gat |
| P10276 | Nuclear receptor | MA0729.1 | M00185 | 1r0n |
| O15119 | T-box | MA1566.1 | M00834 | 2x6v |
| Q8VWJ2 | WRKY | MA1311.2 | M02564 | 6ir8 |

### ModCRE predicts well in the twilight-zone

We simultaneously compared and validated the prediction of the PWM with the state-of-the-art approach based on sequence homology. This approach is also known as prediction by nearest-neighbor. The nearest neighbor approach consists on using the experimental PWM of the closest homolog of a TF^45^. The accuracy of such prediction depends on the degree of similarity between TFs: hypothetically, close homologs should have similar DNA binding domains and in consequence their PWMs should be similar too. As dataset, we used the TFs of CisBP that had been studied by PBMs. First, we compared their sequences using MMseq2^46^. Then, for each target sequence we grouped the other TFs of the dataset by sequence similarity with the target. Each group contains sequences that align with the target between a minimum and a maximum percentage of identical residues defined by the group. The groups range between 15% and 95% binned in intervals of 10% (e.g., the group at 95% contains all closest homologs of a target TF which alignment produces a percentage of identical residues between 90% and 100%). The analysis of the prediction was performed by grouping TFs by families and the results by bins of the same interval of sequence similarity. Notice that it was not always possible to find relatives in all bins for all TFs. The total number of TFs in each bin and family varies, as well as the total number of predictions (homologs with a known PWM). Table S3 shows the number of TFs for which the nearest-neighbor approach can be applied and the total of predictions for each bin and family. For ModCRE, given a family and a bin, we used the same TFs that were tested in the nearest neighbor approach, for which we modelled 100 conformations (yielding 100 motifs). Then, instead of using the P-value as a criterion to test the predictions, we tested the success by ranking: we used TOMTOM^47^ from the MEME suit^44^ to compare the predicted PWMs with all the experimental motifs from CisBP, including the motif that corresponds to the target, and ranked them according to TOMTOM score (from best to worst). The rank of the actual motif of the target indicates the quality of the prediction. We scored and normalized the rank to fit between 0 and 100 (then the highest score is achieved with the best rank). The normalized score is defined as: 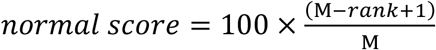, where M is the total amount of PWMs of the dataset. The normalized score is null if the Pvalue of the comparison between the predicted PWM and the actual motif is higher than 0.05 (i.e. non-significant). The prediction for all TFs is produced by the accumulation of results for all families in each bin (see Figure 4).

**Figure 4.**
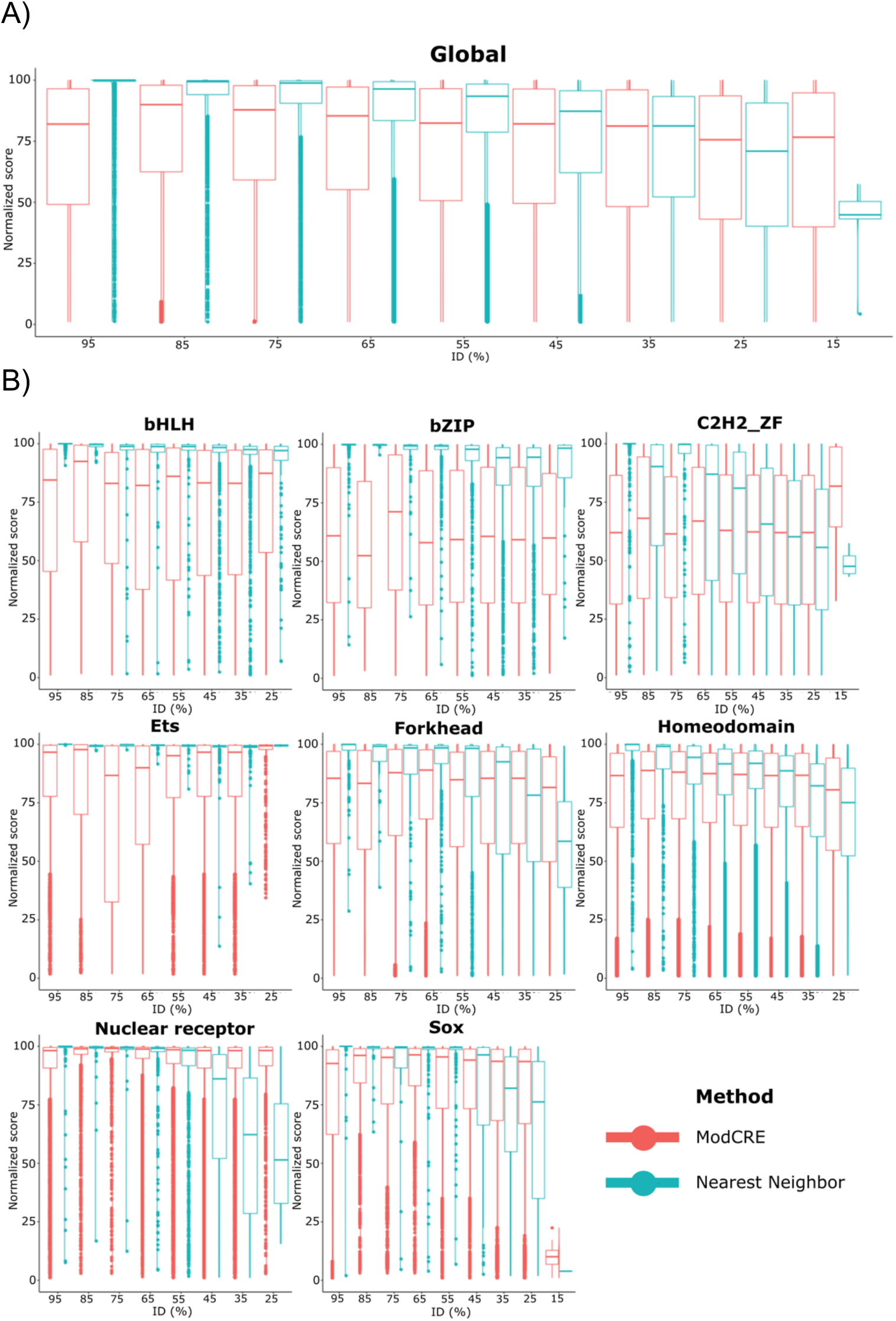
Distribution of the normalized ranking score of motif predictions with the nearest-neighbor (state-of-the-art) approach and the structure-based approach (ModCRE). Results with the nearest-neighbor approach are shown in blue and results with the structure-based approach in red. The top distribution (A), entitled Global, corresponds to the distributions for all TFs (i.e. 2268 TFs, with 2638 motifs) in the dataset of CisBP with PBM experiments. The rank of the correct motif was scored and normalized with respect to the total size of the database of compared PWMs, yielding a normalized score. The normalized score is defined as: normal score = 100 × (2638 – rank + 1)/2638. The total number of TFs, motifs and predictions in each bin are shown in supplementary table S3. Tables of acceptable errors of the ranks are shown in tables S4 as a function of the average of the number of neighbors of the TFs that are significatively similar (see details in tables S4). The distributions of normalized scores of some families of TFs are also plotted in B (the title of each plot indicates the name of the family). Plots of distributions of normalized scores for the rest of families can be downloaded from web (http://sbi.upf.edu/modcre/##faq) and supplementary S7.

The comparison between the nearest-neighbor and ModCRE clearly shows that the latter outperforms the state of the art (nearest-neighbor) at low sequence identity. For example, ModCRE predictions in the twilight zone, around 15-25% sequence identity, outperformed the nearest-neighbor approach for TF families such as C2H2 ZF, ETS and Homeodomains. For some families ModCRE outperformed the nearest neighbor approach at bins around 50% of sequence identity (i.e. families such as Forkhead, Nuclear Receptor and SOX). Nevertheless, for some families such as bHLH and bZIP, the nearest-neighbor method was still the best approach to predict their binding preferences, because these were preserved by distant and remote homologs. This was also reflected in the poor increase of new amino-acid and nucleotide contacts incorporated from PBMs experiments in the statistical potentials, showing that both TF families have a limited landscape of protein-DNA contacts that was easily covered by evolution. Interestingly, Lambert et al.^40^ had already predicted by similarity regression the similarity in DNA sequence specificity between two TFs (specially for members of the Homeodomain family), showing the relevance of the amino-acids in contact with the DNA, and helping to unveil the preservation of the binding motif between remote homologs. Lambert et al.^40^ also noticed many highly or ambiguously similar homolog TFs in the bHLH and bZIP families, appearing rigid in their DNA-binding motifs and suggesting that TFs of these families had diversified through changes in heterodimerization partners. Consequently, we note that an acceptable error of the rank must be considered for each TF family specifically, because many TFs have PWMs like the motifs of other TFs in the same family. Hence, depending on the number of neighbors with similar PWM and how such similarity is defined, the rank can be confused with those of other very similar motifs. Supplementary tables S4 show the acceptable errors of the rank for several families of TFs. Acceptable errors in tables S4 are shown as a function of the P-Value, Q-value and E-value to qualify if two PWMs are significantly similar. This helps us to determine the quality of predictions by nearest-neighbor and ModCRE approaches. Therefore, the evaluation by ranking permits us to compare the results of different TF families with independence of the size and variance of the PWMs of their TFs, while the margins of errors enable us to qualify the quality of the comparison with the predictions.

### On the improvement of the prediction of binding preferences of TFs

As observed in the structure-based prediction of ModCRE, we generated several PWMs for a target TF. Having a successful prediction out of 100 models is useless when we don’t know the real PWM of the target. However, if a relevant number of models points to the same PWM (e.g. more than 50%), we could take this as the final prediction (e.g. by a majority vote selection). This suggests a potential new approach to predict the motif of a TF that can also be applied to the nearest-neighbor approach. We propose as solution (i.e. the predicted motif) the most often selected motif among the best rankings. For the nearest-neighbor approach, instead of selecting the PWM of the closest homolog, we consider all the motifs of TFs with sufficiently similar sequence to the target. Thus, a collection of motifs is used as in the structure-based approach. We name the approach “rank-enrichment prediction”.

As in the previous ranking of motifs of the dataset (which includes the actual motif of the TF target), we rank by the score of TOMTOM the motifs of the database for all the predicted PWMs of the target. We remove all non-significantly aligned PWMs and select a limited number of top solutions. The number of potential solutions selected affects the quality of the prediction, i.e. if we use too many the success is not significant, or in other words, it can be achieved at random (see more details in the supplementary material). In the selected set, some motifs may have been included several times. Then, we calculate the enrichment of a motif as the ratio of the number of times it appears in the selection. The final prediction (i.e. solution) corresponds to the motif with highest enrichment (i.e. the motif that was more often selected, either among homologs of the target in the nearest-neighbor approach, or among modelled conformations in the structure-based approach). Finally, to evaluate the quality of the predicted motif, we compare it with the motifs of the database and calculate the ranking of the experimental motif of the target. If the motif of the target is not selected among the potential solutions, then the ranking cannot be calculated and the prediction is removed, affecting the coverage of predictions (i.e. the sensitivity to find solutions). Besides, depending on the number of models used for the enrichment and the total number of acceptable correct solutions (sufficiently similar PWMs), the prediction may not be significant. Thus, nonsignificant predictions are also removed.

The ranking is scored and normalized as before, and the distribution is plotted for several bins of sequence-similarity. We must note that now a single solution is proposed for each TF in both approaches, nearest-neighbor and structure-based with ModCRE. Figure 5 shows the distribution of the normalized ranking on the application of rank-enrichment prediction using ModCRE and the nearest-neighbor. The rank-enrichment increases the accuracy of the predictions of both methods. ModCRE often achieves better coverage than nearest-neighbor at low percentages of similarity, while preserving most ranking scores at the top (around 98%). Supplementary tables S4 shows the acceptable errors in the normalized score as function of the P-value, E-value and Q-value for all families (e.g. when using all families, the average number of similar motifs with Q-value < 10^-3^ is around 60, and this implies a rank of about 98%, which is also in the same rank and with about the same number of similar motifs with P-value <10^-7^). The acceptable errors for families such as Nuclear Receptors, C2H2-Zf, GATA or SOX are smaller than for Homeodomains. Interestingly, the enriched ranks of ModCRE predictions for TFs of Nuclear receptors and SOX families are high and the coverage is improved with respect to the nearest-neighbor approach.

**Figure 5.**
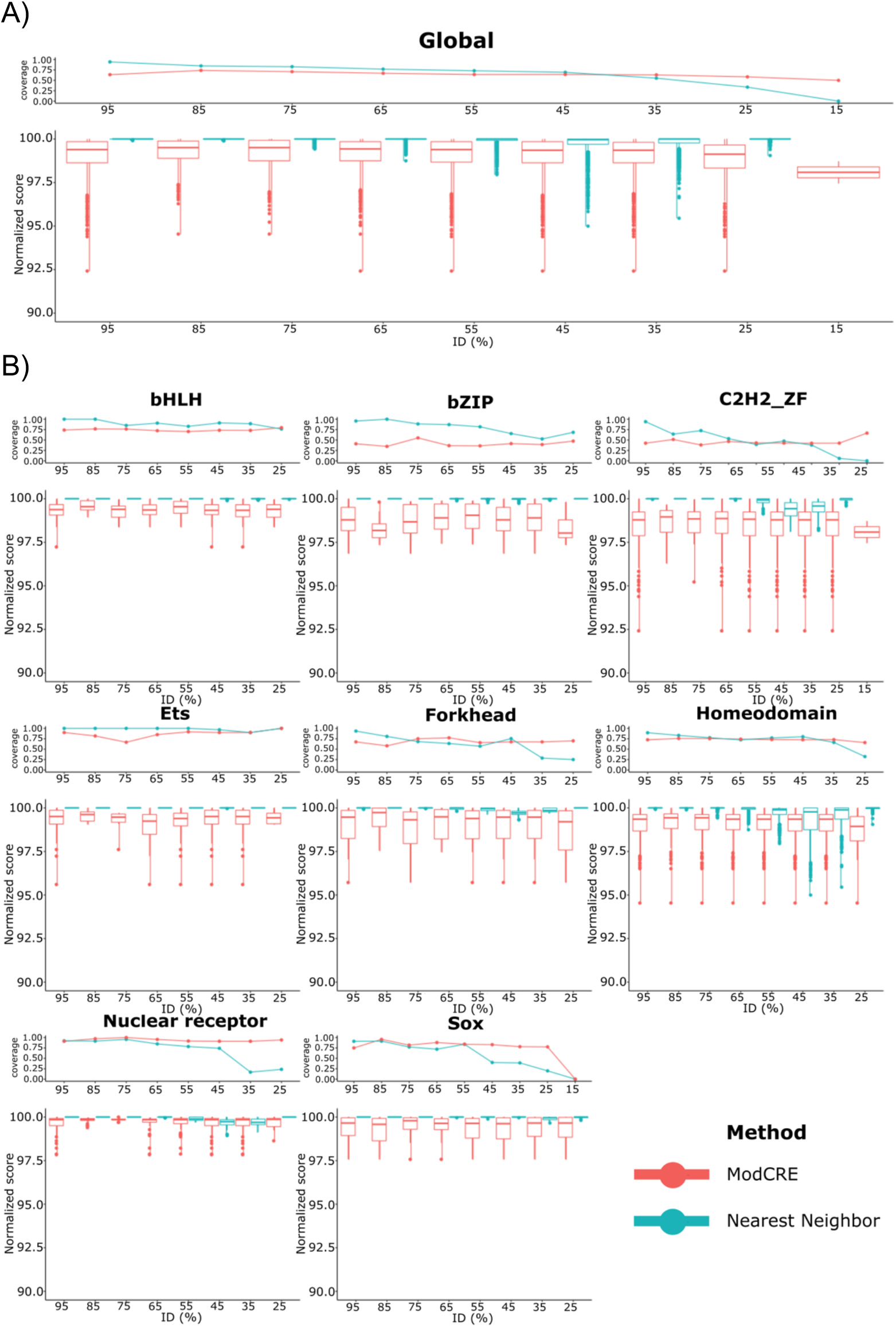
Distribution of the normalized ranking score of motif predictions with the nearest-neighbor (state-of-the-art) approach and the structure-based approach (ModCRE) using rank-enrichment. Results with the nearest-neighbor approach are shown in blue and results with the structure-based approach in red. In A are shown the results for all TFs and in B the results of some TF families. Plot titles and normalized scores are defined as in Figure 4. On top of each plot are shown the curves of coverages after removing non-significant predictions. Coverages are calculated as the ratio of significant predictions over the total of predictions in each bin. The total number of TFs, motifs and predictions in each bin are shown in supplementary table S3. The distributions of normalized scores of some families of TFs are plotted as in Figure 4. The acceptable errors of normalized ranks are the same as in Figure 4. Plots for the rest of families can be downloaded from web (http://sbi.upf.edu/modcre/##faq) and supplementary S7.

### Characterizing/Identifying the binding sites of TFs

In practice, unless the binding preferences of a TF have been experimentally tested, there is no access to the real motif(s) of a TF and consequently the rank-enrichment approach cannot be applied directly. However, the approach indicates that the best prediction is produced by a majority of similar motifs out of a collection of predicted PWMs. Therefore, a sensible approach to predict the binding site of a TF in a DNA sequence is: 1) to predict a set of PWMs, either by nearest-neighbor or with the structure-based approach; the latter if the homologs of the target are not sufficiently similar and the former if the percentage of sequence identity with the homolog is higher than 50%; 2) scan the DNA sequence with all predicted motifs using FIMO^48^; and 3) select the fragment matched by the majority of motifs with a significant score.

To analyze the binding sites of a DNA sequence, we have developed ModCRE as a web server that predicts the PWM of a TF based on its structure or by modelling several conformations with its sequence. Then, the DNA sequence can be scanned with the PWMs and either the accumulation of matches can be profiled (i.e. using a score for the prediction of binding), or a selection of matches can be collected to build the structure of a cooperative binding (i.e. considering the formation of a potential complex of transcription). The webserver permits the substitution of the predicted PWM by another (in MEME format). This is more convenient if we know experimentally the PWM or if we know the PWM of a close homolog of the target (as it had been shown in the previous sections). To facilitate the scanning of a large DNA sequence and the construction of a structural model of cooperative binding, we have included the PWMs of three datasets associated with TF structures. Two datasets are defined with the experimental motifs from JASPAR and CisBP. In each of these sets a motif is associated with the sequence of a TF. Therefore, we have modelled the potential conformations of each one of these TFs and selected the model with the PWM most similar to the experimental motif. Then, a target DNA sequence is scanned with FIMO using the motifs of the corresponding database and the associated models can be selected in ModCRE web server. Similarly, the third database is obtained using the structures of TFs complexed with a DNA double strand: we use BLAST^49^ and HMMER^50^ to obtain the sequences of potential homologs in UniProt and TrEMBL^51^ that align (without gaps in the binding interface) with the sequences of these TFs. A motif is predicted for any of these sequences using the alignment and the template structure. Consequently, we can scan the DNA with any of the sequences of specific species (see further details in supplementary material). Additionally, specific TF sequences can be uploaded to predict their motifs and scan the DNA.

### Integration of transcription factors and co-factors in a regulatory complex

To complete the modelling, co-factors can be included in the network of interactions when the species selected are human or mouse. Interactions between TFs and transcription co-factors (TcoFs) are retrieved from the TcoF-DB database^52^. After selecting a set of protein-protein and protein-DNA interactions, these can be modeled using a homology modeling pipeline^26, 35^. Then, ModCRE models the structure of DNA in a specific conformation (B conformation by default), and for very long DNA sequences the server splits the sequence in fragments of 250 base-pairs (with an overlap of 50bp to be able to assemble them later). Models with clashes between proteins are removed and only acceptable combinations of each fragment are selected to construct a model. Next, the structures are optimized by several steps of conjugate gradient and short annealing dynamic simulations with MODELLER. Finally, distance restraints are extracted from the models of proteinprotein interactions and TF-DNA interactions, and we use the package IMP^53^ to integrate them in a model of DNA with all TFs and TcoFs (see more details in supplementary material). We are not aware of any other web-service method to automate the structural modeling of these complexes and very few experimentally known complexes to benchmark. The most similar approach was recently described if RoseTTAFoldNA ^54^.

### Example 1: the interferon-beta (IFN-β) enhanceosome

We have used the server to automate the modelling of the interferon-beta (IFN-β) enhanceosome, an ensemble of TFs and Cis-regulatory elements that cooperate in the enhancer of the IFN-β gene^21, 55^. The TFs binding at the IFN-β enhanceosome are ATF-2, c-Jun, IRF-3, IRF-7, and NFKβ-1 (subunits p105 and p65). We have used a sequence of 250bp containing the region of the enhanceosome and the database of human TFs, using their predicted PWMs based on their modelled structures, to predict and model the co-operative complex. These PWMs are used to scan the DNA sequence with FIMO. In supplementary Figure S4 we select the bindings of the specific TFs (i.e. ATF-2, c-Jun, IRF-3, IRF-7, and NFKβ-1) that are significant (P-values < 5.0e^-4^). Hence, we are able to recreate a structural model similar to the model provided by Panne^21, 55^ based on experimental data. On the binding sites predicted for NFKβ-1 (subunits p105 and p65) the automated approach produces a homodimer of NFKβ-1 with two subunits p105 instead of a heterodimer with RelA (subunit p65). Not all the binding regions of IRF-3 are detected exactly, but some other regions are predicted instead. Besides, the IRF-7 binding site from Panne’s model is occupied by IRF-3. The analysis highlights the accumulation of TFs in a short section of the DNA and brings a potential explanation for the formation of the transcription complex by gathering TFs (Figure 6).

**Figure 6.**
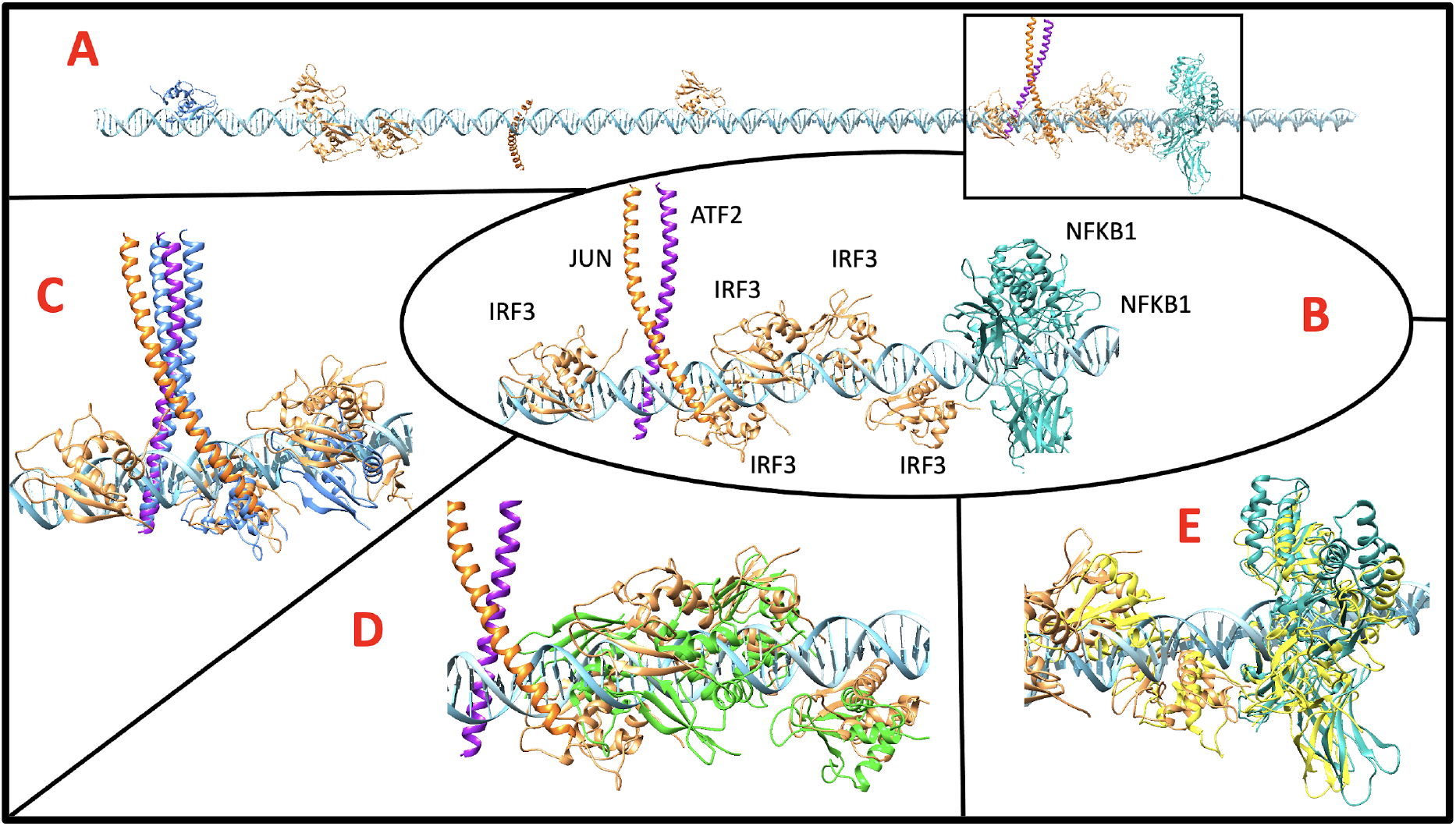
Model of the IFN-β enhanceosome complex. The structure of the complex formed by interactions between proteins and DNA is automatically built with the selected TFs and their binding sites in the enhancer sequence of IFN-β (see supplementary figure S4). A) Model of one of the complex structures obtained with the largest number of TFs while avoiding clashes between them. Due to the large time of computation, not all combinations of distinct conformations, produced by different templates, are tested. The structures of ATF-2 (purple), c-JUN (orange), IRF-3 (yeast) and NFKβ-1 (light blue) are shown on their binding with DNA (cyan), highlighting in a squared framework the region corresponding to the model of the enhanceosome proposed by Panne^21, 55^. B) Detail of the automated model obtained with ModCRE (IRF-3 is indicated as IRF3, ATF-2 as ATF2, c-JUN as JUN and NFKβ-1 subunit p105 as NFKB1): i) IRF-7 is missing; ii) NFKβ-1 forms a homodimer of two p105 subunits instead of the expected heterodimer with subunit p65 (RelA); and iii) an extra IRF-3 is bound at 5’ of the ATF-2/c-Jun binding. C) Detail of the superimposition of the model of ModCRE with the crystal structure of ATF-2/c-Jun and IRF-3 bound to the interferon-beta enhancer (code 1T2K^66^ from PDB, shown in blue). D) Detail of the superimposition of the model with the crystal structure of IRF-3 bound to the PRDIII-I regulatory element of the human IFN-β enhancer (code 2PI0^75^ of PDB, shown in green). E) Detail of the superimposition of the automated model with the crystal structure of NFKβ-1 (subunits p105 and p65), IRF-7, and IRF-3 bound to the IFN-β enhancer (code 2O61^55^ of PDB, shown in yellow).

### Example 2: TF-DNA interactions on top of the nucleosome

An interesting case of co-operation between TFs are the “pioneer factors” (the first to engage target sites in chromatin culminating in transcription by displacing nucleosomes^56^), or TFs that can bind on top of the nucleosome complex (as detected by NCAP–SELEX^57^). A structural view of this complex has shed light on the characteristics of the TF-DNA interaction and its effect upon the conformation of the nucleosome^22^. The server also has the possibility to produce a bent conformation such as the nucleosome that includes histones to form the complex (IMP is not applied because the structure of DNA is already defined). We have used the automated modelling of a nucleosome in complex with SOX2, SOX11 and OCT4 from the study of Dodonova et al. ^22^ to analyze these “pioneer factors”. Interestingly, we found two very significant binding regions of SOX2 and SOX11 (P-value <1.0e^-4^, around 58bp and 85bp positions) with the PWM predicted by ModCRE, but they were hampered by histones. We also found 3 sites with less significance (P-value < 1.0e^-3^) but accessible to SOX11/SOX2 and OCT4 (SOX11 and SOX2 share the same binding site preferences). However, SOX2 is missing in the predicted complex and the models of SOX11 and OCT4 clash with the DNA, implying the need of a posterior distortion to produce the complex (Figure 7). For OCT4 the model recreates the binding with both domains, but we must notice that with only one domain the binding is possible without producing clashes in the nucleosome second turn of DNA (in 98-112bps).

**Figure 7.**
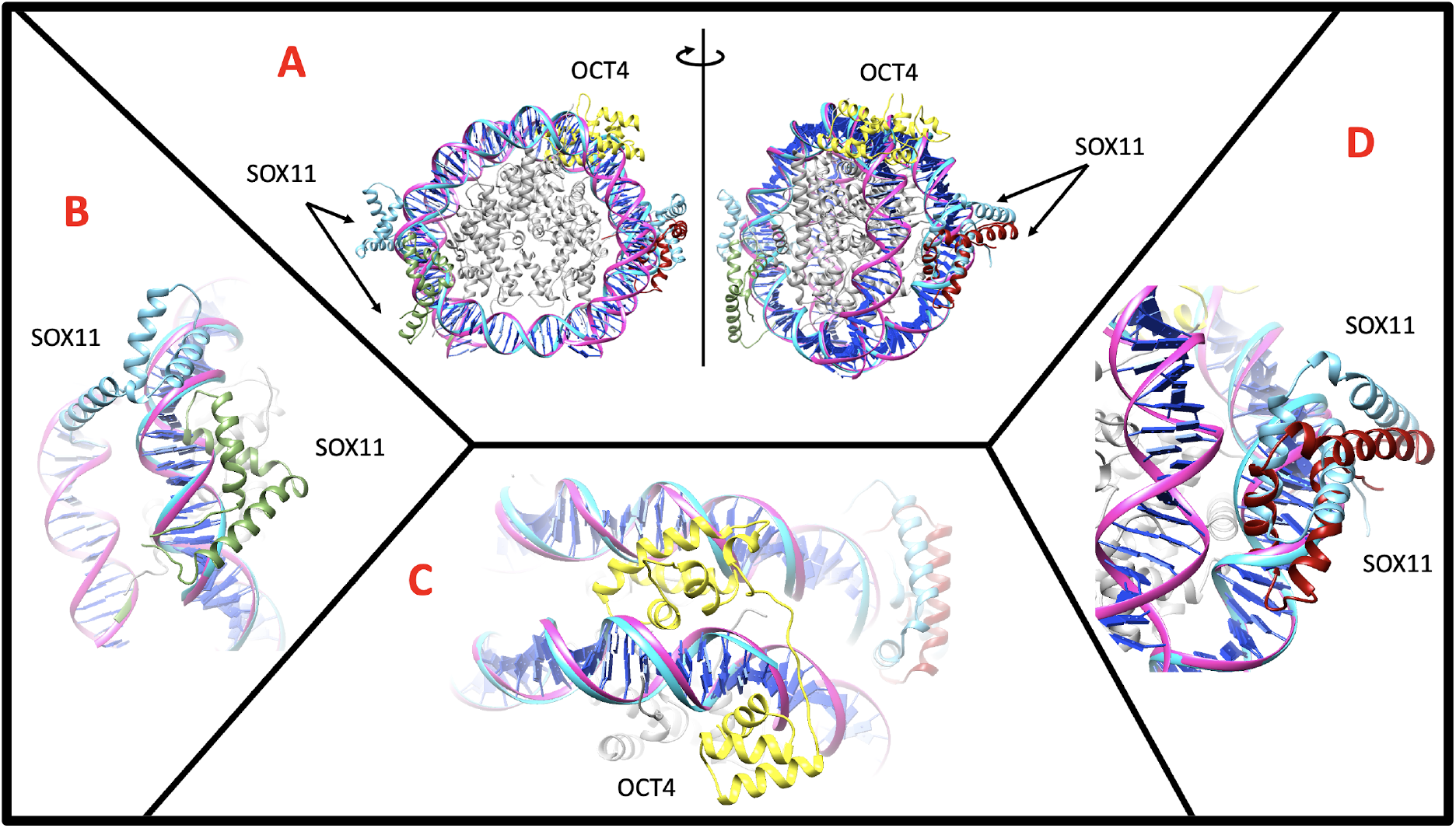
Model of the nucleosome complex with SOX11/SOX2 and OCT4. A) Front (left) and lateral (right) view of the complex obtained automatically with ModCRE superposed with the experimental structure (code 6T7C in PDB). Both model and experimental structures have only SOX11 bound (indicated as SOX11) to the nucleosome. The conformations modelled with ModCRE are shown in green (binding between 54 and 64 bps) and dark red (binding in the interval 86-96bps); the conformations of SOX11 from the experimental structure are shown in blue (binding in the interval of 50-60bps and 86-96 bps, respectively); and the model of OCT4 (indicated as OCT4) is shown in yellow (binding along an interval of 23-35bps). B) Detailed comparison of the binding of SOX11 between the automated model (green) and the experimental structure (blue). The predicted PWM fails to position neither SOX11 nor SOX2 around the 50bp (the closest fragment is in the interval of 54-57bps). C) Detail of the modelled binding of OCT4 (in yellow). Two domains of OCT4 produce the binding in a large interval of the DNA sequence (between 23 and 35bps), where the domain that binds in 3’ (around 28-35 bps) has also “non-binding” contacts (with many clashes) with the DNA fragment around 98-112bps. This suggests a potential weakening of the nucleosome complex conformation that could lead to unfasten it. D) Similar binding of SOX11 around 86-96bps of the model (dark red) and experimental (blue) structures. The DNA sequence used in the model is DNA1 from the study of Dodonova et al. ^22^ : ATCTACACGACGCTCTTCCGATCTAATTTATGTTTGTTAGCGTTATACTATTCT AATTCTTTGTTTCGGTGGTATTGTTTATTTTGTTCCTTTGTGCGTTCAGCTTAAT GCCTAACGACACTCGGAGATCGGAAGAGCACACGTGAT

## Discussion

Knowledge of TF-binding specificities is the foremost condition to understand gene regulation. Still, the binding preferences for many eukaryotic TFs are unknown or very complex *in vivo*^7, 58^. In this regard, computational tools can complement experimental methods. Several approaches have been taken in the recent years to computationally predict the binding of certain TF families such as the C2H2-Zf ^59^ or homeodomains^60^. Other approaches have also considered the use of statistical and molecular-mechanics potentials^61, 62^. In this work we have developed a structurebased approach to predict specific binding motifs of TFs, to identify cis-regulatory elements and to automatically model the structure of the transcription complex entailing the regulation. A similar structural-learning approach was recently developed and applied on homeodomain and C2H2-Zf families^63^ that predicted the PWMs by mapping experimentally known PWMs in the contacts of the interface of the TF-DNA structure, which in principle (but not suggested) it can be applied to any other TF by providing the structure of the complex with DNA. Our approach has been implemented in a server for the scientific community named ModCRE. The main limitation of these approaches is that they can only be applied to TFs for which the structure of the interaction with DNA is known. The scarcity of these experimental structures also affects the number of templates to be used by homology modeling. As shown in supplementary figure S5, ModCRE can be applied to most TF families defined in UniProt, and in combination with JASPAR database, our approach can be applied to 88.7% of the TF sequences (it is worth to remind that if the sequence similarity between the target and a TF in JASPAR is high, the PWM from this TF in JASPAR can be used to identify the binding preferences of the target). Besides, thanks to the recent advent of AlphaFold2 ^20, 64^ the structure of almost all TFs can be predicted, and remarkably the DNA binding motif, that often contains a large percentage of regular secondary structures, can be build.

The structure of any TF-DNA complex can be modelled either by docking or by superposition with other members of the TF family. As a tailored example we have studied the human motif of the CCAAT/enhancer-binding protein alpha (*C/EBPα.*). The human protein has not been crystallized, but the DNA binding motif of rat is 100% identical and the structure of the dimer is available in PDB with code 1NWQ ^65^. We downloaded the AlphaFold structure of *C/EBPα*. from human (AF-P49715-F1 from UniProt) and selected only the DNA binding domain (α-helix residues 284-344). We superposed this domain on each chain of the structure of the heterodimer of ATF-2 and c-Jun (PDB code 1T2K ^66^) to get the dimer complex of *C/EBPα*. with DNA. We used both structures to predict the PWM with ModCRE, one by submitting the sequence and getting the motif with 1NWQ as template, and the other by submitting the structure. This example shows almost identical motifs (see supplementary figure S6). In this line, the latest versión of RoseTTAFold, RoseTTAFoldNA ^54^, has incorporated the *ab initio* modelling of the structure of protein-DNA interactions that could be used straightforward in ModCRE to predict the PWM and scan one or more DNA sequences.

By incorporating the structural variability and flexibility of a TF we have designed an improvement of the prediction of its binding-sites based on the largest preference of motifs, each motif generated with one conformation. Thus, by scanning with several theoretical motifs of a TF, the majority of regions detected and predicted to bind will hit around the right location of the binding site. Using a collection of motifs derived from different models of a TF is in consonance with the idea that TFs can interact with the DNA adopting different conformations^7^. Not only the dynamics of the protein but also the flexibility of the conformation of the DNA plays a relevant role in the identification of the binding site. Molecular dynamics of such complexes have recently been used to predict the binding affinity of TFs and to predict its corresponding PWMs^67^. We used homology modelling in the same line in our approach. Homology modelling is very convenient when several templates are available because it generates a collection of models of a TF without requiring large computational resources. Still, the time of computation to calculate the PWM in the server is between 30’ and 5 hours (depending on the size of the interface). Similarly, AlphaFold can be applied to obtain several conformations. In agreement with this, ModCRE’s modeling pipeline is a valuable resource to study the conformations of large regulatory complexes. Structural models of TF-DNA interactions provide fundamental information to understand TF function and behavior. Our pipeline models complexes of TF-DNA interactions involving DNA bindings and proteinprotein interactions between TFs and transcription co-factors. We hypothesize that the correct binding site is among the selection of sites where most conformations of TFs accumulate when considering the cooperation with other TFs and co-factors. A drawback for the server is that the final steps to model the macro-complex require a large time of computation. Nevertheless, this may be a promising strategy helping to overcome the number of false positives found when scanning a DNA sequence with a single PWM^7, 58^ or at least to narrow the predictions and simultaneously comprehend the cooperativity between transcription factors and co-factors.

## Methods

### Software

We use DSSP (version CMBI 2006)^68^ to obtain the secondary structure and surface accessibility; X3DNA (version 2.0)^69^ to obtain DNA structures; *matcher* and *needle*, from the EMBOSS package (version 6.5.0)^70^, to obtain local and global alignments, respectively; BLAST (version 2.2.22)^49^ to obtain potential homologs of a protein sequence; MODELLER (version 9.9)^43^ to model the structure of a protein by homology; CD-HIT to obtain a non-redundant set of sequences of TFs^71^ and the programs FIMO^48^ and TOMTOM of the MEME suite^44^ to obtain the fragments of a DNA sequence that aligns with a Position-Weight Matrix (PWM) ^72^ and to compare two PWMs, respectively.

### Databases

Atomic coordinates of protein complexes are retrieved from the PDB repository^73^ and protein codes and sequences are extracted from UniProt^51^. We only selected the structures of PDB corresponding to TF-DNA interactions. Binding information of TFs was obtained from protein binding microarrays (PBMs) experiments in the Cis-BP database (version 2.00) ^39, 40^. PBMs experiments indicate the binding affinity between TFs and DNA 8-mers with the E-score value (between −0.50 and 0.50). DNA 8-mers of a TF with E-scores above 0.45 correspond to high affinity interactions (also named positive), while DNA 8-mers with E-scores below 0.37 are considered non-bound (or negative); the rest of E-scores are discarded for statistic analyses.

### Interface of protein-DNA structures

We defined the contacts between TF and DNA using three residues: one amino acid and two contiguous nucleotides of the same strand. The distance of a contact is the distance between the Cβ atom of the amino acid residue and the average position of the atoms of the nitrogen-bases of the two nucleotides and their complementary pairs in the opposite strand^28^. Additional features are considered for a contact, such as the secondary structure and solvent accessibility of the amino-acid or the DNA closest groove (major or minor) of the two nucleotides.

### Statistical potentials

We used the definition of statistical potentials described by Feliu et al.^74^ and Fornes et al.^28^ applied on the selected database of structures of TF-DNA interactions. These were calculated with the distribution of contacts at less than 30 Å, using an interval criterion or a distance threshold. We used positive DNA 8-mers from PBMs to extend the number of contacts. Briefly, we modelled the interactions of the DNA 8-mers with the TFs using the available structures of TF-DNA pairs in PDB or those of their closest homologs (see details in supplementary material). We transformed the statistical potentials into Z-scores to identify the contacts with best scores (best distance and best contact residues: i.e. the amino acid and the two nucleotides). To avoid redundancies in the statistical potentials we used a criterion of around 70-80% identical contacts for family-specific potentials (i.e. calculated with members of the same TF family sharing the same fold structure) and 40-50% for a general scenario.

### Structural modeling of TF-DNA complexes

Several structural models of a TF-DNA complex were obtained using all its available templates from PDB. First we used BLAST to find the homologs with known structure (template), then the sequence of the query was aligned with the sequences of the templates using MATCHER from the EMBOSS package^70^ and a model was built with each template using MODELLER^43^. The modeling of the DNA was obtained with the X3DNA package^69^ preserving the DNA conformation from the template. This approach required that all the templates used for TF-DNA modeling contained both a TF and a double stranded DNA molecule.

### Construction of PWMs using TF-DNA structural models

We used the Z-scores of statistical potentials to obtain the PWM. We selected for each TF family the features optimizing the PWM prediction (see further) and we used the Z-score of *ES3DC_dd_ (ZES3DC_dd_*) as defined in Meseguer et al. ^37^. First, we obtained several models of a TF-DNA interaction using all possible templates. Second, for each model we scored with *ZES3DC_dd_* all the potential DNA sequences of the binding site (i.e. 4^N^ sequences, with N the size of the binding site, or an alternative heuristic approach as explained in the supplementary). Third, we normalized the scores between 0 and 1 and we ranked the DNA sequences. Finally, for each model we selected the sequences with the top scores (this cut-off threshold was also optimized, taking values between 0.7 and 1 in intervals of 0.01). We used the alignment of these sequences to calculate a predicted PWM for each model.

### Optimization of parameters to predict PWMs by grid search

The parameters to predict PWMs that needed to be optimized for each TF family are: 1) the definition of distances’ distribution used to calculate statistical potentials: either by interval-bins (i.e. *x* −1 < *d* ≤ *x*) or a threshold (i.e. *d* ≤ *x*); 2) the use of a theoretical approach to complete the space of contacts (i.e. using a Taylor’s polynomial approach, see supplementary); 3) the dataset of structures used to calculate the potentials: using only the contacts from structures of PDB or adding those from experiments of PBMs; 4) using a general statistical potential calculated with all known TF-DNA structures or a specific potential calculated with the structures of the same family and fold; 5) the maximum distance to include the contacts of an interface (testing distances at 15 Å, 22 Å and 30 Å); and 6) the cut-off threshold to select top ranked DNA sequences used to calculate the PWM (see above). The function to be optimized was the accuracy to predict the PWM of each TF family (i.e. maximum accuracy). A predicted PWM was successful if the alignment with the experimental PWM taken from Cis-BP database was significant (this was calculated with TOMTOM). Then, we selected the parameters that maximized the accuracy of the TF family with the following conditions: 1) maximum number of significant good predictions according to TOMTOM score; 2) best TOMTOM scores when a similar number of significant solutions were achieved; and 3) the lowest value of the threshold, when several similar solutions were obtained.

## Supporting information

Supplementary Theory

## Acknowledgments

The work was supported by grants PID2020-113203RB-I00 and “Unidad de Excelencia María de Maeztu” (ref: CEX2018-000792-M), funded by the MCIN and the AEI (DOI: 10.13039/501100011033). OG acknowledge support from PGC2018-095745-B-I00 and PID2021 −127773NB-I00/MICIN/AEI/10.13039/501100011033/ FEDER, EU (MICIN, AEI and FEDER), and support from RGP0017/2020 (HFSP).

