## Supplementary Tables and Figures for "Structure-based learning to model complex protein-DNA interactions and transcription-factor co-operativity in *cis*-regulatory elements"

### *Supplementary Figures*

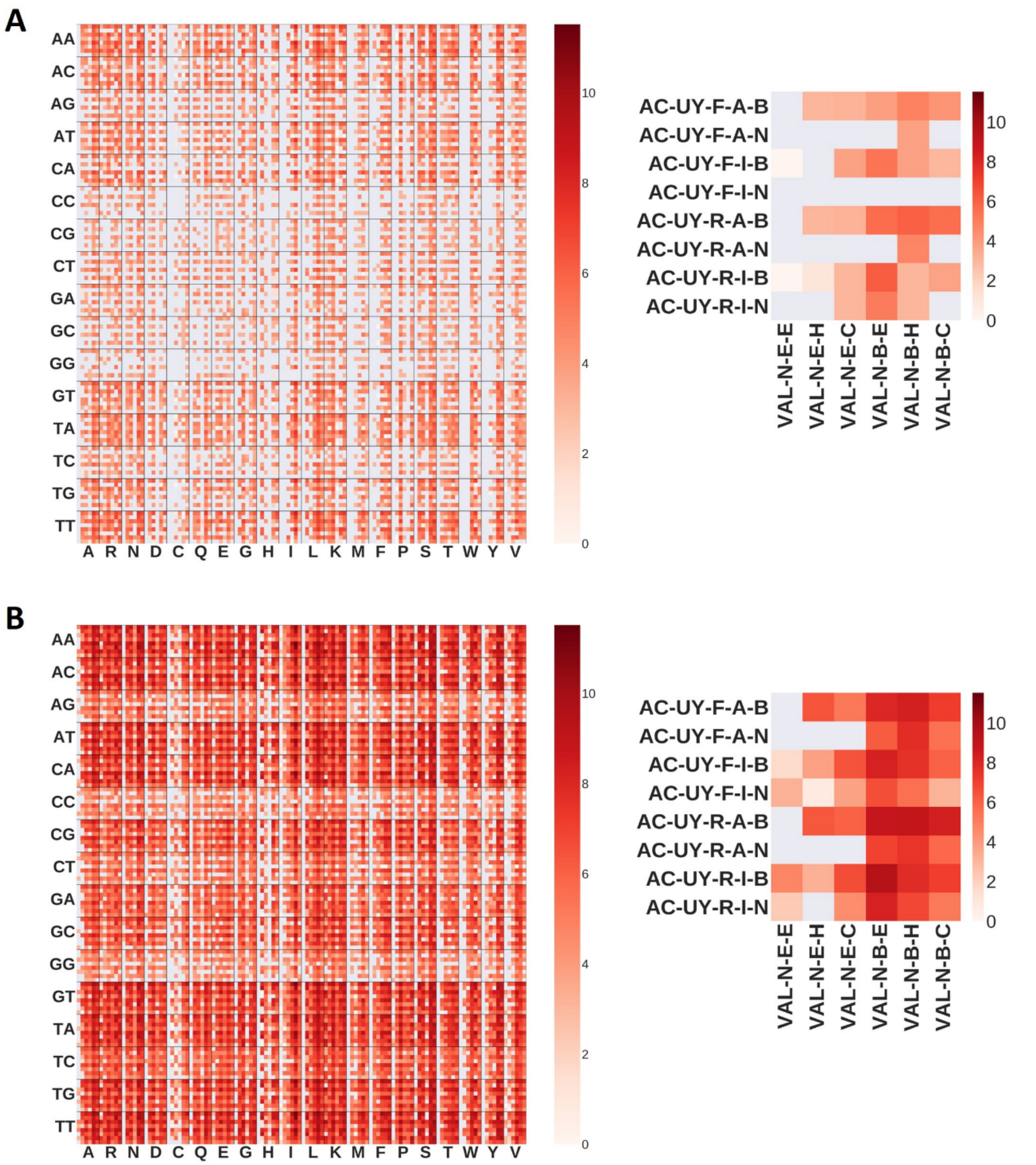


*Figure S1. Examples of heatmap plots showing the number of amino acid—dinucleotide contacts at distance shorter than 30Å in a logarithmic scale. (A) Example of amino-acid and dinucleotide contacts extracted from PDB structures of the Forkhead family. (B) Example of amino-acid and dinucleotide contacts obtained from PDB structures and PBMs experiments of transcription factors of the Forkhead family. Detailed view of a cell in each heatmap is shown in the right side. Each square inside the cell shows the logarithm of the number of contacts with specific features. The example shows the contacts between a valine amino-acid (VAL) and a dinucleotide of adenosine-cytosine (AC), with their specific environment features.*

*Amino-acid environment features are: hydrophobicity (P as polar, N non polar), surface accessibility (E if exposed, B if buried) and secondary structure (E for β-strand, H for helix and C for coil). Dinucleotide environment featuress are: type of nitrogenous bases (U for purine, I for pyrimidine), closest DNA strand (F for forward, R for reverse), closest DNA groove (A for major, I for minor) and closest chemical group (B if phospho-ribose backbone atoms, N if nucleobase). All heatmaps can be downloaded from the web in http://aleph.upf.edu/modcre/##faq.*

*A)*

| | General | | | | Family | | | |
|---|---|---|---|---|---|---|---|---|
| | PDB | | PDB + PBM | | PDB | | PDB + PBM | |
| | - | Taylor | - | Taylor | - | Taylor | - | Taylor |
| **AFT** | 0.0057 | 0.0067 | 0.051 | 0.29 | 0.013 | 0.34 | 0.92 | 0.56 |
| **APSES** | 0.16 | 0.48 | 0.73 | 0.78 | 0.69 | 0.78 | 0.9 | 0.71 |
| **ARID/BRIGHT** | 0.66 | 0.72 | 0.84 | 0.84 | 0.74 | 0.63 | 0.94 | 0.84 |
| **B3** | 0.029 | 0.016 | 0.013 | 0.03 | 0.0076 | 0.014 | 0.92 | 0.93 |
| **C2HC ZF** | 0.53 | 0.58 | 0.36 | 0.35 | 0.73 | 0.91 | 0.98 | 0.96 |
| **CSL** | 0.011 | 0.011 | 0.053 | 0.053 | 0.013 | 0.018 | 0.066 | 0.063 |
| **CxxC** | 0.15 | 0.2 | 0.52 | 0.55 | 0.22 | 0.045 | 0.98 | 0.59 |
| **GCM** | 0.052 | 0.33 | 0.4 | 0.7 | 0.45 | 0.71 | 0.87 | 0.93 |
| **HSF** | 0.044 | 0.036 | 0.2 | 0.17 | 0.076 | 0.13 | 0.72 | 0.69 |
| **MADS box** | 0.092 | 0.078 | 0.33 | 0.41 | 0.7 | 0.47 | 0.99 | 0.97 |
| **MBD** | 0.019 | 0.012 | 0.028 | 0.046 | 0.0072 | 0.11 | 0.57 | 0.65 |
| **Ndt80/PhoG** | 0.033 | 0.014 | 0.85 | 0.76 | 0.0081 | 0.14 | 0.005 | 0.95 |
| **POU** | 0.025 | 0.023 | 0.56 | 0.62 | 0.13 | 0.096 | 0.68 | 0.15 |
| **Paired box** | 0.023 | 0.03 | 0.22 | 0.18 | 0.065 | 0.075 | 0.39 | 0.33 |
| **Rap1** | 0.0091 | 0.0093 | 0.0072 | 0.0071 | 0.0095 | 0.012 | 0.013 | 0.011 |
| **Rel** | 0.17 | 0.17 | 0.15 | 0.14 | 0.11 | 0.18 | 0.55 | 0.52 |
| **Runt** | 0.029 | 0.034 | 0.36 | 0.21 | 0.068 | 0.4 | 0.89 | 0.89 |
| **SMAD** | 0.089 | 0.043 | 0.21 | 0.25 | 0.077 | 0.12 | 0.45 | 0.45 |
| **TCR/CxC** | 0.011 | 0.023 | 0.7 | 0.9 | 0.15 | 0.14 | 0.96 | 0.77 |
| **TEA** | 0.032 | 0.03 | 0.039 | 0.056 | 0.2 | 0.38 | 0.83 | 0.95 |

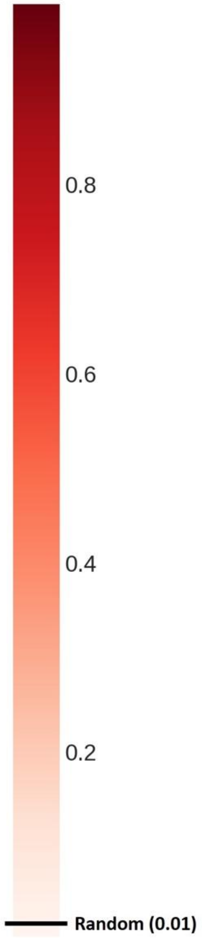

*B)*

| | General | | | | Family | | | |
|---|---|---|---|---|---|---|---|---|
| | PDB | | PDB + PBM | | PDB | | PDB + PBM | |
| | - | Taylor | - | Taylor | - | Taylor | - | Taylor |
| AFT | 0.2 | 0.32 | 0.83 | 0.87 | 0.61 | 0.95 | 1 | 0.98 |
| AP2 | 0.35 | 0.84 | 0.9 | 0.9 | 0.76 | 0.86 | 0.9 | 0.88 |
| APSES | 0.89 | 0.97 | 0.98 | 0.99 | 0.99 | 1 | 1 | 0.99 |
| ARID_BRIGHT | 0.98 | 0.98 | 0.99 | 0.99 | 0.99 | 0.98 | 1 | 0.99 |
| B3 | 0.81 | 0.68 | 0.62 | 0.8 | 0.39 | 0.58 | 1 | 1 |
| C2H2_ZF | 0.67 | 0.67 | 0.71 | 0.7 | 0.67 | 0.71 | 0.81 | 0.81 |
| C2HC_ZF | 0.98 | 0.98 | 0.98 | 0.98 | 0.99 | 1 | 1 | 1 |
| CSL | 0.52 | 0.52 | 0.57 | 0.57 | 0.55 | 0.56 | 0.58 | 0.58 |
| CxxC | 0.78 | 0.79 | 0.91 | 0.91 | 0.9 | 0.78 | 1 | 0.99 |
| DM | 0.72 | 0.7 | 0.72 | 0.7 | 0.81 | 0.88 | 0.9 | 0.9 |
| E2F | 0.64 | 0.75 | 0.94 | 0.97 | 0.94 | 0.98 | 1 | 1 |
| Ets | 0.81 | 0.82 | 0.88 | 0.88 | 0.88 | 0.89 | 0.9 | 0.9 |
| Forkhead | 0.74 | 0.74 | 0.83 | 0.83 | 0.81 | 0.81 | 0.89 | 0.89 |
| GATA | 0.64 | 0.67 | 0.89 | 0.89 | 0.87 | 0.87 | 0.93 | 0.92 |
| GCM | 0.84 | 0.95 | 0.97 | 0.99 | 0.97 | 0.99 | 1 | 1 |
| HSF | 0.73 | 0.68 | 0.85 | 0.83 | 0.66 | 0.85 | 0.93 | 0.94 |
| Homeodomain | 0.81 | 0.81 | 0.86 | 0.86 | 0.83 | 0.8 | 0.79 | 0.73 |
| IRF | 0.61 | 0.65 | 0.86 | 0.86 | 0.71 | 0.66 | 0.94 | 0.93 |
| MADS_box | 0.87 | 0.87 | 0.97 | 0.97 | 0.98 | 0.98 | 1 | 1 |
| MBD | 0.56 | 0.51 | 0.59 | 0.62 | 0.35 | 0.66 | 0.98 | 1 |
| Myb_SANT | 0.71 | 0.74 | 0.91 | 0.9 | 0.82 | 0.87 | 0.99 | 0.99 |
| NAC_NAM | 0.55 | 0.54 | 0.52 | 0.56 | 0.61 | 0.56 | 0.87 | 0.85 |
| Ndt80_PhoG | 0.85 | 0.67 | 1 | 1 | 0.45 | 0.9 | 0.00039 | 1 |
| Nuclear_receptor | 0.73 | 0.72 | 0.85 | 0.84 | 0.8 | 0.8 | 0.9 | 0.9 |
| POU | 0.73 | 0.71 | 0.98 | 0.98 | 0.79 | 0.76 | 0.97 | 0.83 |
| Paired_box | 0.67 | 0.73 | 0.94 | 0.92 | 0.72 | 0.76 | 0.95 | 0.96 |
| Rap1 | 0.44 | 0.45 | 0.31 | 0.31 | 0.45 | 0.48 | 0.52 | 0.51 |
| Rel | 0.79 | 0.78 | 0.78 | 0.78 | 0.81 | 0.82 | 0.84 | 0.81 |
| Runt | 0.78 | 0.79 | 0.98 | 0.95 | 0.88 | 0.97 | 0.95 | 1 |
| SMAD | 0.82 | 0.79 | 0.82 | 0.86 | 0.86 | 0.85 | 0.93 | 0.91 |
| Sox | 0.93 | 0.93 | 0.95 | 0.95 | 0.93 | 0.94 | 0.97 | 0.97 |
| T-box | 0.61 | 0.62 | 0.89 | 0.89 | 0.65 | 0.85 | 0.95 | 0.93 |
| TCR_CxC | 0.57 | 0.72 | 0.99 | 1 | 0.87 | 0.91 | 1 | 1 |
| TEA | 0.82 | 0.82 | 0.77 | 0.79 | 0.89 | 0.91 | 0.99 | 1 |
| WRKY | 0.93 | 0.96 | 0.8 | 0.97 | 0.74 | 0.97 | 0.97 | 0.97 |
| Zinc_cluster | 0.64 | 0.66 | 0.77 | 0.78 | 0.62 | 0.77 | 0.84 | 0.85 |
| bHLH | 0.68 | 0.67 | 0.75 | 0.75 | 0.74 | 0.75 | 0.82 | 0.79 |
| bZIP | 0.67 | 0.65 | 0.69 | 0.69 | 0.68 | 0.7 | 0.74 | 0.71 |

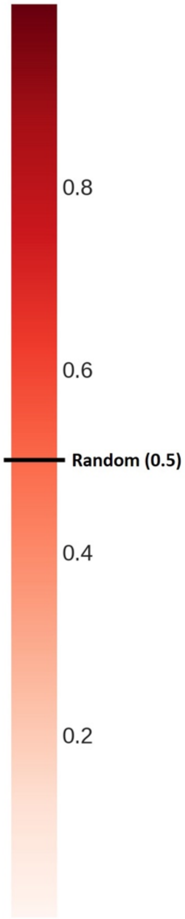


*Figure S2*
*A) Area under the curve of precision-recall (AUPRC) on the prediction of positive and negative 8-mers of the PBM experiments. We analyzed TFs of all families from Cis-BP database with PBMs experiments. We used the PWMs predicted with structural models of TFs using different features to calculate the $ZES3DC_{dd}$ statistical potential (such as the use of the Taylor polynomial approach, using contacts extracted from PDB or from PDB plus those derived from PBMs experiments or using only contacts obtained with TFs of the same family). Not all negative 8-mers were used, forcing the ratio of positive/negatives to be 1 to 100 (negative 8-mers were selected randomly). The table shows the study of families with less than 10 different TFs supporting the results. B) Area under the ROC curve (AUROC) on the prediction of positive and negative 8-mers of the PBM experiments. The table shows the study for all families of TFs.*

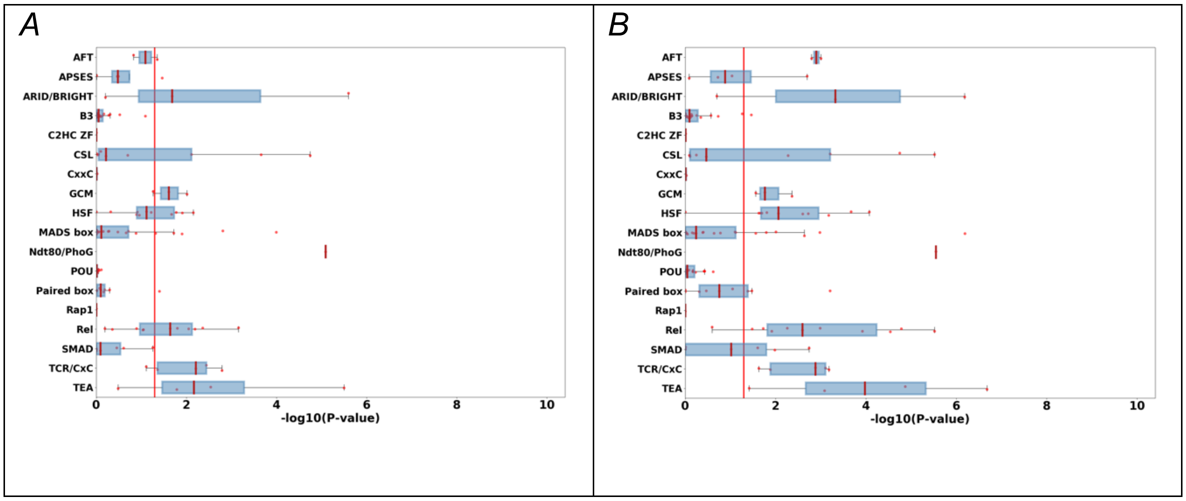


*Figure S3*
*Distribution of “similarity scores” to compare predicted and experimental PWMs of TFs. Legends are as in Figure 3. Plots correspond to TF families with less than 10 different TF sequences.*

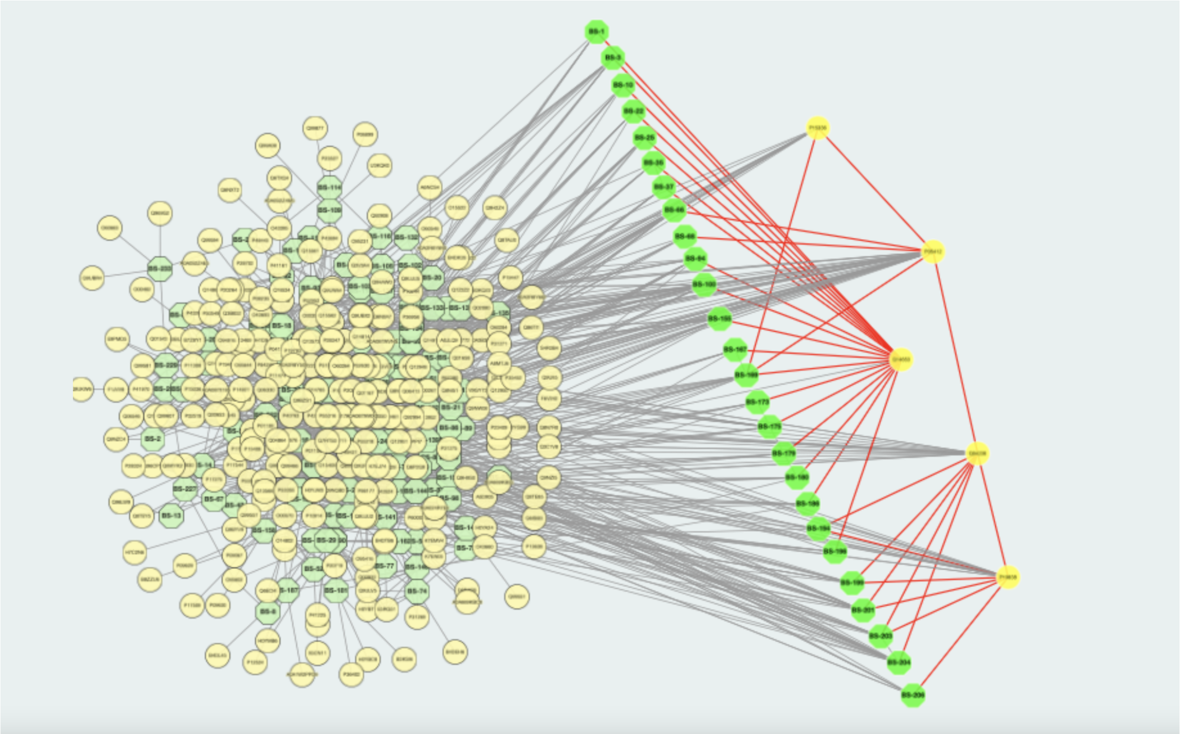

*Figure S4*

*Cis-regulatory elements of the IFN-β enhancer site and its potential transcription factors. Selected bindings of c-Jun (P05412), ATF-2(P15336), IRF-3(Q14653), IRF-7(Q92985), NFKβ-1 subunit p105 (P19838) and RelA(Q04206) human transcription factors on the enhancer region of the IFN-β (formed by 250 bps). Uniprot codes of these proteins are shown within parenthesis. Bindings are selected with a significant match (P-value < $5.0e^{-4}$) of the PWM predicted with the structure using ModCRE and achieving the top scores (i.e. normalized $ZES3DC_{dd}$ values, see supplementary material). Under these conditions IRF-7 is not selected and a molecule of IRF-3 occupies its location. Transcription factors are shown in yellow and binding sites in green. Interactions between TFs and between TFs and binding sites are shown as edges. Edges between the selected TFs and their binding sites are shown in red. The rest of interactions (in black) correspond to interactions of these TFs with other human proteins (also TFs). The sequence of the enhancer region of IFN-β is:*

*GTTTGCTTTCCTTTGCTTTCTCCCAAGTCTTGTTTTACAATTTGCTTTAGTCATT
CACTGAAACTTTAAAAAACATTAGAAAACCTCACAGTTTGTAAATCTTTTTCCCT
ATTATATATATCATAAGATAGGAGCTTAAATAAAGAGTTTTAGAAACTACTAAAA
TGTAAATGACATAGGAAAACTGAAAGGGAGAAGTGAAAGTGGGAAATTCCTCT
GAATAGAGAGAGGACCATCTCATATAAATAGG.*

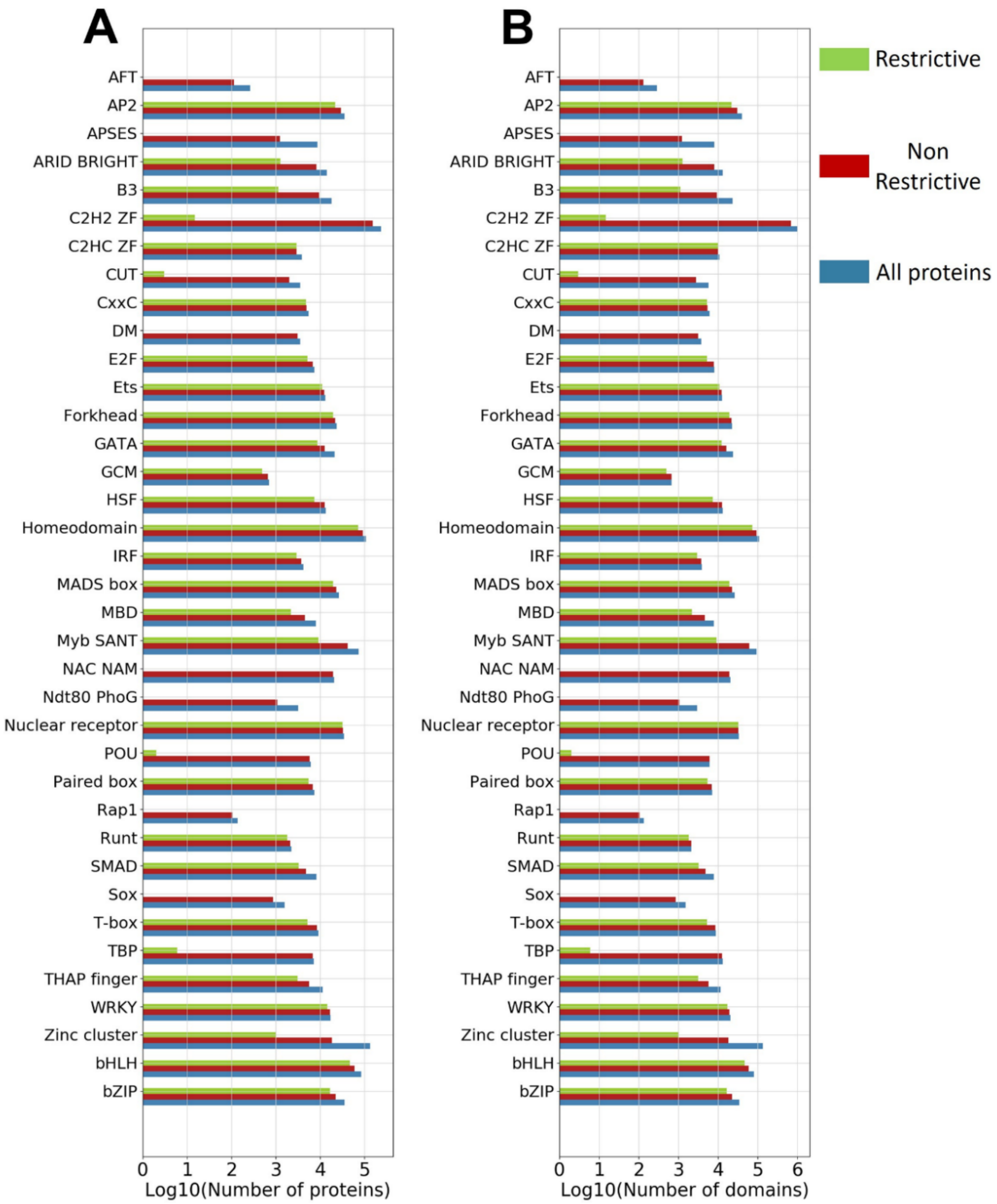


*Figure S5*

*Applicability of the structure-based approach to predict TF motifs using homology modelling. The plots show the total number of proteins (A) and domains (B) for which a PWM can be predicted using homology modelling (applying the server of*

*ModCRE). TF families are indicated in the vertical axis versus the logarithm of the number of proteins (or domains), showing: i) the total number of UniProt sequences that match the PFAM model of the family (blue); ii) those that can be modelled without gaps in the interface of the interaction with DNA (green); and iii) and those with gaps in the interface (red).*

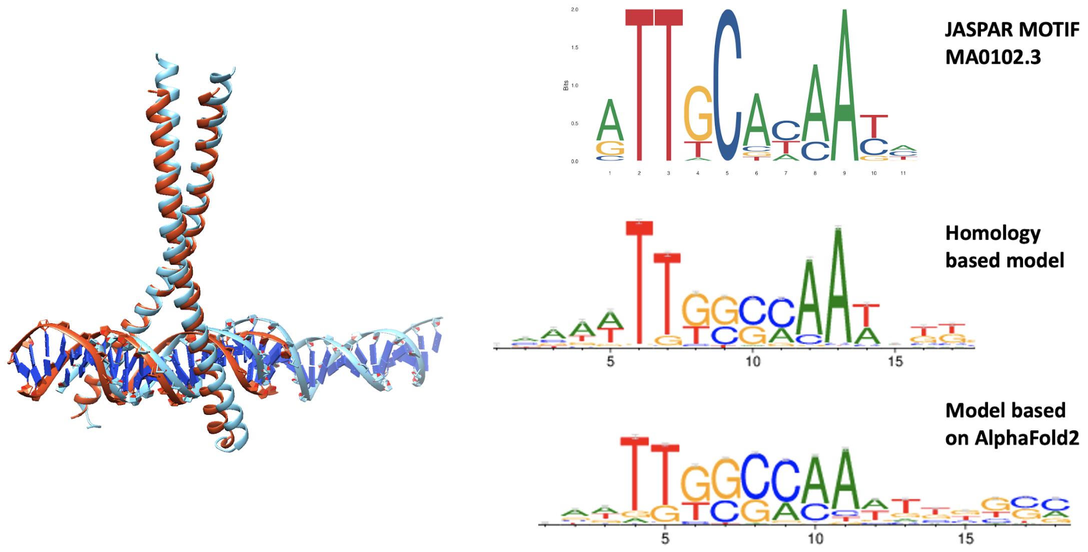


*Figure S6*
*Prediction of PWMs of human C/EBPα. The ribbon plate structures on the left side show the superposition of a model of human C/EBPα (red) obtained with the crystal structure of rat (code 1NWQ in PDB) as template and a model based on the structure predicted with AlphaFold2 (blue). The right side shows a comparison of the motif logos of human C/EBPα: i) the top logo corresponds to motif MA0102.3 from JASPAR; ii) the middle logo is obtained with the prediction based on the homology model of human C/EBPα; and iii) the bottom logo is obtained with the structure modelled upon the structure prediction of human C/EBPα obtained with AlphaFold2.*

*Figures S7*
*Compressed folder with the distribution of the normalized ranking score of motif predictions with the nearest-neighbor (state-of-the-art) approach and the structure-based approach (ModCRE) using rank and rank-enrichment. Legends are as for figures 4 and 5.*

***Legends of supplementary Tables***

*Table S1*
*Logos of experimental PWMs (from JASPAR and CisBP) and of their best predictions. The table shows the logos of one TF per family, selected from Table S2, with a good match with at least one of the experimental PWMs. We only used TFs for which the PWMs were also obtained by PBMs in CisBP as in Table 1. Labels are as in Table 1.*

*Table S2*
*Summary of the analysis of predicted PWMs of TFs with experimental motifs in JASPAR and CisBP (obtained with PBMs). The ID code of the TF as defined in CisBP is shown in the first column (TFID). Second and third columns show the name of the family (also as coded in CisBP). Fourth and fifth columns show the UniProt code of the TF and the encoded name of the model used to generate the PWM most similar to the motif in JASPAR. The code of the model identifies the TF, the first and last amino acids of the modelled sequence, the PDB code of the template and the chain, and a number to identify the model out of 100. Sixth and seventh columns show the codes of the motifs in JASPAR and CisBP datasets, respectively. Eight and nineth columns show the ratio of models yielding PWMs that significantly align with the experimental motif of JASPAR and CisBP, respectively. Successes are identified in the last four columns with a Boolean (1 if right, 0 if wrong). In the first two columns of the last four we define the success by the ratio (if more than 50% of predicted PWMs were significantly aligned with the experimental motif) for the JASPAR and CisBP motifs, respectively. In the last two columns we define the success if at least one of the predicted PWMs aligned significantly with the experimental motif from JASPAR and CisBP, respectively.*

*Table S3*
*Number of TFs and predictions used in Figures 4 and 5 of the main manuscript. Each column shows for each TF family and for the sum of all families (global) the total number of TF (tfs) and their predictions (predictions) in bins 15 to 95 defined as in Figure 4, using the nearest-neighbor approach (nn) and our method (modcre). The table is split in two pages, one for the results of the normalized ranking in figure 4 (NORMAL_RANK), and the other for the normalized rank of the enrichment (NORMAL_RANKENRICHED). It must be noted that the number of TFs is equal to the number of predictions in the rank enrichment, because a single prediction is selected for a TF. Besides, the prediction with ModCRE is performed for each single TF (2283 TFs in total), while for the nearest neighbor approach we used the domains/variants of TFs as selected from CisBP, where each domain corresponds to a single motif (2638 in total).*

*Table S4*

*Tables for all TF families showing the average error when ranking predictions as a function of the accepted similarity between two PWMs (predicted versus experimental). The table "global.error.csv" shows the averaged error of rankings using all TFs. The error of the rank is calculated as:*

$$error = 100 \times (\langle N_{threshold} \rangle - 1)/M$$

*where: M is 2683, i.e. the total number of motifs to be ranked;* $\langle N_{threshold} \rangle$ *is the average of the number of neighbors of a motif that are significantly similar under an accepted criterion of similarity. The neighbors of a motif have a PWM comparable to the experimental motif with SIM <* $10^{-threshold}$*, being SIM the criterion of similarity between PWMs that is obtained with TOMTOM: P-value, E-value or Q-value. The column "threshold" shows the threshold used to calculate the number of neighbors. Columns showing the error are entitled as "error_rank_EV_mean" (for E-value criterion), "error_rank_pV_mean" (for P-value criterion) and "error_rank_qV_mean" (for q-value criterion).*
