## Supplementary Theory for "Structure-based learning to model complex protein-DNA interactions and transcription-factor co-operativity in *cis*-regulatory elements"

**METHODS**

**Index**


1. Software requirements
2. Databases
3. Interface and triads of protein-DNA
4. Statistical potentials
5. Solving scarcity of data by using Taylor's polynomial series approach
6. Z-scores
7. General and family-specific potentials
8. Structural modeling of TFs complexed with DNA
    a. Protein monomer/dimer modeling
    b. Protein homo/hetero dimer modeling
    c. DNA modeling
    d. Modeling protein-protein interactions.
    e. Modeling C2H2 structures for B1H
9. Use of experimental TF-DNA binding to calculate statistical potentials
10. Prediction of the PWM using the sequence of a TF
    a. Straight-forward prediction
    b. Prediction by enrichment
11. Scoring TF-DNA binding with structure
12. Construction of PWMs using TF structures
13. Predicting the PWM with the structure of a TF
    a. Straight-forward prediction
    b. Prediction by enrichment
    c. Comparison of different sets and methods to predict PWMs
14. Optimal conditions to predict PWMs using the structure of a TF (grid search)
15. Scanning of binding sites and TF clusters along a DNA sequence.
    a. Scanning of DNA binding domains
    b. Score per Nucleotide: profiles of a DNA binding site.
    c. TF profiles along a DNA fragment.
    d. Prediction of TFs that bind a DNA fragment
    e. Clusters of TFs and complexes of regulatory elements.
16. Modeling a selected TF-complex of a cluster of close TFs in a DNA fragment


---

### 1. Software requirements

We require the following software: DSSP (version CMBI 2006) (1) provides protein structural features; X3DNA (version 2.0) (2) is used to analyze and generate DNA structures; *matcher* and *needle*, from the EMBOSS package (version 6.5.0) (3), produces local and global alignments, respectively; BLAST (version 2.2.22) (4) and MMseqs2 (5) are employed to search homologs of a target protein; MODELLER (version 9.9) (6) is used to create structural models with all the templates similar to our target; CE-align algorithm (7), as implemented in PyMOL (version 1.5) (8), is used for structural superimpositions of DNA to merge complexes and TMalign (9) to superimpose similar

TF folds; and the programs FIMO and TOMTOM from the MEME suite(10) are used to scan a DNA sequence with a Position-Weight Matrix and to compare two PWMs, respectively.

### 2. Databases

Structural information is retrieved from the PDB repository (11) and protein codes and sequences are extracted from UniProt (January 2022 release) (12). We select all transcription factors as defined in CIS-BP database (version 2.0) (13) to generate the internal database of structures. We distinguish **monomer** and **homodimer** TF structures, accepting that some TF families often act as homodimers ("B3", "bHLH", "bZIP", "Leafy", "MADS box", "Rel", "Nuclear Receptor", "STAT", and "Zinc cluster"). We extend the set with all other structures that interact, with more than 5 contacts, with a double strand DNA molecule (see further for the definition of contacts in section 3) and name the set **$PDB_{DNA}$**. We rearrange the set of structures by separating them in chains and constructing a set of structures formed by single protein-chains interacting with a double-strand helix (**single-chain $PDB_{DNA}$**).

We use the program TMalign to compare structures of proteins of the single-chain $PDB_{DNA}$ set by superimposing them. We group each structure with all others matching a *good* superimposition in a set named "**folds**". For example, code 1PUF_A that identifies protein-chain A of 1PUF in PDB `produces the group named "1PUF_A folds". The superposition is considered *good* if it produces a TM-score higher than 0.5. We say that all TF-DNA structures belonging in the same group of folds have the same fold.

TF-DNA binding information is retrieved from the CIS-BP database (version 2.0). We retrieve for each TF in the database the name (Gene Name, Accession Number, Ensembl, and/or any other code to associate with the protein sequence), the specie, the family group and the Position Weight Matrix (PWM). We also retrieve for each TF the list of 8–mers evaluated in the PBM experiment with the corresponding E–score values. The name and species of the TFs are used to obtain the Entry code from UniProt and the corresponding protein sequence. We use the E–scores of the 8–mers to classify them in positive bindings (E-score > 0.45) and negative bindings (E-score < 0.37). We use the PWM of the TF assigned in CIS-BP to select the correct orientation of the 8-mer strand (i.e. we only use the 8-mer sequence of the strand corresponding to the best match with the PWM). We select the 8-mer with the best score out of all 8-mers of a TF as the best bound DNA sequence. Then, we use the best bound sequence to align all other 8-mers considered positive using the program *needle* of EMBOSS package (3). All 8-mers in the positive binding set aligning with one or more gaps with the best bound sequence are removed from the set. The trimming of the set of positives allows us to have a set of 8-mers, all of them with the correct strand orientation and without gaps that can be used to generate a PWM based on the 8-mers of the PBM.

Binding information of Zinc-finger family C2H2-ZF is retrieved from bacteria one-hybrid (B1H) experiments (14). The experiment distinguishes between Zinc-finger individual domains at the C-tail (F3 domain) and inner domain (F2 domain). The experiment performs the screening of all 64 possible 3bp targets for interactions with C2H2-ZF domains from multiple large protein libraries based on Zif268 structure with six variable amino acid positions on each individual domains F2 and F3 (15).

### 3. Interface and triads of protein-DNA structures

The interface between a protein (e.g. transcription factor) and DNA is defined by the residues (amino-acids and nucleotides) in contact. A general approach for protein-protein interactions is to consider that two residues are in contact if the distance between a pair of atoms from each residue is shorter than 5Å. Usually, a shell around the interface is defined by residues in contact at distance 12 Å. One interface is larger than other if the number of pairs of residues in contact is larger. For TFs in the same fold (i.e. as defined in section 2), we compare the similarity between interfaces of two protein-DNA structures by checking the number of common pairs of residues in both interfaces. The **score of similar-interface** between interfaces is calculated as the percentage of common pairs of residues in both interfaces with respect to the smallest interface (with independence of their residue-number and nucleotide position in their respective structures, but also independent of the distance as long as they form part of the interface).

We define ***triads*** as a type of contacts between the protein and the double-strand DNA helix. Triads are formed by three residues: one amino-acid and two contiguous nucleotides of the same strand. The distance associated with a triad is defined by the distance between the $C_\beta$ atom of the amino acid residue and the average position of the atoms of the nitrogen-base of the two nucleotides plus their complementary pairs in the opposite strand of the helix (16). The triad also has an associated amino-acid residue number in the protein and a dinucleotide position in the DNA, defined by the sequence position of the first nucleotide of the dinucleotide.

For the sake of the comparison of interfaces, we define the **interface** between a protein (e.g. transcription factor) and DNA as: the set of *triads* with associated distances shorter than 15 Å, and their associated amino-acid residue number and dinucleotide position (e.g. a $triad$ with amino-acid residue number $p$, dinucleotide in position $q$ and associated distance $d$ is represented as $(triad, d, p, q)$). This definition is extended up to 30Å in section 8c on the application of structural modeling. Amino-acid residue number and dinucleotide position are specific of the structure, consequently they are irrelevant for the comparison of interfaces and are not taken into account to calculate the score of similar-interface (see further). However, this definition of interface may be too rigorous when we have to compare two interfaces and the number of triads is too short. Therefore, we force requiring a minimum of 10 triads with their associated distances and positions of amino-acids and dinucleotides. If this minimum number of contacts is not achieved, we increase the cut-off distance (i.e. 15 Å) in steps of 1 Å until 10 or more triads can be assigned to the interface or we have reached a maximum of 30 Å.

Specific features can be added on a *triad*, defining an ***extended-triad***:

1) Hydrophobicity of the amino-acid. Amino-acid residues are split in **Polar (P):** {Arg, His, Lys, Asp, Glu, Ser, Thr, Asn, Gln, Cys, Gly} and **Non-polar (N):** {Ala, Ile, Leu, Met, Val, Phe, Trp, Tyr, Pro}.
2) Surface accessibility of the amino-acid. We use DSSP to calculate the percentage of accessibility of the residue in the unbound structure of the protein. If the percentage is smaller than 50% the amino-acid is **buried (B)**, otherwise it is **exposed (E)**.
3) Secondary structure of the amino-acid. We use DSSP to calculate the secondary structure of the protein. The amino-acid of the triad is either in regular secondary structure (**H** if in **$\alpha$-helix**, **E** if in **$\beta$-strand**), or in a **non-regular** secondary structure (**C**)
4) Nitrogenous bases: We classify nucleotides by their nitrogenous bases in two types, **purines (U):** {A, G} and **pyrimidines (Y):** {C, T}.
5) Closest strand. We use X3DNA to define the strands **forward** and **reverse** of the DNA. Next, we calculate the distance of all atoms of the two nucleotides to the $C_\beta$ of the amino-acid. We define the strand closest to the amino-acid (i.e. with the atom at minimum distance) as either the strand of the two nucleotides of the triad or the strand of their complementary pair in the opposite strand, which can be either **forward (F)** or **reverse (R)**.
6) Closest Groove. We calculate the distances between the $C_\beta$ of the amino-acid and the closest phosphates of the dinucleotides in both strands (i.e. the strand of the two nucleotides of the triad and its complementary). We calculate the positions of the closest phosphates in both strands (let be $P_f$ and $P_r$, backbone phosphates of nucleotides f and r, respectively). We select the closest phosphate of both and its corresponding strand. Let assume that $P_f$ is the closest phosphate and define its strand as "s", being "S" the opposite strand. Then, we consider the set of backbone phosphates in "S" around the position complementary of nucleotide "f" (6 nucleotides up and down). Depending on their distance to "f" (towards 22Å is a major groove and towards 12 Å a minor groove), we classify them as part of the minor or major groove with respect to nucleotide "f". This is a classification of 12 nucleotides around the complementary of "f" in two groups: 1) set at large distance (i.e. major groove); and 2) set at short distance (i.e. minor groove). Necessarily, $P_r$ is in the list classified in **major** or **minor groove**. We use the classification of $P_r$ to define the type of the closest groove of the amino-acid (i.e. we should say that this is the groove faced by the amino-acid, defined by the pair $P_f$ and $P_r$, in closest proximity to the amino-acid). The closest groove is defined as **major groove (A)** if $P_r$ is in the list classified in major, otherwise it is defined as **minor groove (I).**
7) Chemical group of the nucleotides. We distinguish two main chemical groups of each nucleotide, the nitrogenous base (**N**) and the backbone (**B**) that includes the phosphate and sugar. We calculate the distances between the $C_\beta$ of the amino-acid and the atoms of the two nucleotides and their complementary. We select the atom with the shortest distance as the closest atom between the nucleotides and the amino-acid. We define the chemical group of the nucleotides of the triad as the chemical group to which belongs the closest atom (i.e. **N** or **B**)

Added features of triads can also be used on their own as **feature-triads** (or environment triads), and every extended-triad has an associated feature-triad, both associated with the same distance, amino-acid number and dinucleotide position. As an example, let be a lysine residue and two nucleotides, adenosine and guanosine, forming the triad [K,(AG)] at 15.6Å, with lysine in residue number 32 and adenosine in 5, described as ([K,(AG)], 15.6, 32, 5). If lysine surface is mainly exposed to solvent, in a α–helix conformation and the closest strand of DNA is the forward strand, the closest atom of the two nucleotides is a phosphate and the amino-acid faces the minor groove, the extended-triad is [{K,(p-H-E)},{(AG),(UU-F-I-B)}], where added features are (p-H-E) for the amino-acid and (UU-F-I-B) for the dinucleotide. This produces a feature-triad defined as [(p-H-E),( UU-F-I-B)] at 15.6Å.

We require to define some functions on the sets of triads, extended-triads and feature-triads to extract some of the values collected from a complex structure and apply other functions:

$$f_{td}(triad, d, p, q) = (triad, d)$$
$$f_t(triad, d, p, q) = (triad)$$
$$f_d(triad, d, p, q) = (d)$$
$$f_a(triad, d, p, q) = (p)$$
$$f_n(triad, d, p, q) = (q)$$

The same functions are applied to extended-triads and feature-triads accordingly modified. For example, we first **apply $\boldsymbol{f_t}$** for all triads with d<15Å in order to calculate the **score of similar-interface**. Still, if any of the interfaces has less than 10 different elements (i.e. $(triad, d, p, q)$) the score of similar-interface may not be significant (e.g. if we compare one interface with only one element and another with many, the coincidence of one of them already achieves a score of similar-interface of 100%). Therefore, we use a less rigorous approach to calculate the interface by increasing the cut-off distance until both interfaces have at least 10 elements to perform the comparison and calculate the score of similar-interface.

We also define functions to substitute some of the elements of a triad, extended-triad or feature-triad (the example is given for *etriads* without loss of generality):

1) $\varepsilon_a(etriad, r)$ is a function that substitutes amino-acid residue "a" of the *etriad* by amino-acid-residue "r", with the corresponding change of hydrophobicity but preserving the rest of features and measures associated with the triad.
2) $\eta_\nu(etriad, n)$ is a function that substitutes dinucleotide $\nu$ by $n$, in $\Lambda = \{A, C, G, T\} \times \{A, C, G, T\}$, with the corresponding change of nitrogenous bases and preserving the rest of features and measures associated with the triad.

### 4. Statistical potentials

We use the definition of **statistical potentials** described by Feliu et al (17) and Fornes et al. (16) to define several **scoring functions** for the interaction between a protein and a DNA binding site using contact triads. We use triads, their associated measures (i.e. distance, amino-acid number and dinucleotide position) and their added features to calculate the frequencies per distance in bins of 1Å (i.e. intervals [0,1], [1,2], [2,3], [3,4], [4,5] etc.) up to 30 Å. From now, we use a distance of 1Å to define the bins without loss of generality, but we can use 2 Å and 3 Å, respectively defining sample bins in intervals [0,2], [2,4], [4,6], [6,8] etc. and [0,3], [3,6], [6,9], [9,12] etc. up to 30 Å. We also calculate the frequencies using the distance as cut-off (i.e. the frequency of triads at distance shorter than "d", with d= 1,2,3,4, etc.). We increase the cut-off distance in 1 Å without loss of generality, but we can increase the cut-off by 2 Å or 3 Å (i.e. defining cut-offs respectively at 2,4,6,etc. and 3,6,9,12, etc.). To obtain the frequencies, we first calculate the size (cardinality, defined by the function "Card") of the sets of triads, associated with a distance (d), taken from the set of structures of protein-DNA interactions ($PDB_{DNA}$) and grouped by their associated distance, limited to a maximum of 30Å (i.e. with $i \in [1,30]$). The set of triads, associated with distances, amino-acid residue-number and dinucleotide position, is named ***3Dset***. Then, frequencies are defined using functions defined in section 3 as follows:

$N_D(triad, i) = Card(\{f_{td}(x) | where\ x \in 3Dset\ and\ (i-1) < d \leq (i)\})$ (eq. 1)
$N_c(triad, i) = Card(\{f_{td}(x) | where\ x \in 3Dset\ and\ 0 < d \leq i\})$ (eq. 2)

Where $x = (triad, d, p, q)$ is a triad associated with a distance d, amino-acid residue-number $p$ and dinucleotide position $q$, taken from the set *3Dset*, $N_D$ is defined using bins and $N_c$ using cut-offs. Similarly to ***3Dset*** we define the sets ***e3Dset*** and ***f3Dset*** for extended-triads (*etriad*) and feature-triads (*ftriad*), and calculate $L_D$, $L_c$ , $M_D$ and $M_c$ with $i \in [1,30]$ as:

$L_D(etriad, i) = Card(\{f_{td}(x) | where\ x \in e3Dset\ and\ (i-1) < d \leq (i)\})$ (eq. 3)
$L_c(etriad, i) = Card(\{f_{td}(x); where\ x \in e3Dset | 0 < d \leq i\})$ (eq. 4)
$M_D(ftriad, i) = Card(\{f_{td}(x) | where\ x \in f3Dset\ and\ (i-1) < d \leq (i)\})$ (eq. 5)
$M_c(ftriad, i) = Card(\{f_{td}(x) | where\ x \in f3Dset\ and\ 0 < d \leq i\})$ (eq. 6)

Then, we define the frequencies (F for triads, G for extended-triads and H for feature-triads) as:

$F(triad, i) = N(triad, i) / \sum_{j=1}^{30} N(triad, j)$ (eq. 7)
$G(etriad, i) = L(etriad, i) / \sum_{j=1}^{30} L(etriad, j)$ (eq. 8)
$H(ftriad, i) = M(ftriad, i) / \sum_{j=1}^{30} M(ftriad, j)$ (eq. 9)

Where N can be $N_D$ or $N_c$, L can be $L_D$ or $L_c$ , and M can be $M_D$ or $M_c$, depending on the approach to group the triads. This definition forces us to consider independent the groups obtained by cut-offs, instead of using the ratios with respect to the limit at 30 Å. We tested in artificial data that this approach preserves the curve of the statistical potential similar to the classical definition by bins of distances, but it's less affected by the scarcity of data.

To define a reference-state for the statistical potential, we require two more frequencies, one for $N_D$ and another for $N_c$ , using the **total number of triads** in the database (***triads***):

$O(i) = \sum_{triads} N(triad, i) \,/\, \sum_{j=1}^{30} \sum_{triads} N(triad, j)$ (eq. 10)

Where $triad \in triads$, and it's easy to proof that:

$$\sum_{j=1}^{30} \sum_{triads} N(triad, j) = \sum_{j=1}^{30} \sum_{ftriads} M(ftriad, j) = \sum_{j=1}^{30} \sum_{etriads} L(etriad, j)$$

(eq.11)

Where ***ftriads*** is the set of feature-triads and ***etriads*** the set of extended-triads, with $ftriad \in ftriads$ and $etriad \in etriads$. Using these definitions and following previous works (16), we define the potentials E3DC, ES3DC and PAIR per *triad* and distance *d*, using the round value of d (i.e. k), as follows:

$k = 1 + int(d - 1)$

$PAIR(triad, d) = K_B T log_e(O(k)) - K_B T log_e(F(triad, k))$ (eq.12)

$ES3DC(etriad, d) = K_B T log_e\big(H(ftriad, k)\big) - K_B T log_e(G(etriad, k))$ (eq. 13)

$E3DC(ftriad, d) = -K_B T log_e\big(O(k)\big) + K_B T log_e(H(ftriad, k))$ (eq. 14)

Where, F, G, H and O are frequencies calculated by bins or using cut-offs.

The total potential of an interaction is calculated as the sum of the corresponding potential of all triads, feature-triads and extended-triads at distances shorter than 30Å. We use each potential to **score the quality** (or potentiality) of the interaction. Let be I, E and D the sets defined respectively as the set of all triads, extended-triads and feature-triads with their associated distances (d), amino-acid residue number (p) and dinucleotide position (q) in the binary interaction of a TF-DNA structure. Therefore, we define the **energy-based *scores*** (as they are based on total potentials) as:

$PAIR = \sum_{(triad,d,p,q) \in I} PAIR(triad, d)$ (eq. 15)

$ES3DC = \sum_{(etriad,d,p,q) \in E} ES3DC(etriad, d)$ (eq. 16)

$E3DC = \sum_{(ftriad,d,p,q) \in D} E3DC(ftriad, d)$ (eq. 17)

Similarly, we also use the potentials defined for *triads, ftriads* and *etriads* as *scores* of the quality of a single interaction between one dinucleotide and one amino-acid residue. Then, the potential ES3DC of an extended-triad associated with distance d can be rewritten as:

$$ES3DC(etriad, d) = E3DC(ftriad, d) + K_B T log_e\big(O(k)\big) - K_B T log_e(G(etriad, k))$$

(eq. 18)

Where $k = 1 + int(d - 1)$. We then define two scoring terms, one distance independent ($ES3DC_{di}$) and another distance dependent ($ES3DC_{dd}$), as follows:

$ES3DC_{di}(etriad) = K_B T log_e\big(\sum_{j=1}^{30} L(etriad, j) / \sum_{j=1}^{30} \sum_{triads} N(triad, j)\big)$ (eq.19)

$ES3DC_{dd}(etriad, d) = -K_B T log_e(L\,(etriad, k) / \sum_{triads} N(triad, k))$ (eq. 20)

Using eq. 8, eq. 19 and eq. 20, we rewrite $ES3DC(etriad, d)$ as:

$$ES3DC(etriad, d) = E3DC(ftriad, d) + ES3DC_{di}(etriad) + ES3DC_{dd}(etriad, d)$$
(eq. 21)

And the global energy-based *scores:*

$$ES3DC_{dd} = \sum_{(etriad,d,p,q)\in E} ES3DC_{dd}(etriad, d)$$
$$ES3DC_{di} = \sum_{(etriad,d,p,q)\in E} ES3DC_{di}(etriad)$$
(eq.22)

We consider $ES3DC_{dd}$ to evaluate the quality of an interaction because it is more specific than PAIR, as it uses extended-triads and distances, and it is also highly sensible, because it is normalized over the total of triads at a given distance with independence of the features. However, we have to note that this can only be considered as a scoring of quality, because the complete statistical potential requires the other terms E3DC and $ES3DC_{di}$ to complete ES3DC in equation 21.

Finally, due to the scarcity of data the curves of statistical potentials may be jagged. Therefore, we use a sliding window of approximately W samples defined by distances (by bins or cut-off) to **smooth the potential curves**. Let be $Scr(d)$ a distance-dependent score, as defined previously, then we define the smoothed score, $Scr_{smooth}$, as:

$$Scr_{smooth}(d) = \frac{1}{size_{samples}} \sum_{k=k_{min}}^{k_{max}} Scr(k)$$
(eq.23)

Where $k_{max}$ and $k_{min}$ are defined as:

$$k_{max} = \min(30;\ 1 + int(d-1) + int(W/2))$$
$$k_{min} = \max(0;\ 1 + int(d-1) - int(W/2))$$

Where, $size_{samples}$ is the total of bins between $k_{min}$ and $k_{max}$ with defined score ($Scr$), which is around W (i.e. $size_{samples} \approx W$)

### 5. Solving scarcity of data by using Taylor's polynomial series approach

The main problem of using extended-triads instead of triads is the size of data required to fill the distribution by distances for all types. The current number of structures is not enough to complete such a large distribution. Consequently, for some extended-triads the distribution by their associated distances is scarce and discontinuous. In order to fill these gaps, we propose an approximation based on a Taylor's polynomial approach of the statistical potentials. We generalize the approach over a potential function P as follows:

Let P be a potential defined over extended-triads (i.e ES3DC, $ES3DC_{dd}$ or $ES3DC_{di}$), then it can always be expressed in terms of a function $\theta$ of the extended-triad with associated distance $d$ (rounded to $k = 1 + int(d-1)$), and a constant (Q) independent of the extended-triad:

$$P(etriad, d) = Q - K_B T log_e(\theta(etriad, k)) \quad \text{(eq. 24)}$$

We approach $\theta$ for an extended-triad, with amino-acid residue "a" and associated distance rounded to k, as:

$$\theta(etriad, k) = \theta_0(etriad, k) + \sum_{r \in (A \neg a)} \phi(r, a)\theta_0(\varepsilon_a(etriad, r), k) \quad \text{(eq. 25)}$$

Where, $\theta_0$ is function $\theta$ calculated with raw data and without approximation, $\varepsilon_a(etriad, r)$ is the function defined in section 3 to substitute amino-acid residue "a" of the *etriad* by amino-acid-residue "r"; $\phi(r, a)$ is the probability of substituting amino-acid residue "r" by "a"; A is the set of 20 amino-acids and $A \neg a$ is the set of all amino-acids except "a". According with these definitions, if there is no data for $\theta_0(etriad, k)$, we approach it by the weighted average of all other potential amino-acid residues ("r") for which this data exist, located in the same featured triad and associated distance, that can substitute amino-acid "a". We use the substitution matrix BLOSUM62 to calculate the probability of substitution of "r" by "a".

If $\theta_0(etriad, k) = 0$, we select the amino-acid "b" for which we get the maximum value of $\phi(r, a)\theta_0(\varepsilon_a(etriad, r), k)$, this is:

$max = maximum(\{\phi(r, a)\theta_0(\varepsilon_a(etriad, r), k); \forall r \epsilon A \neg a\})$ ; and
$max = \phi(b, a)\theta_0(\varepsilon_a(etriad, b), k)$ (eq.26)

If $max = 0$, we cannot do the approach because at this distance we have no information of any other residue. Otherwise, if $max \neq 0$, we rewrite eq. 25 as:

$$\theta(etriad, k) = \max\left(1 + \frac{1}{max}\sum_{r \in (A \neg a, b)} \phi(r, a)\theta_0(\varepsilon_a(etriad, r), k)\right) \quad \text{(eq. 27)}$$

Where $A \neg a, b$ is the set of all amino-acids except "b" and "a".

Then, we define $P_0$ as the potential obtained without approximation and obtain $\theta_0$ as a function of $P_0$:

$$\theta_0(etriad, k) = e^{Q/K_B T} e^{-P_0(etriad,k)/K_B T} \quad \text{(eq. 28)}$$

We rewrite max as a function of the potential

$$max = \phi(b, a) e^{Q/K_B T} e^{-P_0(\varepsilon_a(etriad,b),k)/K_B T} \quad \text{(eq. 29)}$$

And we also define the difference of the potential by substituting amino-acid "a" by any residue "r" with respect to the substitution with the maximum potential (i.e. amino-acid "b"), as:

$$\Delta P_0(b, r, k) = P_0(\varepsilon_a(etriad, r), k) - P_0(\varepsilon_a(etriad, b), k) \quad \text{(eq. 30)}$$

Then, substituting eq. 27 in eq. 24, and using eq. 28, eq.29 and eq. 30, results in:

$$P(etriad, d) = P_0\big(\epsilon_a(etriad, b)\big) - K_B T log_e\big(\phi(b,a)\big) - K_B T log_e\left(1 + \sum_{r\in(A\neg a,b)} \frac{\phi(r,a)}{\phi(b,a)} e^{-\Delta P_0(b,r,k)/K_B T}\right)$$

(eq. 31)

Here we apply Taylor's polynomial series of the logarithm function, assuming that "max" is higher than any other value for the rest of residues and we use Maclaurin series (i.e. centered in 0):

$$log_e(1+x) \approx x - \frac{1}{2}x^2 + \frac{1}{3}x^3 - \cdots + \delta$$

Then, we rewrite eq. 31 using only the first term of Maclaurin series as:

$$P(etriad, d) \approx P_0\big(\epsilon_a(etriad, b)\big) - K_B T log_e\big(\phi(b,a)\big) - K_B T \sum_{r\in(A\neg a,b)} \frac{\phi(r,a)}{\phi(b,a)} e^{-\Delta P_0(b,r,k)/K_B T} + \delta$$

(eq.32)

Similarly, if $\theta_0(etriad, k) \neq 0$, we can use the same definitions of *max* and "b" , $\theta_0$ and $P_0$ to rewrite eq. 24 as:

$$P(etriad, d) = P_0(etriad, k) - K_B T log_e\left(1 + \frac{max}{\theta_0(etriad, k)}\left(1 + \sum_{r\in(A\neg a,b)} \frac{\phi(r,a)\theta_0(\varepsilon_a(etriad, r), k)}{max}\right)\right)$$

eq. 33

And using eq. 28, eq. 29 and eq.30, applying Maclaurin polynomial series:

$$P(etriad, d) \approx P_0(etriad, k) - K_B T \phi(b,a) e^{\Delta P_0(b,a,k)/K_B T}\left(1 + \sum_{r\in(A\neg a,b)} \frac{\phi(r,a)}{\phi(b,a)} e^{-\Delta P_0(b,r,k)/K_B T}\right)$$

eq.34

With the requirement that $0 < \phi(b,a)e^{\Delta P_0(b,a,k)/K_B T} \ll 1$ in order to satisfy the conditions applied for the Taylor polynomial-series approach. Finally, as in section 5, potential curves are **smoothed** by using a sliding window of W samples defined by distances (by bins or cut-off) to smooth the potential curves.

### 6. Z-scores

We have to note that frequencies are always smaller than 1, consequently their logarithm is smaller than 0. However, statistical potentials are not necessarily negative, because by definition the use of a reference state is required, which is obtained by the sum of all triads at a given distance (also *ftriad* or *etriad*, depending on the potential of interest). Consequently, the comparison with a reference state can change the sign of the final sum of terms (for example, PAIR and ES3DC can be negative while E3DC may be positive). The variability of signs of the potentials affects the criterion of quality of the scores and they may become unclear. Nevertheless, indistinctly of the sign, the best interaction between an amino-acid and a dinucleotide is produced at the distance where PAIR and ES3DC are minimum, because it implies the highest frequency of a *triad* (or *etriad*) with respect to all triads (or feature-triads). For example, $ES3DC_{dd}$ has a minimum for the highest frequency of an extended-triad with respect to all triads. We then define *z-scores* in order to follow a criterion that incorporates the sign to score the quality of the interaction between an amino-acid and a dinucleotide as a function of the distance. We wish that the z-score identifies simultaneously the best distance associated with a triad and the best pair formed by one amino-acid and one dinucleotide. Consequently, we construct a ***zscore* function** with any type of *score*, applying without loss of generality on an extended-triad (*etriad*) and an associated distance d, as:

$$zscore(score) = \frac{score(etriad, d) - \mu_A\big(score(etriad, d)\big)}{\sigma_A\big(score(etriad, d)\big)}$$

(eq. 35)

Where: A is the set of all amino-acids (i.e. $Car\{A\} = 20$), and we use the classical functions of **average** ($\mu$) and **standard deviation** ($\sigma$), defined as:

$$\mu_A\big(score(etriad, d)\big) = \frac{1}{Car\{A\}} \sum_{r \in A} score\big(\epsilon_a(etriad, r)\big)$$

$$\sigma_A\big(score(etriad, d)\big) = \frac{1}{Car\{A\}} \sqrt{\sum_{r \in A} \Big(score(etriad, d) - \mu_A\big(score(etriad, d)\big)\Big)^2}$$

Notice that we use the function $\varepsilon_a(etriad, r)$ as above and that we use the term *score* instead of potential (P in previous section) to emphasize that this is used only as a criterion of the quality of the interaction.

It's easy to proof that this definition satisfies our requirements for the new function *zscore*: 1) the minimum value of *score* produces a negative value of *zscore* (and *vice-versa*, the maximum yields positive); 2) as the *zscore* compares the *score* of a particular amino-acid with all other, the *etriad* becomes ranked from minimum to maximum allowing us to select the best residue among amino-acids in the same position, depending on the quality criterion of the score. Finally, if we choose to smooth the curves dependent on distance (see section 4), the *zscore* has to be calculated first with the unsmoothed scores and subsequently smoothed to avoid introducing biases.

### 7. General and family-specific potentials

One of the requirements for the construction of the statistical potentials is to remove redundancies of complex structures. The motivation that instigates such reduction is the presence of structures of transcription factors of the same family, with very similar sequences in both protein and DNA binding sites. These structures would introduce an important bias in the statistical potentials, limiting the capacity to score and predict the bindings. Therefore, we define two different types of potentials: 1) **general potentials**, requiring the minimum bias that avoids redundancies of similar interactions; and 2) **family-specific potentials** that allow the similarity but can only be applied on TFs for which we know their family (i.e. the group of folds in which they belong). For "general potentials", in order to obtain an unbiased set of TF-DNA structures, we reduce the set of *triads* and their associated distances, amino-acid residue-number and dinucleotide position (*3Dset*) by excluding complexes of structures with more than 40-50% of score of similar-interface. For family-specific potentials we wish to accept an important degree of similarity that avoids too similar or redundant structures. Consequently, for "family-specific potentials" we use groups of folds and exclude complexes with more than 70-80% of score of similar-interface between interfaces of TFs in the same group of fold. We note that "family-specific potentials" are constructed specifically for each group of folds.

Besides differences on the construction of potentials, there is also an important difference on its application. The "general potential" can be applied to all TF structures automatically without further requirements, while for the application of "family-specific potentials" on a TF-DNA interaction we first need to assign the structure to one of the groups of folds (see section 2 on "Databases"), then we apply the "family-specific potential" obtained for this fold. On the one hand, if the structure of the TF-DNA structure is known we can assign the "fold" group straight-forward. On the other hand, If the structure of the TF-DNA structure is not known but has been modelled, then we apply the "family-specific potential" of the template with which the TF-DNA interaction was modelled.

### 8. Structural modeling of TFs complexed with DNA

Given the sequence of a TF (named protein target) and a DNA binding site fragment (named DNA target), we have developed an automated method to obtain the structure of the complex by means of homology modelling using the program MODELLER (6). First, we search potential templates of the protein target among the set of sequences with known structure in $PDB_{DNA}$ using BLAST (18). We select the first match, with highest score and minimum E-value, to define a fragment of the sequence of the target localized by the number of one residue in the N-tail position and another in the C-tail position. Next, we select all codes of the matches of BLAST with E-value smaller than 0.1, where the sequence fragment of the TF in the match overlaps in more than 50% with the selected fragment of the target in the first match. We define the minimum position at the N-tail (MinNt) and maximum position at the C-tail (MaxCt) of the target sequence using the BLAST alignments with the sequences of the selected codes. We collect the structures and sequences of the selected codes and align the sequences with the fragment of the target between MinNt and MaxCt using the program *matcher* of the EMBOSS package (3). We select only those sequences where the alignment with the fragment of the target produces a sufficiently high percentage of identical residues as a function of the length of the alignment (i.e. in agreement with the curve of the Twilight-Zone described by Rost (19)). The final selected sequences correspond to the templates for modelling the structure of the sequence fragment of the target between MinNt and MaxCt. We use the alignment obtained with *matcher* and the sequence and structure of each template to construct one or more models with MODELLER (depending on the total amount of models we wish to obtain). Each model is based on a different template and is stored with the name of the TF target, the code of the selected template, the start and end residues of the fragment (MinNt and MaxCt), and a number for the collection (in case we model more than one structure per template). We include the DNA structure of the template in the model of the target as a co-factor (heteroatom).

After modeling the main fragment (sequence of the protein target defined between MinNt and MaxCt), we proceed similarly with the remaining fragments of the protein target (i.e. maximum two fragments: between 1 and MinNt, and between MaxCt and the last residue of the target). This implies an iterative procedure by which the TF target is split and modelled, producing models for one (if no more matches are obtained with BLAST) or several domains. Finally, we keep only those models with more than 5 contacts between the protein and the DNA structure to proceed with further modelling and analyses. In particular, we still need to model the structure of the DNA target and substitute the structure of DNA of the template (see further).

#### a. Protein monomer/dimer modeling

When facing the selection of templates to model the conformation of a TF, the automated approach can select a template acting in the form of a dimer. However, as we select sequences among all chains of known structures, the automated approach models the structure as a monomer, one for each chain from the same PDB file of the template. In order to use the correct form, i.e. the expected for a particular family of TF, we have predefined those families that usually act as dimers (see section 2): “B3”, “bHLH”, “bZIP”, “Leafy”, “MADS box”, “Rel”, “Nuclear Receptor”, “STAT”, and “Zinc cluster”. By default, we proceed by modelling the homodimer or monomer structure as the corresponding conformation in the template, unless the user specifically requires a specific model. In order to model the homodimer, we select only homodimer template structures and duplicate the sequence alignment of the monomer sequence between the target and each template; then, we run MODELLER [Webb, 2017 #32] with each template as defined in the examples of multiple-chain modelling.

### b. Protein homo/hetero dimer modeling

To model a heterodimer (or force a homodimer) we first need to select dimer templates out of all potential templates of the target. We use a file with the two sequences of the dimer in FastA format. Each sequence of the target dimer (both are the same in case we model a homodimer) is aligned and scored with the sequence of the potential templates using BLAST (see above). We select only those templates sharing the same PDB code from $PDB_{DNA}$, but different chain code for each sequence of the target. This implies that we select a dimer complex structure and each sequence can be modelled with a different chain. Besides, in contrast to monomer modelling, only the first sequence of the file can be split and subsequently tested and modelled (see above). This restriction is forced in order to avoid multiple solutions, because the application is specifically selected by a user who decides the sequences in the input and expects an answer in the shortest time (i.e. this is applicable only in a web-service). We realign each sequence of the target with the corresponding sequences of the template using *matcher* and run MODELLER as defined in multiple-chain modelling including the DNA structure as heteroatoms.

### c. DNA modeling

After modeling several conformations of the protein target with each template and bound with the same DNA of the corresponding template, we still need to substitute the DNA structure in the model by the DNA of the target. With this purpose, we define some previous requirements to do the modeling of the DNA target: 1) the target DNA sequence belongs in the strand defined by the first DNA chain in the PDB format file, which is named chain of reference; 2) the target DNA sequence corresponds only to the DNA binding site, oriented from 5' to 3' of the chain of reference, numbering the nucleotides in increasing order; 3) we assume that the binding site sequence is limited between 5' and 3' positions by the region in contact with the target protein (i.e. the DNA region before 5' and after 3' is not in contact or not in the interface); and 4) the model of DNA preserves the same conformation as in the original template.

We first assume that the sequence of the DNA target has the same length of the binding site (i.e. all other potential nucleotide residues before and after the DNA target sequence are not in contact with the TF). We model the DNA structure by substitution of the nucleotides in each model with the corresponding nucleotides of the target. We use the program X3DNA to substitute the nucleotides of one strand and automatically model its corresponding pair. Therefore, the alignment between the DNA sequence of the target and the sequence of the corresponding DNA strand in the template is required to apply the substitution. To obtain this alignment we first trim the DNA structure of the model constructed in sections "8.a" or "8.b" (note that they still belong on the structure of the DNA original template) by removing all nucleotides that are not in contact with the TF before 5' and after 3' of the chain of reference. The alignment is then obtained by the exact match between the length of the DNA sequence of the template and the length of the DNA target sequence, yielding a one to one correspondence in the same order of the nucleotide sequence.

Alternatively, instead of trimming the DNA of the chain of reference from the template, we can suggest the user to extend the DNA target sequence in order to preserve a one to one correspondence between the sequences of the DNA target and the DNA template (i.e. the length of the DNA sequence of the interface is shown in the web-service).

If the length of the DNA sequence of the target is longer than the length of the DNA sequence in contact with the TF, we need to define the margins of the binding site. We trim the DNA structure in the model by removing the nucleotides before and after the binding site defined by the contacts with the TF as before in the directions of 5' and 3'. However, in order to consider the impact of long-range interactions, we extend the definition of the interface up to 30Å, as used in the definition of the statistical potentials, and all nucleotides in triads with associated distances under 30Å are considered in contact with the TF. If the length of the DNA target is still different than the length in the structure it implies the sequence is incorrect or incomplete and the modeling stops, otherwise we proceed as above.

**d. Modeling protein-protein interactions.**

To model a protein-protein interaction (target) we use a file with two protein-sequences (e.g. A and B) in FastA format in a similar way as for heterodimers. Each sequence is aligned and scored with the sequences of potential templates found by BLAST (see above) when searching on the complete set of structures (i.e. all structures from PDB). Both target sequences are split by parts according to the BLAST search as explained above. We select those templates that have the same PDB code but different chain for each fragment of the target sequences (one fragment of A and another of B). Each sequence-fragment is then modelled with a different chain, the use of several templates generates several models. Consequently, this procedure allows for modeling several interfaces of both sequences (A and B). We realign with *matcher* each sequence-fragment of the target with the corresponding sequences of the single-chain templates. Then, we use the multiple-chain approach of MODELLER to model several structures of the interactions of the split parts of both sequences. Structural models are validated by confirming that their interface has more than 5 residue-residue contacts and the number of clashes and perforations between surfaces is limited.

**e. Modeling C2H2-ZF structures for B1H**

The DNA binding sequence of each testing 3-domain C2H2-ZF protein is formed by 9bp nucleotides, a set of 3bp bound by each individual domain. For the selection of the binding sequences associated with each finger domain we use the same sequences as in the B1H experiment (14): 1) for the selection of F2, the finger sequences in F1 (N-tail domain) and F3 (C-tail domain) involved in the interface are RSDNLRA(F1) and RSANLVR (F3), respectively binding AAG and GAG; and 2) for the selection of F3, the finger sequences in F1 (N-tail domain) and F2 (inner domain) involved in the interface are RSDELTR (F1) and RSDNLRA (F2), respectively binding GCG and AAG. The structure of Zif268 binding DNA is modelled with three different template structures to introduce structural variability: 1p47 (chain A), 1zaa (chain C) and 1g2f (chain C). We compare the sequence of Zif268 used in the experiment with the templates by a sequence alignment with CLUSTALW (20). We identify by WT the original sequence, and by F2 and F3 the sequences used for the selection of F2 and F3 binding sequences, labeling by “X” the amino-acids that are modified. This alignment is shown here, highlighting in bold and red the attention of the specific binding sequence of each finger (blue box for F1, yellow box for F2 and green box for F3):

```
F2              GTERPYACPVESCDRRFSRSDNLRAHIRIHTGQKPFQCRICMRNFSXXXXLXXHIRTHTG
F3              GTERPYACPVESCDRRFSRSDELTRHIRIHTGQKPFQCRICMRNFSRSDNLRAHIRTHTG
WT              GTERPYACPVESCDRRFSRSDELTRHIRIHTGQKPFQCRICMRNFSRSDNLRAHIRTHTG
1g2f_C          -MERPYACPVESCDRRFSQKTNLDTHIRIHTGQKPFQCRICMRNFSQQASLNAHIRTHTG
1p47_A          --ERPYACPVESCDRRFSRSDELTRHIRIHTGQKPFQCRICMRNFSRSDHLTTHIRTHTG
1zaa_C          ---RPYACPVESCDRRFSRSDELTRHIRIHTGQKPFQCRICMRNFSRSDHLTTHIRTHTG
                   ***************    *  *********************    *  *******

F2              EKPFACDICGRKFARSANLVRHTKIHLRGS
F3              EKPFACDICGRKFAXXXXLXXHTKIHLRGS
WT              EKPFACDICGRKFARSANLVRHTKIHLRGS
1g2f_C          EKPFACDICGRKFATLHTRTRHTKIHLRQK
1p47_A          EKPFACDICGRKFARSDERKRHTKIHLRQ-
1zaa_C          EKPFACDICGRKFARSDERKRHTKIHLR--
                **************       ******
```

After modeling the structure of Zif268, we complete the complex by modeling the structure of the DNA binding sequence. However, each template has DNA sequences of different length that do not correspond with the DNA used in the experiment. The full DNA sequence of the experiment is longer (29bp) than the binding (9bp), which is shown next, embedded in positions 11 to 19 (nucleotides highlighted), labelling by “N” those under test. Two DNA sequences are considered depending on the experiment, one for the selection of F2 and another for F3:

```
F2:    5'- GCGGCCGCAAGAGNNNAAGTAACGAATTC - 3'
F3:    5'- GCGGCCGCAANNNAAGGCGTAACGAATTC - 3'
```

The structure of the full DNA sequence bound by Zif268 is obtained with the program X3DNA(2) by modifying the DNA structure in the complex. First, we locate the 9bp binding region in the structure of the template and identify the positions at 5’ and 3’ (*first* and *last*). Next, we construct two frames of B-DNA structure with the lengths required to extend the template at 5’ and 3’ up to 29 nucleotides. The lengths in both sides depend on the location of the 9bp binding sequence and the DNA sequence of the experiment. Then, we use 3DNA/DSSR to perform a least-squares fitting that locates each base reference frame in the *first* and *last* positions. Finally, the structure of DNA is completed with the right sequence following the same procedure as above (in 8.c).

We also model several structures with the complex of Zif268 binding a non-specific DNA region using the same approach. These structures are be used as non-binding examples. The non-binding sequence is taken randomly by selecting a region of the sequence of the weak promoter GAL1, constructing 10 DNA fragments of 29bp for each known binding. A potential binding test in this region, which is part of the B1H experiment, may well represent the background. The forward weak promoter sequence of GAL1 is formed by 118bp that are shown here:

```
5'- GAGATTAAGGAGCAGAAGGGGTGACAGCCCTCCGAAGGAAGA
    GAGATTAAGCTCTCCTCCGTGCGTCCTCGTCTTCACCGGTCG
    CGTTCCTGAAACGCAGATGTGCCTCGCGCCGCACTGCTCCG - 3'
```

### 9. Use of experimental TF-DNA binding to calculate statistical potentials

One of the main problems to obtain statistical potentials for all families and folds of TFs is the scarcity of known interactions. Even if we address this problem by using a theoretical approach such as the use of Taylor's polynomial series, we still require enlarging the number of experimentally known structures. In fact, what we need to enlarge is the number of interacting triads. Therefore, here we propose to use the experimental knowledge of TF-DNA interactions to derive interacting triads without requiring the complete knowledge of TF-DNA complex structures. We then use the sets of derived triads associated with distances to calculate the statistical potentials.

We select all TFs from the set of CIS-BP with experimentally known interactions with DNA by means of PBM. For each TF in this set, we collect all the 8-mers with accepted interactions and check if the structure is known. If the structure is known (i.e. there is a specific file in PDB with the structure of the interaction), we confirm that the DNA sequence extracted from the structure is among the 8-mers. This is done by a sequence alignment without intra-sequence gaps with all 8-mers classified as positive bindings. Then, we use the PWM obtained with all positive binding 8-mers to align the DNA sequence extracted from the structure and all positive binding 8-mers. We skip all alignments with intra-sequence gaps and trim the rest of alignments by removing the tails with gaps at the beginning and end. This results in an alignment with a length of maximum 8 nucleotides. For each alignment, we define a mapping function ($map_D$) between the fragment of the DNA sequence from the structure ($S_{DNA}$), as seen in the alignment, and any of the positive binding 8-mer sequences ($S_{8-mer}$), as follows:

$$n_m, q_n = map_D(\nu_m, q_\nu) \qquad \text{(eq.36)}$$

Where $n_m$ is a dinucleotide, with $n_m \in S_{8-mer}$, $\nu_m$ is also dinucleotide, with $\nu_m \in S_{DNA}$, $m$ is the position of the dinucleotide in the alignment between $S_{8-mer}$ and $S_{DNA}$, $q_\nu$ is the position of the dinucleotide $\nu_m$ in $S_{DNA}$, $q_n$ is the position of the dinucleotide $n_m$ in $S_{8-mer}$, and the position of each dinucleotide is defined (i.e. equal to) the position of the first nucleotide in the DNA sequence. For the sake of simplicity, when the length of $S_{DNA}$ is the same as $S_{8-mer}$ and the position in the alignment, $m$, coincides with $q_\nu$ and $q_n$ (i.e. $m = q_n = q_\nu$), we write:

$$n_m = \widehat{map}_D(\nu_m) \qquad \text{(eq.37)}$$

We can define a set of mapping functions with the alignments of all the DNA sequences extracted from the structures of TF-DNA complexes in $PDB_{DNA}$ that are aligned with positive binding 8-mer sequences in CIS-BP. Then, we use the dinucleotide substitution function (as defined in section 3), $\eta_v(etriad, n)$, of an extended-triad, *etriad*, containing a nucleotide $v$ which is substituted by *n* in the dinucleotide, to generate more extended-triads.

There are still TFs from the set of CIS-BP with PBM experiments for which the structure is not known but can be modelled. This implies that the sequence of the TF can be aligned with sufficient percentage of identical residues to ensure its modeling (see above in section 8). Then, we define another mapping ($map_P$) between the protein sequence of the TF in CIS-BP and the TF sequence of a known structure (template), with:

$$r_m, \; p_r = map_P(a_m, p_a)$$

(eq.38)

Where, $a_m$ is an amino-acid residue in the sequence of a template and $r_m$ is the amino-acid in the sequence of a TF in CIS-BP, in position $m$ of the alignment of both TF sequences that correspond with positions $p_r$ and $p_a$ for $r_m$ and $a_m$, respectively. Also, for the sake of simplicity, if the position in the alignment, $m$, coincides with $p_a$ and $p_r$ (i.e. $m = p_r = p_a$ ) we write:

$$r_m = \widehat{map}_P(a_m)$$

(eq.39)

We define the set of all mapping functions with all the alignments between the sequences of TFs in CIS-BP (with PBM experiments) and the TFs with known structure (complexed with DNA). Also, we define $etriads(a, v)$ as the set of extended-triads, extracted from the 3Dset, containing amino-acid residue $a \in A$ (the set of 20 amino-acids) and dinucleotide $v \in \Lambda = \{A, C, G, T\} \times \{A, C, G, T\}$.

We remind the definition of $\varepsilon_a(etriad, r)$ as the function that substitutes amino-acid residue "a" of an extended-triad, *etriad*, by the amino-acid residue "r". With all these definitions, we increase the set of extended-triads of the e*3Dset* to *e3Dset'*, using all the mapping functions $map_D$ (simplified as $\widehat{map}_D$) and $map_P$ (simplified as $\widehat{map}_P$). We use the simplified maps without loss of generality, as all sequences and alignments can be renumbered. Both mappings, $\widehat{map}_D$ and $\widehat{map}_P$, are respectively defined with: 1) the alignments of the DNA sequences extracted from the structures in the PDB aligned with positive binding 8-mer sequences; and 2) the TFs that can be modelled using the alignment with their structural templates (the set is defined as $PBM_{3Dset}$). The new set of extended-triads, *e3Dset'*, is defined as:

$$e3Dset' = \left\{ (etriad, d, p, q) \middle| \begin{array}{c} with\ etriad = \eta_v\left(\varepsilon_a\left(x, \widehat{map}_P(a)\right), \widehat{map}_D(v)\right);\ x \in etriads(a, v); \\ x\ associated\ with\ d, p\ and\ q;\ \forall a \in A;\ \forall v \in \Lambda;\ \forall\ \widehat{map}_D, \widehat{map}_p \in PBM_{3Dset} \end{array} \right\}$$

(eq. 40)

And we recalculate $L_D$ and $L_C$ in equations eq.3 and eq.4 as:

$$L_D(etriad, i) = Card(\{f_{td}(x) | where\ x \in e3Dset' and\ (i-1) < d \leq (i)\}) \text{ (eq. 41)}$$
$$L_c(etriad, i) = Card(\{f_{td}(x) | where\ x \in e3Dset' and\ 0 < d \leq i\}) \text{ (eq. 42)}$$

Similar approach is also taken for the sets of *triads* and *ftriads*, modifying accordingly the corresponding functions to substitute amino-acid and dinucleotide residues of a triad or a featured-triad. In order to avoid biases produced by binding similar sequences in the PBM experiment we exclude complexes of structures with more than 50% of score of similar-interface when applying the mapping and substitution of the dinucleotide sequence. To improve the speed of the calculation we also neglect DNA sequence bindings of the same TF modelled with the same template if it differs in less than 2 nucleotides of a previous selected sequence. However, we only use this reduction to calculate the general potential, while for family potentials we allow all DNA bindings to avoid a considerable computational cost (using a limit on 80% of score of similar-interface for members of the same fold has little effect but implies a long time of calculation).

We proceed similarly with B1H experiments on C2H2-ZF family. For each finger (F2 and F3) and combination of 3bp nucleotides, we collect all protein sequences producing significant binding signal in the B1H experiment. We use the modelled structures of the DNA testing sequence of 29bp with different templates and introduce the mappings for the DNA sequence and the modified residues in F2 or F3 from the multiple sequence alignment. For the DNA sequence the mapping is on the 3bp modified nucleotides, affecting 4 dinucleotides, while for the protein sequence the mapping affects 6 amino-acids, both mappings being different for F2 and F3 selections. This is, for the DNA sequence the mapping is $n_m = \widehat{map}_D(v_m)$, where $n_m$ is a dinucleotide of the 3bp under test, $v_m$ is a dinucleotide of the 29bp of the modelled template, and the position, $m$, is between 11 and 19 (11-13 for F3 and 14-16 for F2). For the substitution of residues of Zif268 in the interface we require a mapping of the native sequences RSANLVR (F3) and RSDNLRA (F2), for all selections of the binding finger. This mapping is $(r_m, p_r) = map_P(a_m, p_a)$ where $m$ is the position in the alignment (residues 47-53 for F2 and 75-81 for F3), $p_r = m$, the position in the template, $p_a$, depends on the template used to model Zif268 and $a_m$ is the corresponding residue in the template in position $m$ of the alignment. From the template structure we extract the contacts (*triads, etriads* and *ftriads*) between amino-acids and dinucleotides and generate the statistical potentials. However, this introduces a bias by overestimating the constant amino-acids and nucleotides that have not been modified in the experiment. Therefore, only the triads affecting the amino-acids and nucleotides under test are considered to generate the potentials. This is, we only consider the contacts between the 6 amino-acids labelled by "X" and dinucleotides containing one of the 3bp labelled by "N" to generate two potentials, one for F2 and another for F3 finger positions. We restrict each set of Zif268 sequences to those with highest signal of the B1H binding experiment in order to obtain potentials more specific or associated with the strongest binding. We define two thresholds based on the affinity percentile of a sequence: 1) higher than 80%; and 2) higher than 50%. To calculate the affinity percentile of a sequence we follow the same definition as the authors (14). Each sequence in its corresponding domain has a logged and normalized frequency of its observation. Hence, the affinity percentile is defined as the sum of all other frequencies lower or equal to the frequency of the sequence (e.g. an affinity percentile of 80% implies that the sequence is on the tail with highest number of observations, around the top 20%). However, the number of selected sequences may be too different between experiments (i.e. the 3bp binding TGA may have 30 sequences with affinity percentile higher than 80, while AAA has only 2), which produces the opposite bias on the expected potential. To avoid a bias on the number of sequences selected, we force to have around 500 sequences for all 3bp experiments, by repeating as many times as we need each sequence (e.g. if only 2 sequences are selected for AAA and they are equally representative, we should repeat 250 times each). Each sequence is then repeated $500 \times \frac{p_{seq}}{\sum_{k \in A_{80}} p_k}$, where $A_{80}$ is the set of sequences with affinity percentile higher than 80 (or we use $A_{50}$ for affinity percentile higher than 50). As a consequence of the approach, the contacts derived from the B1H experiment are limited to relatively short distances (the largest contacts are around 15-20Å). However, we note that we also use contacts extracted from other structures of the C2H2-ZF family in the PDB and from the use of PBM experiments, covering larger distances up to 30 Å.

### 10. Prediction of the PWM using the sequence of a TF

Let be a TF, named target TF in the set CIS-BP and with known PWM. We do a blind test assuming this PWM is unknown and develop two prediction approaches using the rest of TFs in the database, their sequences and their PWMs: 1) the first approach is straight-forward based on sequence similarity; 2) the second approach uses the enrichment of predictions. Both approaches are explained below.

#### a. Straight-forward prediction

The straight-forward prediction is based on sequence similarity. We use MMSeqs2 (5) to search sequences of TFs in CIS-BP that align with the sequence of target TF with sufficient percentage of identical residues. We classify the matches of this search by percentage of identical residues aligned (ID). Then, we analyze the relevance of sequence similarity on the prediction of the PWM. The group of TFs classified as "$ID_a$" is the set of TFs from CIS-BP which sequence aligns with the sequence of target TF with a percentage of identic residues (ID) around $ID_a$ in an interval of 10% (i.e. $ID_a > ID \geq (ID_a - 10)$). Let's consider the group "$ID_a$", the **straight-forward prediction** assumes that the PWM of target TF is the PWM of the TF with higher ID in the group. We define as **solutions at $\boldsymbol{ID_a}$** all PWMs of TFs aligned with target TF with ID around $ID_a$and define as **straight-forward prediction at $\boldsymbol{ID_a}$** the PWM of the TF with higher ID. We also group TF targets by ID, where a TF target belongs to a group $ID_a$ if it has one or more solutions of TFs in group $ID_a$.

To evaluate the quality of the prediction, we compare the PWM of target TF with the PWM predicted. We use TOMTOM, from MEME suite (10), to compare two PWMs and define the **score of similarity** as:

$$score_{similarity} = -\log(P_{value})$$

(eq.43)

Where $P_{value}$ is the significance of the TOMTOM comparison of both PWMs. We calculate the distribution of scores of similarities for all solutions of TF targets of CIS-BP as a function of the percentage of identical residues (i.e. grouped by ID). To analyze the results of several solutions within the same interval of ID, we calculate the maximum, the minimum and the average score of similarity of all solutions of TF targets as a function of ID.

We also show the quality of the prediction by ranking the score of similarity between the prediction and all PWMs in CIS-BP. Then, we define the **score of ranking** by the position in the ranking of the score of similarity with the PWM of the target:

$$score_{ranking} = (size - rank + 1)$$

(eq. 44)

Where "size" is the total number of TFs with known PWMs in CIS-BP and "rank" is the ranking of the score of similarity between the prediction and the PWM of target TF. If the $P_{value}$ between the PWM of the target and the predicted PWM is not significant (i.e. $P_{value} > 0.05$ ), then the score of ranking is null. We calculate the distribution of scores of ranking for all solutions of TFs of CIS-BP as a function of ID. As for the scores of similarities, we also calculate the maximum, minimum and average of all solutions of each target TF, grouped by ID.

We use the score of ranking to calculate the accuracy of the prediction at different values of ID. We use two criteria to consider a prediction **successful**: 1) the score of ranking is $size$ (i.e. the predicted PWM is the most similar to the PWM of the target); and 2) the score of ranking is among a top threshold, allowing for acceptable errors or the incidence of other PWMs similar to the PWM of the target (e.g. top 1% means a ranking score higher or equal to 99). The **accuracy of the prediction** is calculated as the ratio of successful predictions among the total number of TFs with a prediction. Hence, the accuracy is calculated as a function of ID as the ratio of successful predictions of TFs over all TFs with at least one solution at a given ID (i.e. the TF targets belonging in set ID).

The goal of predicting a PWM is to find the potential binding site(s) of the target TF in a DNA sequence. Thus, the straight-forward prediction applies the predicted PWM of a target TF. This is the PWM of the TF which sequence is aligned with the sequence of the target showing the highest ID. We scan a DNA sequence with the selected PWM and collect the potential bindings detected with FIMO. Still, for the sake of analysis or comparison, we test the solutions (i.e. in general more than one PWM per target) for different intervals of ID.

#### b. Prediction by enrichment

The prediction by enrichment uses the search of similar TFs of target TF as in the previous approach and it classifies the matches by percentages of identical residues aligned (ID). Then, for an interval of ID, i.e. $ID_a$, we rank the matches as before, but instead of selecting the top ranked solution with higher ID we collect the PWMs of all solutions at $ID_a$. We calculate the score of ranking as before but for all solutions at $ID_a$ and select the PWMs of the top ranked scores over a threshold (i.e. top ranked PWMs) for each solution. We have to note that we only select those for which the score of similarity is significant (i.e. $score_{similarity} > -\log(0.05)$). By definition, the number of selected PWMs is smaller than the total number of TFs with known PWMs in CIS-BP. Consequently, some PWMs are selected more often than others. We define the **enrichment** of a PWM as the percentage of the number of times the PWM has been selected:

$$enrichment(PWM) = 100\frac{n_{times}(PWM)}{n_{solutions}}$$

(eq. 45)

Where $n_{times}(PWM)$ is the number of times PWM has been selected by any of the solutions of target TF and $n_{solutions}$ is the total number of solutions of the target TF at $ID_a$. By its definition, $n_{times}(PWM) \leq n_{solutions} \leq size$ . If the threshold of top ranked scores is as large as the whole dataset of PWMs (i.e. "size"), the enrichment would be 100 for all PWMs. Therefore, it's clear that we need to limit this threshold by the probability to obtain an enrichment by random, otherwise the value of enrichment is meaningless.

Let T be the threshold of top ranked scores to calculate the enrichment. Thus, we collect a total of $T \times n_{solutions}$ PWMs and each can be selected a maximum of $n_{solutions}$ times. Let "size" be the total number of TFs with known PWMs in CIS-BP as before; then, the random probability to select one of them is $1/size$ while the probability to obtain a PWM in a group of T selected PWMs is $T/size$. Let E be the enrichment of one PWM, meaning that we select $n_{times} = n_{solutions}\frac{E}{100}$ this PWM. Then, the probability ($\rho$) to select $n_{times}$ a PWM out of $n_{solutions}$ when we have selected T PWMs is obtained by the binomial distribution:

$$\rho = \binom{n_{solutions}}{n_{times}}\left(\frac{T}{size}\right)^{n_{times}}\left(1-\frac{T}{size}\right)^{n_{solutions}-n_{times}}$$

(eq. 46)

We then require that the enrichment should be larger than the enrichment at which $\rho$ is higher than $1/size$, and this is obtained by substituting $n_{times}$ by $n_{solutions}\frac{E}{100}$ in equation 46, otherwise we define it null (i.e. $enrichment = 0$).

We calculate as a function of ID the enrichment of the PWM of each target TF with at least one solution at such ID. This can be plotted to help us evaluating the quality of the enrichment on the prediction of PWMs. Enrichment values vary between 0 and 100, but we have to note that: 1) if none of the solutions produces a significant score, except its own PWM, the enrichment is null; 2) when analyzing the enrichment of the PWM of a target TF, if within the number of top selected PWMs the PWM of the target is never selected, the enrichment is null; and 3) by limiting the enrichment as a function of the number of top selected PWMs to those with more significant probability than random, some enrichments are nullified. In conclusion, neglecting all target TFs with null enrichment, the coverage of TFs that can be analyzed is downsized.

To show the accuracy of the prediction we rank the enrichment of solutions of each target TF when they are significant. We define the **score of ranking of enrichment** as in equation 44, using the rank of the enrichment instead of the score of similarity. The score of ranking is undefined if the enrichment is null, as this should affect the coverage, not the accuracy. Then, we calculate the score of ranking of the enrichment of one solution of each TF (target) as a function of ID. When plotting the "score of ranking of enrichment" it is also important to show the **coverage**, as for some TFs either their PWM is not selected among the top or the probability of achieving the corresponding rank of enrichment is not significant. Finally, we use the score of ranking of enrichment as above to calculate the accuracy of the prediction (i.e. we apply the same two criteria to consider a prediction successful).

As before, the goal of predicting a PWM is to find the potential binding site(s) of the target TF in a DNA sequence. However, when applying the prediction based on enrichment, we use all solutions of target TF. We use the solutions to scan with FIMO a DNA sequence and collect the resulting fragments detected. DNA sequence fragments, sometimes overlapping or even the same, are collected as many times as they are detected by the different solutions. Then, we calculate the enrichment for each DNA fragment (or nucleotide) instead of the enrichment of a PWM. The enrichment is calculated with equation 45, using the number of times the fragment (or nucleotide) has been collected over the total number of solutions. Due to the scanning with FIMO, the same nucleotide can be collected more than once by different PWMs overlapping around the same region. Therefore, it is more convenient to plot the enrichment by nucleotide along the DNA sequence (i.e. number of times a nucleotide is collected over $n_{solutions}$) than by fragments (this type of plot is defined as **nucleotide profile**). We also limit the enrichment by the significance of the matches of FIMO, controlling the probability of the match (using the P-value obtained by FIMO) to filter out unreliable results and ensure the quality of the prediction. As for straight-forward predictions, we can limit the number of solutions by the percentage of identical residues (ID), in order to have an additional control of the enrichment, or to analyze the quality of results and compare methods of prediction.

### 11. Scoring TF-DNA binding with structure

#### a. Scores of single domain structures

Given the structure of a protein-DNA complex, either experimentally obtained (i.e. from crystallography and identified by a PDB code) or modelled (i.e. as described in section 8), we define several scores of the interaction based on statistical potentials. First, we calculate the interface of the interaction and extract all triads, extended-triads and feature-triads associated with distances shorter than 30Å, as defined in section 3. Then, the score of the interaction is defined as the sum of the scores (i.e. potential) of all triads with their associated distances (or extended-triads or feature-triads, depending on the type of score). The same approach is applied for z-scores. Let $Scr$ be a potential as defined in section 4, or a z-score as defined in section 5. Let C be the set of triads (extended-triads or feature-triads, depending on the definition of $Scr$) and their associated distances (d), amino-acid residue number (p) and dinucleotide position (q). The score of the interaction is defined as:

$$score_{Scr} = \sum_{(triad,d,p,q)\in C} Scr(f_{td}(triad,d,p,q))$$

(eq.47)

We can obtain the score of a TF without knowing the structure of the TF-DNA binary complex if it can be modelled. We use the structure of a template to generate the set of triads and the mapping of amino-acids derived from a sequence alignment, mapP as defined in section 9 (eq.38), between the TF sequence and the sequence of the template. We also need the mapping of dinucleotides between the DNA sequence we wish to model and the DNA sequence in the template interface (see section 8.c). Instead of modeling the structure of the TF-DNA complex, we modify the scores in equation 47 by applying the substitution of the corresponding amino-acids, using the functions defined as in section 3, $\varepsilon_a(triad, r)$ and $\eta_v(triad, n)$, and the mappings mapP (in eq. 38) and mapD (in eq. 31) between the templates and the sequences of TF and DNA, respectively. Here, instead of using simplified mappings ($\widehat{map}_D$ and $\widehat{map}_P$ from equations 37 and 39), we generalize the formula by defining special functions, $f_1$ and $f_2$, to extract the dinucleotide or amino-acid positions in the DNA or protein sequences:

$$f_1(r,q) = r$$
$$f_2(r,q) = q$$

(eq. 48)

Where $r$ is either a dinucleotide or an amino-acid residue, and $q$ is a position of a dinucleotide or an amino-acid, respectively for DNA or protein sequences. Then we calculate the score of the interaction as:

$$score_{Scr} = \sum_{(triad,d,p,q)\in C} Scr(\eta_v(\varepsilon_a(triad, f_1(map_P(a,p))), f_1(map_D(v,q))), d)$$

(eq.49)

**b. Multiple domain TFs**

There are TF structures with more than one domain, where each domain uses a different potential to evaluate the interaction (i.e. when using family potentials and domains belong to different families). One example is the particular case of the C2H2-ZF family, where the TF has several domains like F2 (internal) and two extreme domains, at the N-tail (F1) and C-tail (F3). For TFs of the C2H2-ZF family we generate two potentials for finger domains in F2 and fingers in F3, then we will use the F2 potential for F1 too. The application is straight forward as the score in equation 49 is the sum of the scores of triads; therefore, we apply equation 49 and each triad is calculated with the potential of the domain corresponding to the amino-acid position of the triad. However, we have to note that this approach is limited to the use of normalized scores, such as z-scores, to avoid the combination of different scales. To calculate the score, we require an input assigning the domain for each amino-acid position (i.e. for the triad) or split the structure of the TF in its domains, calculate the score of each domain separately and then sum. In general, for any TF we split the sequence in domains (e.g. using PFAM domains) and model the structure of the complex for each individual domain. For the particular case of the C2H2-ZF family, we have to manually split the domains in first and inner domains (using the potential of F2) and the last domain (using the potential of F3).

### 12. Construction of PWMs using TF structures

Given the structure of a protein-DNA complex, either experimentally obtained (i.e. from crystallography and identified by a PDB code) or modelled (see section 8), we obtain the PWM of the TF by means of statistical potentials, using scores or zscores, calculated with the complex structure (we use the zscore of $ES3DC_{dd}$ as example). Let be $length$ the number of nucleotides of the DNA sequence in the complex. We use a sliding window of 8 nucleotides to generate fragments of 8 continuous nucleotides. Fragment "k" is defined as the interval of nucleotides [k,k+7], where k ranges from 1 to $length - 7$. If the length of the DNA sequence is shorter or equal to 8, we use only one fragment defined as the DNA sequence itself. For each fragment "k" of the DNA sequence in the complex structure ($DNA_k$), we collect the set of triads, extended-triads and feature-triads with their associated distances between protein and DNA at less than 30Å and the associated amino-acid and dinucleotide positions (i.e. $C_k,\ eC_k, fC_k$, respectively), where the dinucleotide in the triad belongs in fragment "k". We remind the substitution function $\eta_v(etriad, n)$ from section 3, to substitute the dinucleotide $v$ of an $etriad$ by the dinucleotide $n$. Similar functions are defined to substitute the dinucleotide in triads and feature-triads (these are only affected in the change of the nitrogenous bases).

Then, for each fragment $DNA_k$, we obtain all possible DNA sequences with the length of the fragment (i.e. for a length of 8 residues this is $4^8$) forming a set, named set $F_k$. We define a mapping $map_{DNA_k}$ for the sequence of $DNA_k$ between sequence position and dinucleotides (i.e. $map_{DNA_k}(j) = \omega_j\omega_{j+1}$ with $\omega_j$ nucleotide in position $j$ of $DNA_k$). We also define a mapping $map_{seq}$ for any sequence $seq$ in $F_k$ between sequence position and dinucleotides (i.e. $map_{seq}(j) = \omega_j\omega_{j+1}$ with $\omega_j$ nucleotide in position $j$ of $seq$). We calculate the score of any sequence on $F_k$ with the set of triads, extended-triads and feature-triads, using the associated distances, residue number and dinucleotide positions from the complex structure (i.e. triads, extended-triads and feature-triads as $C_k,\ eC_k, fC_k$, respectively) and using the corresponding substitution function (e.g. $\eta_v(etriad, n)$) as defined in section 3. Let $Scr$ be the score of application and assume we apply it on extended-triads without loss of generality, then the score of a sequence $seq$ in $F_k$ is:

$$score_{seq} = \sum_{(etriad,d,p,q)\in eC_k} Scr(\eta_v(etriad,n),d,p,q)$$

(eq. 50)

Where, for each $(etriad,d,p,q) \in eC_k$, $etriad,d,v$ and $n$ are calculated using the functions as defined in section 3 and the mappings defined above:

$$etriad = f_t(etriad,d,p,q)$$
$$d = f_d(etriad,d,p,q)$$
$$q = f_n(etriad,d,p,q)$$
$$v = map_{DNA_k}(q)$$
$$n = map_{seq}(q)$$

We normalize the scores of sequences in $F_k$ between 0 and 1, by transforming the score:

$$normal(score_{seq}) = \frac{score_{seq} - \min(\{score_{seq};\forall seq \in F_k\})}{\max(\{score_{seq};\forall seq \in F_k\}) - \min(\{score_{seq};\forall seq \in F_k\})}$$

(eq.51)

Then, we rank the normalized scores and select only the DNA sequences producing the top scores over a **cut-off threshold**. The cut-off is specific for the family of the TF (**family threshold for PWM**) if the scores are calculated with the family-specific potential, otherwise it is a general threshold (**general threshold for PWM**) as the scores are calculated with the general potential (see above in section 7). These thresholds are also dependent on the use of PBM data and Taylor's approach.

The selected sequences correspond to a fragment that may be only part of the original sequence length (as taken from the complex). Therefore, we complete all fragments with dummy nucleotides at 5' and 3'. Naming the dummy nucleotide as "N", the sequence with the original length ($length$) derived of a sequence ($seq$) selected from $F_k$ is defined as $full(seq)$, where:

$$full(seq) = N_{[1,k-1]} + seq + N_{[k+8,length]}$$

(eq. 52)

We proceed similarly for all fragments and obtain a multiple sequence alignment with all selected extended sequences, all with the same length and without gaps. Finally, in order to calculate the percentage of each nucleotide in each position of the alignment, we neglect the dummy nucleotides (N) and we use the percentages to define the PWM.

Following the approach in section 11, we also obtain the PWM of a TF without knowing the structure if this can be modelled. We require only the structure of a template to generate the set of triads and the mapping of amino-acids derived from a sequence alignment, $map_P$ as defined in equation 38 (sections 9 and 11), between the TF sequence and the sequence of the template (we assume a simplified mapping, $\widehat{map}_P$, without loss of generality). We don't need to model the structure of the TF-DNA complex, we only need to modify the scores in equation 50 by applying the substitution of the corresponding amino-acids, using the function defined in section 3, $\varepsilon_a(etriad, r)$, and the mapping, $\widehat{map}_P$, and preserving the rest of definitions as in equation 50:

$$score_{seq} = \sum_{(etriad,d,p,q)\in eC_k} Scr\left(\eta_v\left(\varepsilon_a\left(etriad, \widehat{map}_P(a)\right), n\right), d, p, q\right)$$

(eq. 53)

The rest of the approach follows the same procedure for the construction of the PWM up to the full DNA sequence.

### 13. Predicting the PWM with the structure of a TF

Let be a TF from the set CIS-BP with its PWM already assigned, named target TF. As in section 10 (see above), we assume the PWM of such TF to be unknown and develop two approaches to predict it. Conversely to the prediction by sequence, we use the structural models of the TF in complex with DNA. We use as above two different approaches: 1) the first approach is straight-forward, based on the use of one single model obtained with one template structure; 2) the second approach uses the enrichment of predictions using several conformations (either obtained with different templates or the same). Both approaches are explained below. For the sake of analysis, we classify all target TF sequences from CIS-BP in in groups of ID as in section 10 and obtain the predictions for each of them using their structural model.

#### a. Straight-forward prediction

The straight-forward prediction is based on the selection of one single model as the structure of TF bound with DNA. The straight-forward option is to select as template the structure closest to target TF (i.e. with the highest percentage of identical residues in the alignment between both sequences). Then, we model the structure of the complex (see section 8) and calculate the PWM (see section 12). However, in order to analyze the results with several models, we obtain $n_{models}$ models (i.e. $n_{models} = 100$) for each template, producing hundreds of conformations of the same TF and several PWMs that we name as in section 10, **solutions**. We evaluate the quality of the prediction using TOMTOM to compare the solutions with the PWM of target TF. We define the **score of similarity** as in section 10 (i.e. $score_{similarity} = -\log(P_{value})$). Then, we calculate the distribution of scores for all TFs with known PWM in CIS-BP that can be modelled as a function of ID (i.e. with target TFs grouped by ID as in section 10, using the criteria explained on section 8 for modelling). To analyze the results, we calculate the maximum, the minimum and the average of the scores of similarities of all solutions of each target TF.

We define the **score of ranking** by the position in the ranking of the score of similarity with the PWM of the target using equation 44 (in section 10). We calculate the distribution of scores of ranking for all solutions of TFs of CIS-BP as a function of the

ID. We also calculate the maximum, minimum and average of ranking of all solutions of each target TF, grouped by ID. We use the score of ranking to calculate the accuracy of the prediction with the same criteria of **success** as in section 10. However, as we accept several solutions with the same template for the analysis, we use the score of ranking among a top threshold to calculate the ratio of success (i.e. if at least one of the solutions among the top selected is correct the prediction of target TF is successful). The accuracy of the prediction is calculated as the ratio of successful predictions among the total number of TFs with a prediction. The accuracy is calculated as a function of ID as the ratio of successful predictions of TFs over all TFs that were used for the prediction by sequence at the corresponding ID (i.e. grouped as in section 10).

The goal of predicting a PWM is to find the potential binding site(s) of the target TF in a DNA sequence. Consequently, the approach to apply the straight-forward prediction uses the predicted PWM of a target TF using one single model (i.e. obtained with the closest template). We use the PWM to scan a DNA sequence and collect the potential bindings detected with FIMO.

#### b. Prediction by enrichment

As in section 10, the prediction by enrichment uses a few selected PWMs of a target TF for the prediction. We calculate the score of ranking as before, i.e. for each target we obtain several models and with each model a PWM, named solution. The solutions are compared with the PWMs of CIS-BP. Then, we select the PWMs of CIS-BP with the top ranked scores over a threshold (i.e. T top ranked PWMs as in section 10) for each solution. The **enrichment** is defined as in equation 45, where $n_{solutions}$ is the number of models generated for target TF ($n_{models}$ times the number of templates). We calculate the probability ($\rho$) to select $n_{times}$ a PWM out of $n_{solutions}$ when we have selected T PWMs using equation 46. As in section 10, we define null the enrichment if $\rho$ is higher than $1/size$, being $size$ the total number of TFs in CIS-BP with known PWM and modelled (or known) structure (using the criteria explained on section 8 for modelling). We calculate the distribution of enrichment as a function of ID as before (i.e. with target TFs grouped by ID as defined in section 10).

We calculate the accuracy of the prediction by ranking the enrichment of solutions of each target TF as in section 10. The **score of ranking of enrichment** is defined with equation 44, using the rank of the enrichment instead of the score of similarity. We calculate the score of ranking of the enrichment and coverage of the prediction of each target TF as a function of ID, following the same approach as in section 10. Then, we use the score of ranking of enrichment to calculate the accuracy of the prediction.

As in section 10, we use all solutions of a target TF to predict the binding site in a DNA sequence. We use FIMO to scan the DNA sequence with each solution and collect the resulting fragments (it has to be noted that fragments are collected as many times as they are detected by the different solutions). Then, we calculate the enrichment for each DNA fragment (or nucleotide) using the number of times the fragment (or nucleotide) has been collected over the total number of solutions. We calculate and plot the enrichment by nucleotide along the DNA sequence (i.e. a **nucleotide profile**) as in section 10. We limit the enrichment by the significance of the matches of FIMO to filter out unreliable results and ensure the quality of the prediction.

#### c. Comparison of different sets and methods to predict PWMs

We note that the comparison between results of predicting PWMs of TF targets by means of structures and by sequences is not straight forward. The quality of the PWMs predicted by sequence is determined by the PBM experiments, while for the prediction with structural models there is no experimental information. Therefore, the scores of similarities of different solutions have dissimilar ranges: while for the sequence approach the scores of similarities may reach values of 100, the solutions by structural models never reach values higher than 10. In order to compare the results of these different approaches we need to normalize them. Given a score of similarity or a score of ranking, $score$, we define the normalized score as in equation 51 but scaling between 0 and 100:

$$normal(score) = 100 \times \frac{(score - min)}{(max - min)}$$
(eq. 54)

Where $min$ and $max$ are respectively the minimum and maximum values of the scores obtained with all solutions of TF targets.

The analysis and comparison of approaches can also be done by groups of TFs in families. TF targets are grouped according to their family, as defined in CIS-BP. A TF may have more than one DNA binding domain, consequently it may be ascribed to more than one different family. Then, instead of belonging in more than one group, we define the family of the TF as the sum of family names and form a new group. The values of $max$ and $min$ to normalize the scores in the family specific analysis are restricted to the specific set of TF targets, which implies varying the expectation of scores for each particular set. The approach doesn't implies using solutions of a specific family but entails the results of TF targets of a specific family. The different accuracies proof that for some families it may be easier to correctly predict a PWM than others. Therefore, the analysis by families affects the scores of rankings and consequently the accuracy of the prediction per family.

Furthermore, we calculate the average and statistic deviations of scores with the results of families as a function of ID (ID groups of TF targets defined as in section 10), with and without normalization:

$$score_f(ID) = \frac{1}{Card(f)} \sum_{TF \in f} score_{TF}(ID)$$

$$RMSD_f(ID) = \sqrt{\frac{\sum_{TF \in f} \left(score_{TF}(ID) - score_f(ID)\right)^2}{Card(f)}}$$

(eq.55)

Where, $f$ is a family group of TFs, $score_{TF}(ID)$ is any of the scores defined above for a TF target in set ID that is calculated by modelled structures or by sequence (see above and sections 10, respectively). Then, we calculate the accuracies of prediction per family using the scores of ranking (i.e. $accuracy_f$ ) and calculate the average and deviation using all families:

$$\langle accuracy\rangle_{\Gamma}(ID) = \frac{1}{Card(\Gamma)}\sum_{f\in\Gamma} accuracy_f(ID)$$

$$RMSD_{accuracy}(ID) = \sqrt{\frac{\sum_{f\in\Gamma}\left(accuracy_f(ID) - \langle accuracy\rangle_{\Gamma}(ID)\right)^2}{Card(\Gamma)}}$$

(eq.56)

Where $\Gamma$ is the set of all different families grouping the TF-targets.

**14. Optimal conditions to predict PWMs using the structure of a TF (grid search)**

We use the experimentally known PWMs in CIS-BP to optimize the structural-based prediction of PWMs. The definition of the statistical potentials and the conditions for modeling the structure of TF-DNA interactions and obtaining the PWM require some parameters (i.e the selection of cut-off and binned distances on potentials, using a general or a family-specific potential, etc.) that need to be optimized. Thus, we predict the PWM by means of structure for all TFs in CIS-BP that can be modelled (see section 12) and compare the predicted PWMs with the experimental PWMs by means of TOMTOM. We test the following the parameters/conditions:

1) Use potentials derived by PDB data only or adding PBM data.
2) Use general or family specific potentials.
3) Use Taylor's approach or none to infer triads when the amount of data is limited.
4) Use several **family and general thresholds for PWM** as described in section 12. We test values from 0.7 to 1.0 in steps of 0.01 and select the thresholds yielding the best results of the TOMTOM comparison. Among the best results of TOMTOM we select the smallest threshold value.

We select the combination of conditions producing the best score and p-value with TOMTOM and obtain different sets of optimized parameters for all TFs grouped in different TF families as defined in CIS-BP.

The comparison between experimental and predicted PWMs is done using TOMTOM from the MEME suite (see sections 1 and 13). We select the parameter combination producing the largest number of solutions in a family set with a TOMTOM p-value under 0.05 (i.e. $score_{similarity} > -\log(0.05)$ ). If more than one combination can be accepted, we select the combination that produces the maximum average of $score_{similarity}$ in the set of the family.

### 15. Scanning of binding sites and TF clusters along a DNA sequence.

When we introduce a DNA sequence as input ($DNA_{seq}$), we test the capacity of one or several TFs, with known or modeled structure, to bind in one or more binding sites of the DNA. This analysis is performed by scanning and scoring the DNA sequence. We split this analysis in two main parts: 1) scanning and scoring the binding and calculating a binding profile pattern specific of a TF (sections a-c); and 2) selecting several TFs and grouping them in clusters of specific binding regions (sections d and e).

#### a. Scanning of DNA binding domains

The DNA sequence is scanned with the PWMs obtained with all the structures of TFs and protein-DNA interactions collected from the PDB (defined in section 1 as $PDB_{DNA}$). To obtain these PWMs we need to consider that several proteins and TFs interact with a DNA sequence in the form of hetero- and/or homo- complex of proteins forming a quaternary structure. Therefore, for each complex structure of $PDB_{DNA}$ we obtain one or several PWMs as follows: 1) we detect all protein chains interacting with any of the two DNA chains (i.e. strands) forming the double-strand helix; 2) we construct the structures of all combinations of protein-chains that bind the same helix of the original structure; and 3) we calculate the PWMs of these structures and store them associated with the corresponding structure (we name this set as **3D-PWM**). This approach produces PWMs of individual chains and their combinations. For example, given a heterodimer with two protein chains, A and B, we obtain three different PWMs: one for the binding of chain A, another for chain B and another for the heterodimer formed by A and B. In addition, for the web service the user can also upload a specific PWM associated with a TF sequence, then the service checks the closest PWM stored in 3D-PWM associated with a structure and replaces it with the uploaded PWM for the scanning search, hence accommodating our approach to the specific needs of users.

The scanning is performed with the program FIMO, using all PWMs stored in 3D-PWM set, limiting the search by a P-value threshold of significance, or with the specific selection of TFs (potentially allowing in the web service to use the corresponding PWMs uploaded by the user). We use a maximum P-value threshold of 0.05 by default, but this may be increased in order to enlarge the number of potential binding proteins.

#### b. Score per Nucleotide: profiles of a DNA binding site.

We define a **nucleotide profile** as a function on $\mathbb{R}$, $f: \mathbb{N} \longrightarrow \mathbb{R}$, of the nucleotide position in a DNA sequence. We have introduced some examples in sections 10 and 13. Here, we define **score-nucleotide profiles** when the function is obtained with the scores and z-scores developed in sections 4-6 with statistical potentials. Given a TF-DNA complex structure, **3D-TF**, and a distance dependent score, $score$, obtained with statistical potentials (e.g. the smoothed z-score of *ES3DC<sub>dd</sub>*). Let be $j$ a nucleotide position of the DNA sequence in the complex, we define a new score, $Scr_{score}$ in $j$, as:

$$Scr_{score}(j) = \sum_{(etriad,d,p,q)\in eC_j} score(etriad, d)$$

(eq.57)

Here we use extended-triads without loss of generality, although depending on the statistical potential we can use triads or feature triads instead. The set $eC_j$ is the set of *etriads* with all associated distances (i.e. any $d$), and amino-acid residue numbers (i.e. any $p$), where dinucleotide in position $q$ implies that $j$ is the position of one of the nucleotides in the dinucleotide at $q$ (i.e. $q = j \; or \; q = (j-1)$). This new score can be normalized by considering the contribution of the nucleotide to the total score or as the percentage of the contribution of all nucleotides of the DNA sequence. Assuming that the score is the smoothed z-score of $ES3DC_{dd}$ and without loss of generality, the normalized nucleotide profile is:

$$Normal(j) = \frac{\sum_{(etriad,d,p,q)\in eC_j}\left(zscore(ES3DC_{dd}(etriad, d))\right)_{smooth}}{\sum_{j=1}^{length}\sum_{(etriad,d,p,q)\in eC_j}\left(zscore(ES3DC_{dd}(etriad, d))\right)_{smooth}}$$

(eq. 58)

or

$$Normal(j) = \frac{\sum_{(etriad,d,p,q)\in eC_j}\left(zscore(ES3DC_{dd}(etriad, d))\right)_{smooth}}{factor_j \times ZES3DC_{dd}}$$

(eq. 59)

Where $factor_j = 2$ for all $j$ but the extremes at 5’ and 3’, in which is 1, and from equation 47 we write $ZES3DC_{dd}$ as:

$$ZES3DC_{dd} = \sum_{(etriad,d,p,q)\in E} \left(zscore\left(ES3DC_{dd}(etriad, d)\right)\right)_{smooth}$$

(eq. 60)

$E$ is the set of extended-triads (with their associated distances, amino-acid numbers and dinucleotide positions) as defined in section 4 and $length$ is the length of the DNA sequence. The factor of 2 in equation 59 is produced by the fact that each nucleotide is counted twice in $eC_j$, with the exception of the extreme positions in 5’ and 3’ where the nucleotides are only reckoned one time.

The curve of $Scr_{score}(j)$ (raw or normalized) along the positions in the DNA sequence is defined as the **nucleotide profile based on 3D-TF** for the (raw or normalized) potential defined in $score$ (e.g. the smoothed z-score of $ES3DC_{dd}$). Hence, profiles defined upon scores derived from statistical potentials are dependent on the structure of the TF-DNA interaction complex. We have to note that, if the structure of this complex has been modelled, several models may be considered (i.e. we define the set of models of TF-DNA as $MDL_{TF}$). Besides, some models may be obtained using different templates, implying that these models introduce a relevant variability on the conformational space of the TF-DNA interaction. Consequently, several nucleotide profiles of scores are accumulated for the same DNA sequence. We then calculate the average and standard deviation of $Scr_{score}(j)$ for all positions $j$ along the DNA sequence with the nucleotide profiles of score based on each model structure $m$ of the $MDL_{TF}$ set (i.e. described as $\left(Scr_{score}(j)\right)_m$ ), using the following equations:

$$\langle Scr_{score}(j)\rangle = \frac{1}{Card(MDL_{TF})} \sum_{m \in MDL_{TF}} \left(Scr_{score}(j)\right)_m$$

$$RMSD\left(Scr_{score}(j)\right) = \sqrt{\frac{\sum_{m \in MDL_{TF}} \left(\left(Scr_{score}(j)\right)_m - \langle Scr_{score}(j)\rangle\right)^2}{Card(MDL_{TF})}}$$

(eq. 61)

Equations in 61 describe two new nucleotide profile functions. The average function is defined as the **nucleotide profile of $score$** (e.g. the smoothed z-score of *ES3DC<sub>dd</sub>*) and the RMSD defines its margins of error or variability.

As seen in sections 10 and 13 and mentioned above, the nucleotide profile can also be calculated with other "scores" different than those derived by statistical potentials, for example by the enrichment or the number of times that nucleotide *j* is selected by several PWMs assigned to a TF (see above). This extends the definition of score nucleotide profiles to other scores different than those obtained with statistical potentials.

**c. TF profiles along a DNA fragment.**

We define **TF profiles** as nucleotide profiles where the value of the function is obtained with the whole interaction between the TF and DNA that can be associated to a nucleotide position. Let be a DNA sequence ($DNA_{seq}$ ) and a TF with known (or predicted) PWM. We use FIMO to calculate and rank the score ($score_{FIMO}$) and significance (i.e. $P_{value}$) for several positions along the DNA sequence where the significance is acceptable (i.e. $P_{value} < 0.05$). Both, score and significance, are associated with a specific interval of the DNA sequence, $[k,l]_r$, that identifies the nucleotides matching the PWM in a ranking order of quality (identified by r). The position of a nucleotide, $j$ in $[k,l]_r$, can be matched more than once by the same PWM (depending on the limit of significance to accept the matches). Then, considering that position $j$ is matched by PWM $n$ times, and identifying the score and significance of each instance, $i$, respectively as $score_{FIMO}(i)$ and $P_{value}(i)$, we define the following **TF profile based on the PWM**:

$$Scr_{FIMO}(j) = \sum_{i=1}^{n} score_{FIMO}(i)$$
$$Scr_{sig}(j) = \sum_{i=1}^{n} -\log\left(P_{value}(i)\right)$$
(eq. 62)

As for nucleotide profiles, more than one PWM can be predicted for a TF, either by sequence or by structure. Consequently, if $PWM_{TF}$ is the set of PWMs predicted for the TF, we define the **TF profile** of the **score of FIMO** and the **score of significance** (respectively) as:

$$\langle Scr_{FIMO}(j)\rangle = \frac{1}{Card(PWM_{TF})} \sum_{m \in PWM_{TF}} \left(Scr_{FIMO}(j)\right)_m$$
$$\langle Scr_{sig}(j)\rangle = \frac{1}{Card(PWM_{TF})} \sum_{m \in PWM_{TF}} \left(Scr_{sig}(j)\right)_m$$
(eq. 63)

When the prediction is based on the structure of a TF-DNA complex, $PWM_{TF}$ set is the set of PWMs obtained by the models in $MDL_{TF}$ from the previous section.

As above, we also define other two TF profiles for the variability and deviations from the average, RMSD, as:

$$RMSD\left(Scr_{FIMO}(j)\right) = \sqrt{\frac{\sum_{m \in PWM_{TF}} \left(\left(Scr_{FIMO}(j)\right)_m - \langle Scr_{FIMO}(j)\rangle\right)^2}{Card(PWM_{TF})}}$$
$$RMSD\left(Scr_{sig}(j)\right) = \sqrt{\frac{\sum_{m \in PWM_{TF}} \left(\left(Scr_{sig}(j)\right)_m - \langle Scr_{sig}(j)\rangle\right)^2}{Card(PWM_{TF})}}$$
(eq. 64)

Another set of **TF profiles** is obtained by using the scores and z-scores developed in sections 4-6 with statistical potentials for the score of a TF-DNA interaction as in section 11. Let be $DNA_{seq}$ the DNA sequence and $TF_{3D}$ the known (or modelled) structure of the TF with a DNA sequence in a crystal (or the DNA in the template, if $TF_{3D}$ is modelled). We use $TF_{3D}$ to calculate the interface and determine the length, $length_{TF_{3D}}$, of DNA in contact with the TF, as in section 8. We split the sequence in fragments using a sliding window of $length_{TF_{3D}}$ nucleotides. Then, we define a DNA fragment similarly as in section 12, $DNA_{seq}(k)$, of $DNA_{seq}$ in position k, as the fragment of $DNA_{seq}$ in the interval [k,k+$length_{TF_{3D}}$]. We model the TF-DNA complex structure, **3D-TF$_k$**, for any position k along the DNA sequence and calculate the binding score as in section 11 (see eq. 36) for any score or z-score. Let be the example, without loss of generality, of the smoothed z-score of *ES3DC<sub></sub>*$_{dd}$. This example scores the TF-DNA interaction in 3D-TF$_k$ as:

$$ZES3DC_{dd}(k) = \sum_{(etriad,d,p,q)\in E_k} \Big(zscore\big(ES3DC_{dd}(etriad,d)\big)\Big)_{smooth}$$

(eq. 65)

Where, $E_k$ is the set of extended-triads (with their associated distances, amino-acid numbers and dinucleotide positions) for the complex in 3D-TF$_k$.

Let be $j$ a position in DNA$_{seq}$. According to the previous definitions, we calculate a total of 2 times $length_{TF_{3D}}$binding scores containing position $j$, as if the TF was sliding along the DNA sequence from $[j - length_{TF_{3D}}, j]$ to $[j,\ j + length_{TF_{3D}}]$. Then, we define the following **TF profile based on TF$_{3D}$**, using the smoothed z-score of *ES3DC*$_{dd}$, as:

$$Scr_{ZES3DC_{dd}}(j) = F(j) * \sum_{k=j-length_{TF_{3D}}}^{j+length_{TF_{3D}}} ZES3DC_{dd}(k)$$

(eq. 66)

The factor F(j) is used to normalize (i.e. to average) the sum. This is:

$$F(j) = \left(length_{TF_{3D}}\right)^{-1}$$

except for the start and end of the interval, where only counts the number of times the z-scores have been summed, which is less than ($length_{TF_{3D}}$). The score can also be weighted by a binding/non-binding weight (1 or 0, respectively). We will consider that the TF will bind DNA$_{seq}$(k) if $P_{value}(k)$<0.001, with the Pvalue obtained by FIMO using the PWM of the TF, matching the fragment DNA$_{seq}$(k). The threshold of significance (often defined as 0.001) can also be selected (i.e. taking values 0.5, 0.1, 0.05, 0.01 or 0.001). Then, eq 66 becomes:

$$Scr_{ZES3DC_{dd}}(j) = F(j) * \sum_{k=j-length_{TF_{3D}}}^{j+length_{TF_{3D}}} \omega(k) * ZES3DC_{dd}(k)$$

(eq. 67)

Where $\omega(k)$ is 0 if FIMO $P_{value}$ is not significant and 1 if the probability to bind is significant (according to the selected threshold).

We normalize the score by scaling it between 0 and 1, being 1 the best score and 0 the worst. This may imply a change on the original sign of the score. For the normalization we require to calculate the scores with the best and the worst interface DNA sequences. The best score is obtained by testing the scores of the DNA sequences considering the most probable nucleotides in each position, according to the PWM obtained with the structure of the TF-DNA complex (see section 12). Using $ZES3DC_{dd}$ in equations 47 and 49 without loss of generality, the best score is the minimum of the values tested and we write it as $ZES3DC_{dd}(min)$. Similarly, for the worst score we test the values of this function on several DNA sequences considering the less probable nucleotides in each position and we take the maximum, $ZES3DC_{dd}(max)$. Then, we define the **normalized TF profile based on TF$_{3D}$** as:

$$Normal_{ZES3DC_{dd}}(j) = \frac{\sum_{k=j-length_{TF_{3D}}}^{j+length_{TF_{3D}}} (ZES3DC_{dd}(k) - ZES3DC_{dd}(max))}{length_{TF_{3D}} \times (ZES3DC_{dd}(min) - ZES3DC_{dd}(max))}$$

(eq.68)

Also, the normalized score can be weighted as in eq 67, using the same weight function to become:

$$Normal_{ZES3DC_{dd}}(j) = \frac{\sum_{k=j-length_{TF_{3D}}}^{j+length_{TF_{3D}}} \omega(k) * (ZES3DC_{dd}(k) - ZES3DC_{dd}(max))}{length_{TF_{3D}} \times (ZES3DC_{dd}(min) - ZES3DC_{dd}(max))}$$

(eq.69)

Another way of normalizing is to consider only the best scoring energy of the DNA$_{seq}$(k) fragment, instead of summing up the scores. This can be defined as:

$$Normal_{ZES3DC_{dd}}(j) = max\left(NSC(k); \forall k \in \left[j - length_{TF_{3D}}, j + length_{TF_{3D}}\right]\right)$$

With

$$NSC(k) = \frac{ZES3DC_{dd}(k) - ZES3DC_{dd}(max)}{ZES3DC_{dd}(min) - ZES3DC_{dd}(max)}$$

(eq.70)

As previously shown, a TF may have more than one structural model or conformation. Following the same approach as before and defining $MDL_{TF}$ as the set of models of the TF-DNA interaction, we define without loss of generality the **TF profile** of the smoothed z-score of *ES3DC$_{dd}$* as:

$$\langle Scr_{ZES3DC_{dd}}(j) \rangle = \frac{1}{Card(MDL_{TF})} \sum_{m \in MDL_{TF}} \left(Scr_{ZES3DC_{dd}}(j)\right)_m$$

(eq. 71)

Where $\left(Scr_{ZES3DC_{dd}}(j)\right)_m$ is the **TF profile based on model $m$** of the smoothed z-score of *ES3DC$_{dd}$*. And the normalized version of eq. 71 is:

$$\langle Normal_{ZES3DC_{dd}}(j)\rangle = \frac{1}{Card(MDL_{TF})}\sum_{m\in MDL_{TF}}\left(Normal_{ZES3DC_{dd}}(j)\right)_m$$

(eq.72)

Following the same approach as in equation 64 we also define the TF profiles for the deviations in equations 71 and 72, respectively as:

$$RMSD\left(Scr_{ZES3DC_{dd}}(j)\right) = \sqrt{\frac{\sum_{m\in MDL_{TF}}\left(\left(Scr_{ZES3DC_{dd}}(j)\right)_m - \langle Scr_{ZES3DC_{dd}}(j)\rangle\right)^2}{Card(MDL_{TF})}}$$

$$RMSD\left(Normal_{ZES3DC_{dd}}(j)\right) = \sqrt{\frac{\sum_{m\in MDL_{TF}}\left(\left(Normal_{ZES3DC_{dd}}(j)\right)_m - \langle Normal_{ZES3DC_{dd}}(j)\rangle\right)^2}{Card(MDL_{TF})}}$$

(eq. 73)

**d. Prediction of TFs that bind a DNA fragment**

Given a DNA sequence as input, our goal is to predict the set of TFs that binds it in a cell or specific species. The first step is to collect all PWMs of TFs with known structure and scan the DNA sequence (see previous section 15.a). This identifies the potential structures and families but not yet all potential sequences. We use BLAST to collect the potential protein homologs of all structures in $PDB_{DNA}$ (see sections 1 and 8) and the corresponding BLAST alignments. Next, we select the proteins and their alignments and construct two mapping functions: 1) the mapping between the protein sequence and the template TF in $PDB_{DNA}$ ($map_P$); and 2) the mapping of dinucleotides in the DNA interface ($map_D$ as defined in section 9, which is obtained with the alignment between the sequence matched by the PWM and the DNA sequence in the template). Instead of modeling the TF-DNA complex, we use the mapping functions to calculate the scores, using equation 36.b in section 11, and rank the potential binding proteins by scores. If the TF belongs to one of the families that may work as homodimer, we calculate the scores and their ranking using the structures of the monomer and the dimer.

**e. Clusters of TFs and complexes of regulatory elements.**

Given a DNA sequence as input, we group the predicted TFs and their binding sites (see previous section 15.d) by the proximity in the DNA sequence (using as cut-off a limiting number of nucleotides, $n_{connect}$). In case a TF acts as a dimer, we force in the same group both monomers of the dimer. The group may contain TFs with overlapping binding sites. This must be allowed because only the disposition of the structure in 3D allows us to determine if the complex can be formed. Furthermore, we add other proteins that interact with any of the TFs binding the DNA sequence. We use the database IntAct of protein-protein interactions (PPIs) to identify the potential partners that interact with the predicted Tfs and consider only those of the same species as the TFs. If two TFs belonging to different groups of TFs (and their bindings) are connected by PPIs (either interactions between TFs or with a common partner) we merge both groups in one. In order to find the connection of TFs by means of PPIs we construct a network in the following steps:

1) We define the DNA binding sites predicted as main nodes
2) We define the TFs as nodes. They are connected to the one or more main nodes if they are predicted to bind them.
3) We add the proteins that interact with the TFs as nodes connected to the corresponding TF partners.

The approach can be continued, adding proteins (nodes) and interactions (edges) that interact with the proteins of the network as it was left in step 3 and then iteratively continue. Two TFs are connected if they interact in the database of PPIs or there is an intermediate protein that interact with both. In the web service the user can select a subnetwork of proteins in this network and model the structure of the binary interactions of TF-DNA and PPI. The ranking of the scores of TF-DNA interactions helps on this selection. Each group of the final set of groups formed by proteins and binding sites is named **cluster**, and the selection of elements (proteins and DNA binding sites) of a cluster in the web service is a potential **TF-complex** or **regulatory complex**.

### 16. Modeling a selected TF-complex of a cluster of close TFs in a DNA fragment

Following the selection of TFs of a cluster, their protein-interacting partners and DNA binding sites from sections 15.d and 15.e, we use the modelling approaches from section 8 to construct each individual binary-complex structure (i.e. PPIs and TF-DNA). To model the structure of their combination forming the TF-complex, we have first to define the expected structure of the fragment of DNA sequence that contains all the binding sites selected from the cluster. We allow for few automated options, either as B-DNA, Z-DNA or curved DNA (as in nucleosomes). The structure of the DNA fragment with the binding sites selected (i.e. a continuous interval from 5' to 3' taken from the original DNA sequence input, $DNA_{seq}$) is modelled with X3DNA according to one of the options (this is named **DNA-frame**). Because the approach for modelling binary interactions (either TF.DNA or PPIs) allows for several potential conformations, it is likely that each PPI and TF-DNA complex has more than one structural model. Therefore, we produce two lists: 1) one list (**TF-DNA list**) with all the combinations of individual TF-DNA binary-complexes; and 2) another list (**PPI list**) with all combinations of conformations of the selected PPIs (the list includes the interactions between TFs and interactions with **co-factors**). The structures in each list are ranked by the maximum length of protein sequences that can be modelled, both for TF-DNA and PPI complexes. Next, for each combination of the TF-DNA list, individual binary-complexes of TF-DNA of all binding sites selected in the list are superimposed on the DNA-frame. The DNA structure of the binary-complexes of TF-DNA are removed after the superimposition, leaving each TF structure bound on the DNA-frame with the corresponding conformation as in the binary-complex (named **DBD-complex** and identified by a **number** from TF-DNA list). Often the conformation of the DNA binding domain of a TF is separated from the structure of other domains. Consequently, with the exception of few cases, it is not possible to combine the lists of PPI structures and each DBD-complex. Thus, in the web service the user can select a specific DBD-complex and one of the combinations of PPIs and handle it for its modeling. Then, the selected set of PPIs and DBD-complex is further processed to obtain the **model of the TF-complex**. We differentiate between the DNA-frame of a nucleosome complex and a linear complex. The nucleosome DNA-frame is formed by the DNA plus the histones of the nucleosome and it models only those TF binding the nucleosome without clashes with the histones. For the model(s) using a linear form of DNA we proceed in two steps: 1) in the first step the DNA is split in fragments of 250 bp or less and modelled as B-DNA and TFs are bound to each fragment; 2) in the last step PPIs are used to identify new binders and model the full complex using IMP with distance restrictions (protein-protein and protein-DNA) identified on each fragment.

**a. Structural model of TF-DNA complexes in fragments of 250bp (or less).**

The DNA sequence is split in fragments with a maximum size of 250bp. Each fragment is modelled with X3DNA in conformation of B-DNA. On the one hand, if the number of DBD-complexes selected from the list is limited or sufficiently small, all conformations are tested by superimposing each DBD-complex on the structure of the DNA fragment. TFs having clashes with other TFs are removed. At the end of the approach one or more combinations of TFs are modelled bound with each DNA fragment. On the other hand, if the number of selected TF-DNA bindings and its many conformations is very large, the number of combinations of TF-DNA conformations for each DNA fragment requires an unfeasible computational time. Then, we proceed with a heuristic approach: once the conformation of a DBD-complex, bound in a specific site, is superimposed on the DNA fragment without clashes with other TFs, the algorithm proceeds with the rest of binding sites until all sites are covered. This approach models the structure of at least one full-complex and it repeats the approach by selecting the model of a new starting binding site and DBD-complex. The number of trials (DBD-complexes been superimposed and tested) is limited to obtain one or more large models of each fragment in a reasonable time.

**b. Linear B-DNA modelled with TF and TcoF.**

**i. Selection of proteins to form the complex and distance restraints**

To obtain a large complex of the whole DNA, we start with the combinations of the structural models of each fragment. For each combination we obtain a list of TFs and the set of amino-acids involved in interfaces of protein-protein and protein-DNA interactions. The list identifies each TF and its conformation (i.e. template-based modeling). The list of proteins is named **protein-list of the complex**. Similarly, for the list of selected PPIs, the interface of each protein is identified and associated with its specific conformation in the PPI model. Next, we select a conformation among the list of PPIs where only one of the partners is in the protein-list of the complex and test the overlap between the interfaces of the common protein (the interface of the interaction with its partner from the list of PPIs and the interface in the interaction of the protein-list of the complex). If the overlap is formed by a minimum number of residues (i.e. 2 amino acids) the PPI is accepted and the protein-interaction partner is incorporated in the protein-list of the complex including (or changing) the interfaces of the new and previous proteins of the list. The procedure is repeated until all proteins of the original protein-list of the complex are tested.

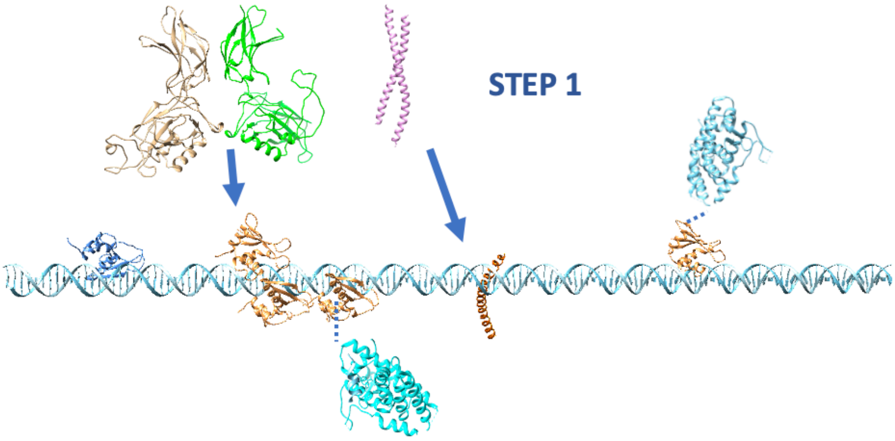


After all proteins in the protein-list of the complex has been tested, we select a conformation among the list of PPIs with both partners in the protein-list and test the overlap between the interfaces of both proteins in both interactions (from the PPI list and the protein-list of the complex). As above, if the overlap is formed by a minimum number of residues (i.e. 3 amino acids) the interaction is accepted. No new partners are included in the protein-list of the complex, but the interfaces of the proteins involved are modified. The procedure is repeated until all proteins of the protein-list of the complex are tested, including its successive modifications.

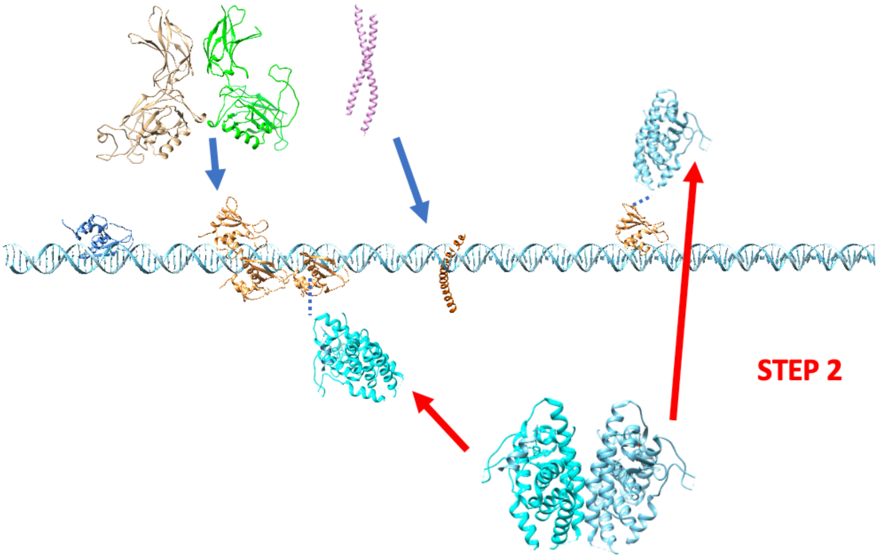

The new protein-list of the complex is used to iterate the approach and to repeat the procedure until no more new proteins are added in the protein-list of the complex, nor the interfaces of the proteins are modified. Then, the interfaces and their associated conformations are used to define a list of distance constraints between the CA atoms of the residues involved in each interface.

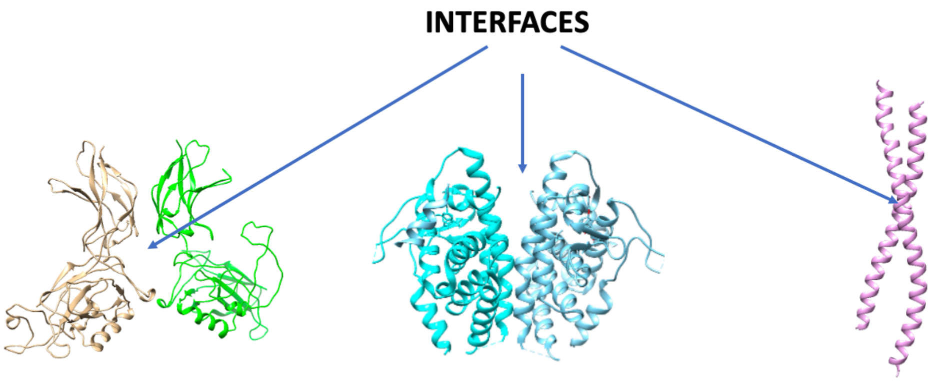


ii. **Integrative Modeling of the complex.**

After the proteins required to form a specific complex have been selected, their interfaces calculated and the distances restraints between CA atoms (or between CA and P atoms for the interaction with nucleotides) extracted, we use the information to generate two files: one for the topology of the complex and another for the distance restraints. The topology is constructed using as “rigid bodies” the whole or partial structures of the proteins from one or several PDB files, while restraints are constructed with the extracted distance-restraints plus a new set defined to maintain the double helix of the DNA. Protein rigid-bodies are connected by beads of 10 amino-acid size allowing for flexibility, but the distance between two rigid-bodies is restricted to less than 15 Å.

The restraints to maintain the DNA double helix are taken from a standard B-form of DNA, using the distances between phosphate atoms of neighbouring nucleotides, up to 5 nucleotides in both directions 5’ and 3’ in the same chain and in the reversed chain (see image). Additional neighbours to calculate distance-restraints can be included and the force constant dampened. The restraint applied will follow a Gaussian behavior, allowing deviations of ±2Å. The restraints on the interfaces (protein-protein and protein-DNA) are taken from the lists of interactors calculated above, using a similar Gaussian formulae. Restraint and topopology files will be used with the Integrative Modelling Platform (IMP) (21) (22) to model the scaffold structure of the full complex.

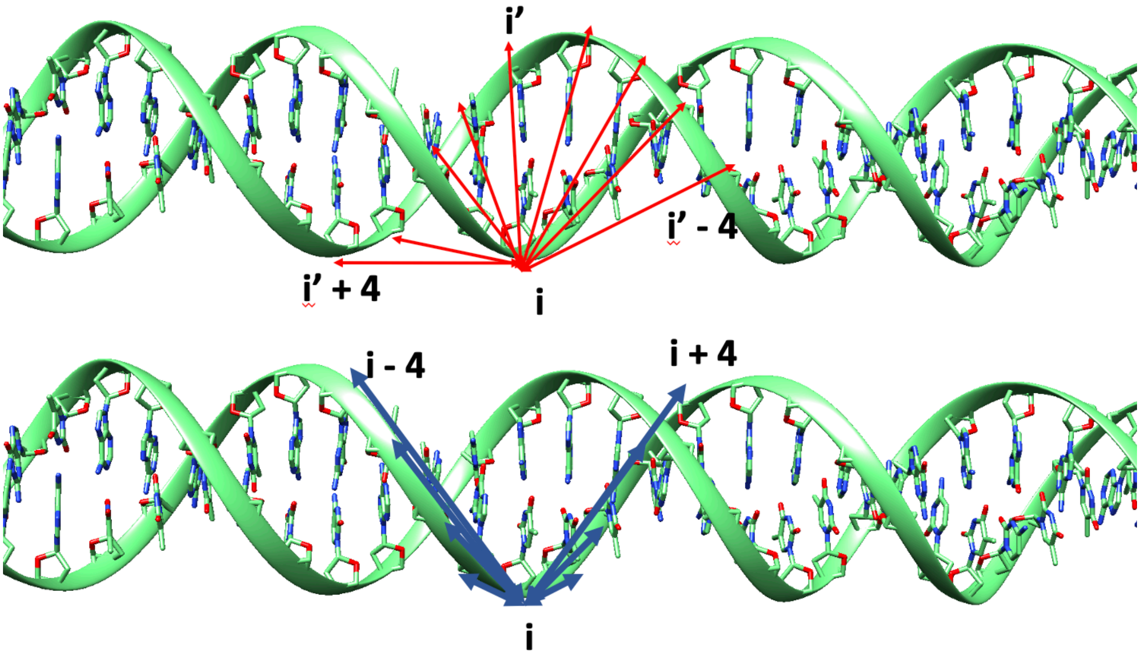
