## Supplementary Table S1 for "Structure-based learning to model complex protein-DNA interactions and transcription-factor co-operativity in *cis*-regulatory elements"

| Uniprot | Family | JASPAR | CisBP | ModCRE |
|---|---|---|---|---|
| Q9SFE4 | AP2 | 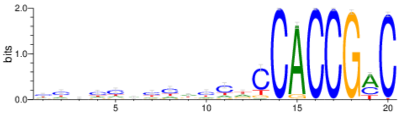 MA1248.1 | 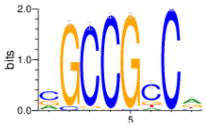 M01615 | 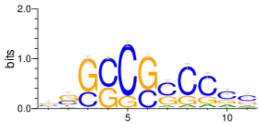 1gcc |
| P61244 | bHLH | 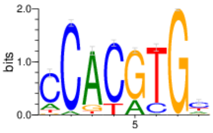 MA0058.3 | 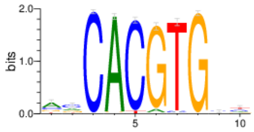 M00938 | 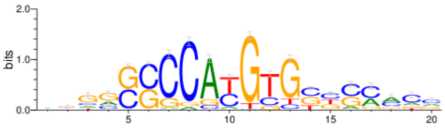 1an4 |
| P17535 | bZIP | 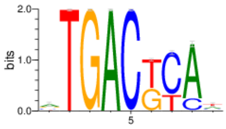 MA1143.1 | 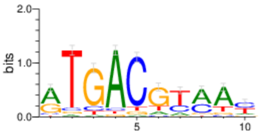 M01001 | 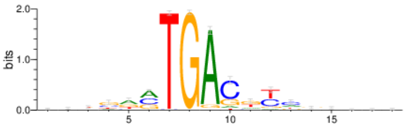 2h7h |
| P22561 | C2H2-ZF | 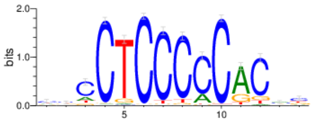 MA1627.1 | 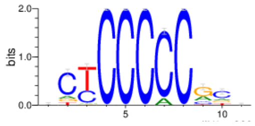 M00249 | 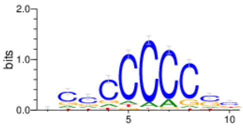 2jp9 |
| Q8IXT2 | DM | 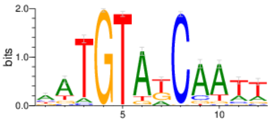 MA1479.1 | 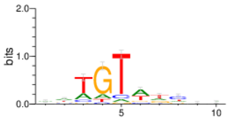 M01933 | 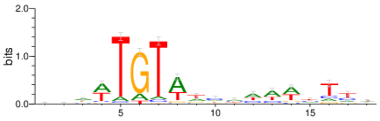 4yj0 |
| P41161 | Ets | 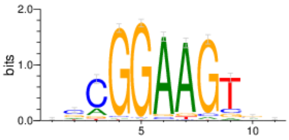 MA0765.2 | 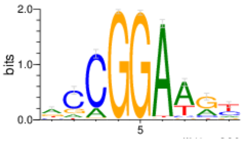 M01485 | 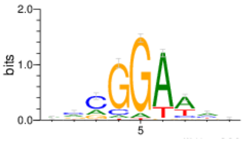 4uuv |
| Q9R1E0 | Forkhead | 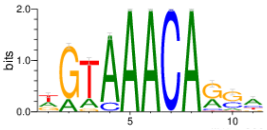 MA0480.1 | 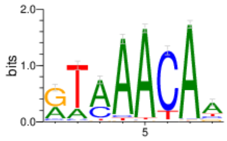 M00797 | 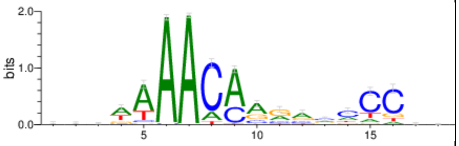 4lg0 |
| Q9NP62 | GATA | 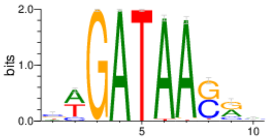 MA0766.1 | 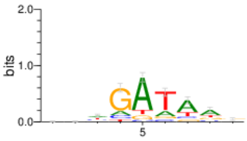 M00167 | 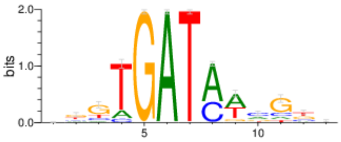 4gat |
| Q9NP62 | GCM | 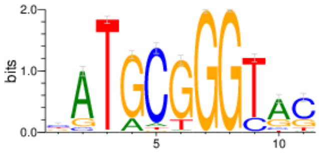 MA0646.1 | 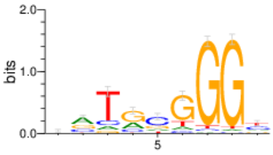 M02020 | 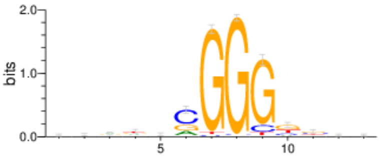 1odh |

| Uniprot | Family | JASPAR | CisBP | ModCRE |
|---|---|---|---|---|
| Q9UBX0 | Homeo-domain | 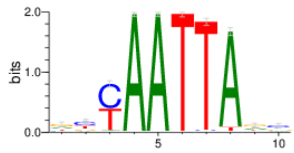<br>MA0894.1 | 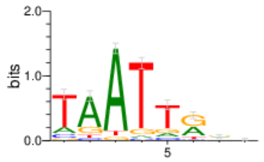<br>M00307 | 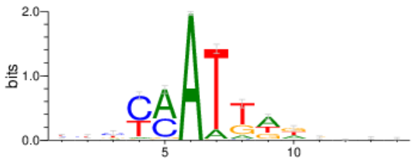<br>3lnq |
| O43316 | Homeo-domain Paired box | 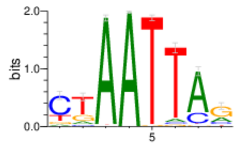<br>MA0068.2 | 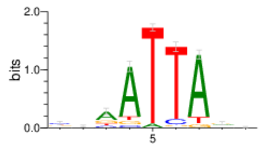<br>M00349 | 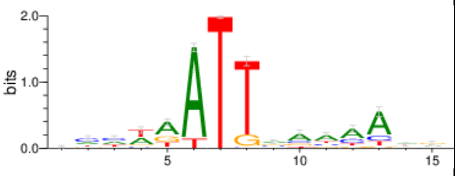<br>5no6 |
| Q14863 | Homeo-domain POU | 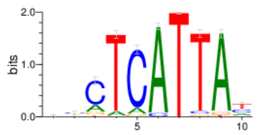<br>MA1549.1 | 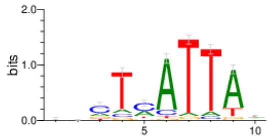<br>M00546 | 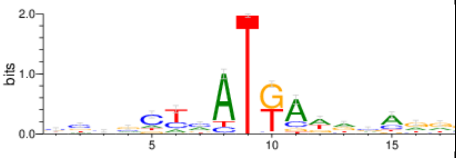<br>2r5y |
| Q03933 | HSF | 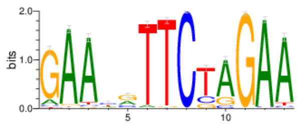<br>MA0770.1 | 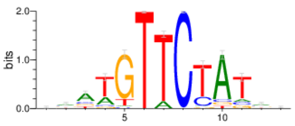<br>M02248 | 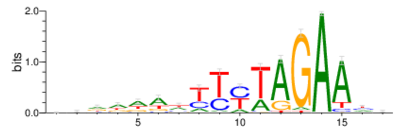<br>5d8l |
| Q92985 | IRF | 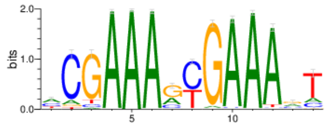<br>MA0772.1 | 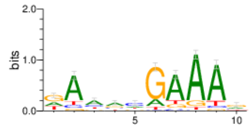<br>M02255 | 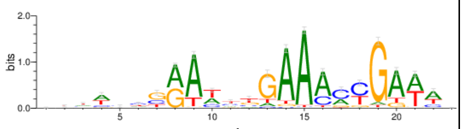<br>1t2k |
| P10243 | Myb/SANT | 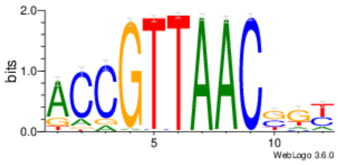<br>MA0776.1 | 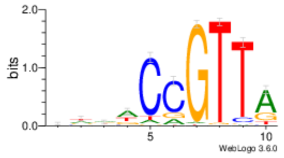<br>M02322 | 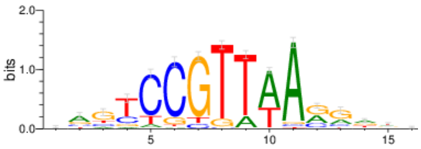<br>1h88 |
| P10276 | Nuclear receptor | 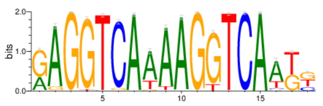<br>MA0729.1 | 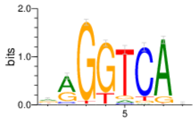<br>M00185 | 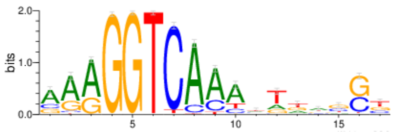<br>1r0n |
| P32114 | Paired box | 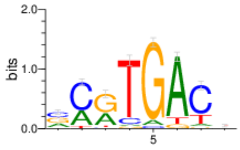<br>MA0067.1 | 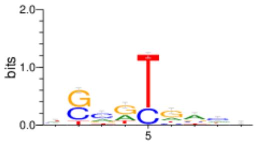<br>M01301 | 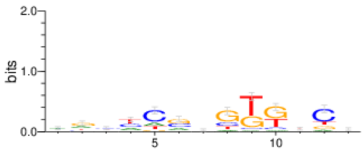<br>1pdn |

| Uniprot | Family | JASPAR | CisBP | ModCRE |
|---|---|---|---|---|
| G1MTN6 | Rel | 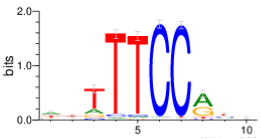 MA0624.1 | 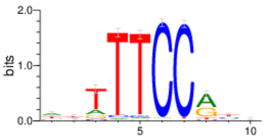 M02443 | 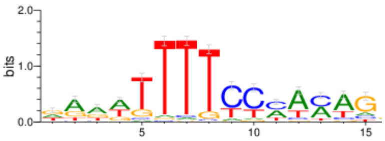 2o93 |
| Q13950 | Runt | 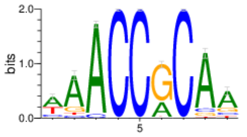 MA0511.2 | 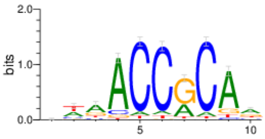 M01307 | 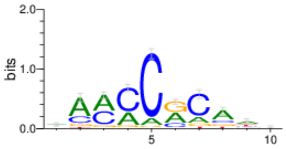 1h9d |
| P40645 | Sox | 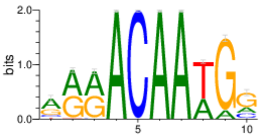 MA0515.1 | 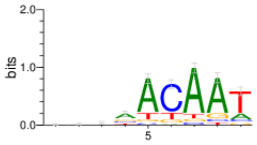 M00828 | 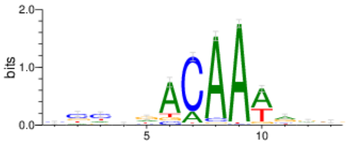 4s2q |
| O15119 | T-box | 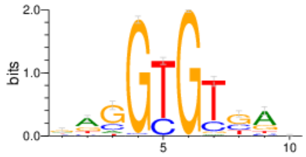 MA1566.1 | 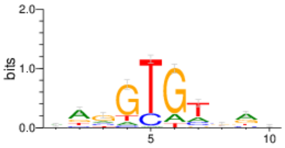 M00834 | 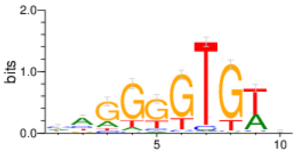 2x6v |
| Q95QD7 | TCR/CxC | 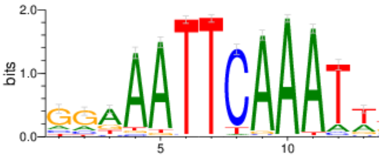 MA1450.1 | 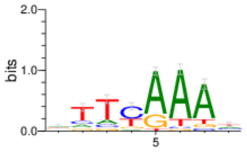 M02526 | 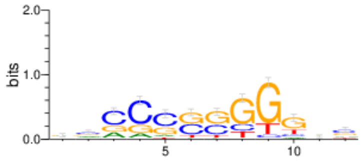 5fd3 |
| Q8VWJ2 | WRKY | 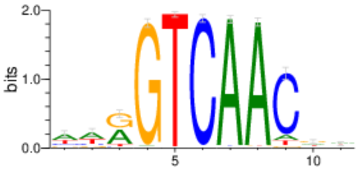 MA1311.2 | 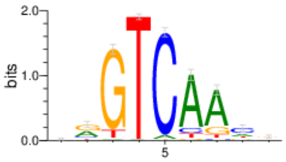 M02564 | 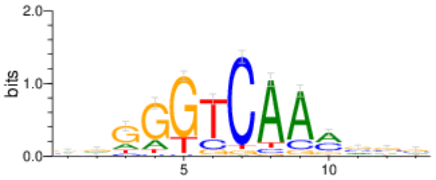 6ir8 |
