## Supplementary figures and images for "Structure-based learning to model complex protein-DNA interactions and transcription-factor co-operativity in *cis*-regulatory elements"

### ABF1_Enriched.png

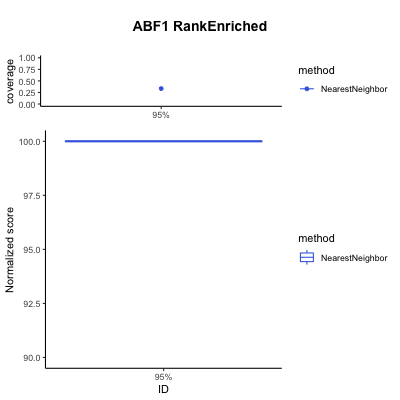

### AP2_Enriched.png

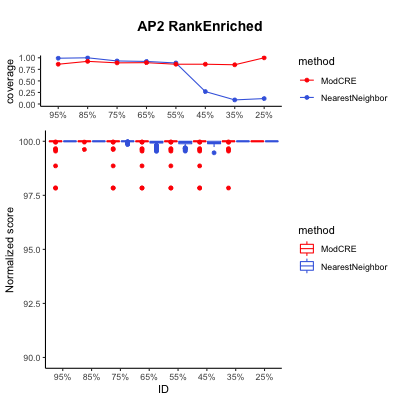

### APSES_Enriched.png

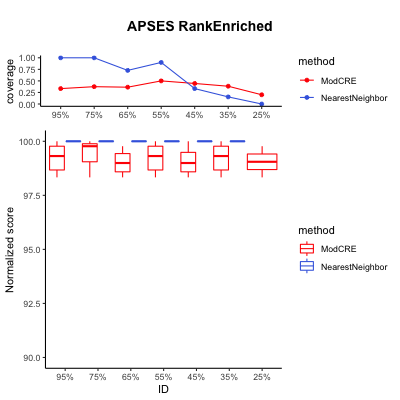

### ARID_BRIGHT_Enriched.png

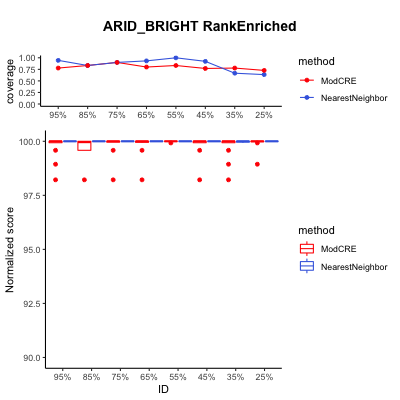

### B3_Enriched.png

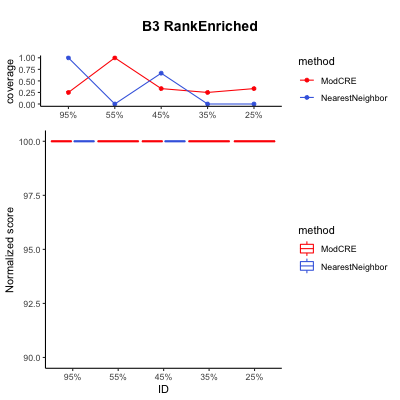

### Brinker_Enriched.png

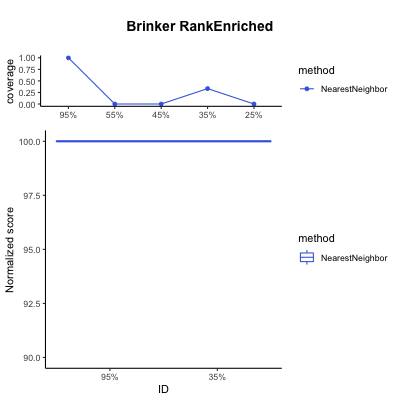

### C2H2_ZF_Enriched.png

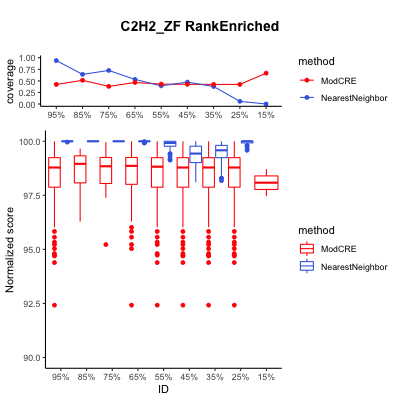

### C2H2_ZF_Homeodomain_Enriched.png

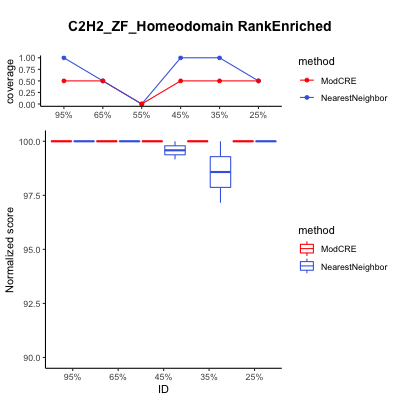

### C2H2_ZF_MADF_Enriched.png

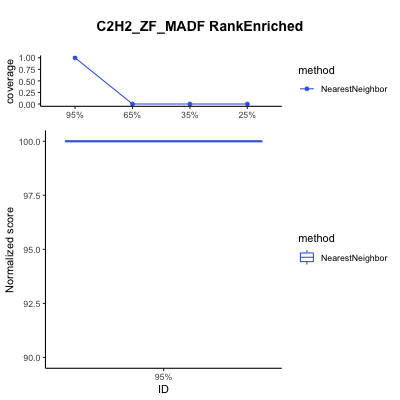

### C2H2_ZF_Zinc_cluster_Enriched.png

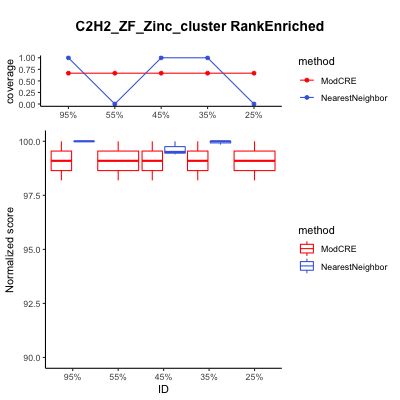

### CCCH_ZF_Enriched.png

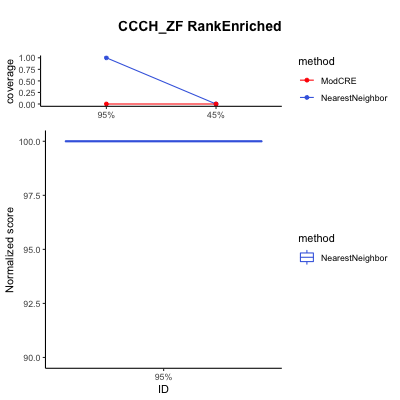

### CSD_Enriched.png

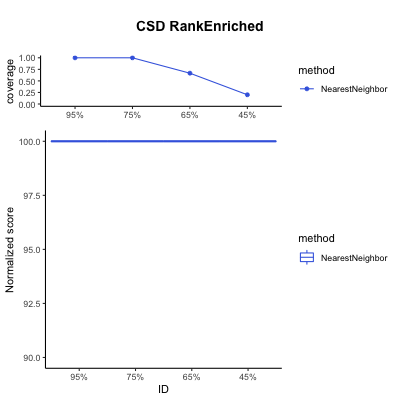

### CUT_Homeodomain_Enriched.png

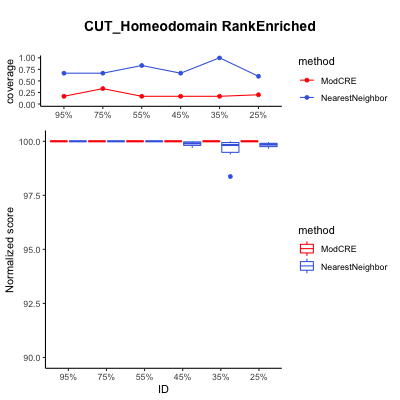

### CxxC_Enriched.png

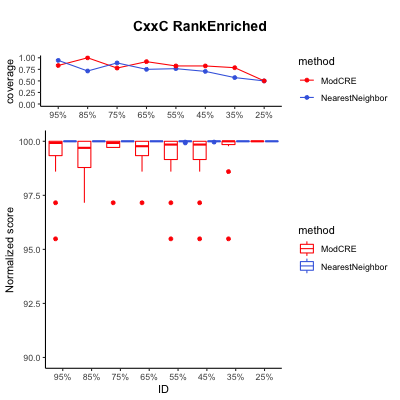

### DM_Enriched.png

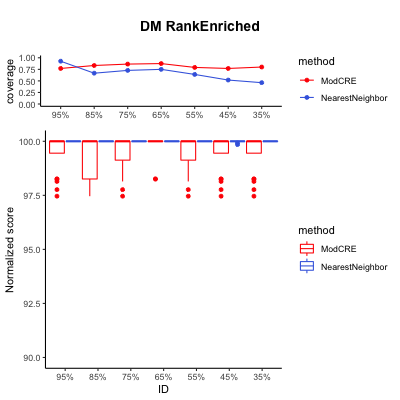

### Dof_Enriched.png

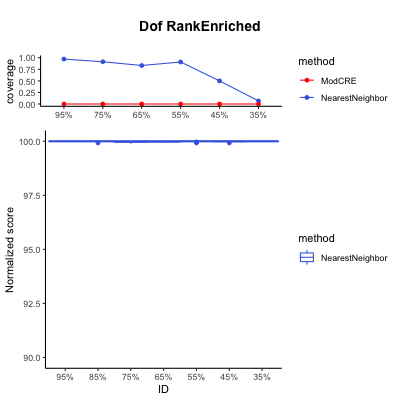

### E2F_Enriched.png

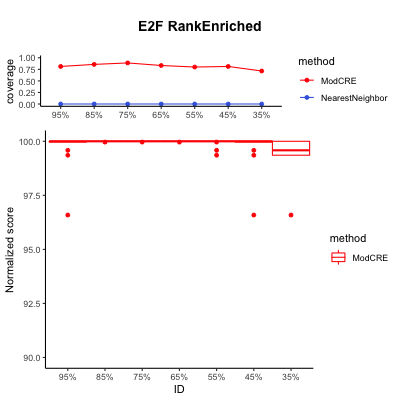

### EIN3_Enriched.png

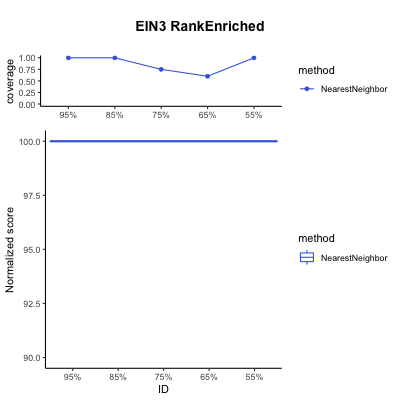

### Forkhead_Enriched.png

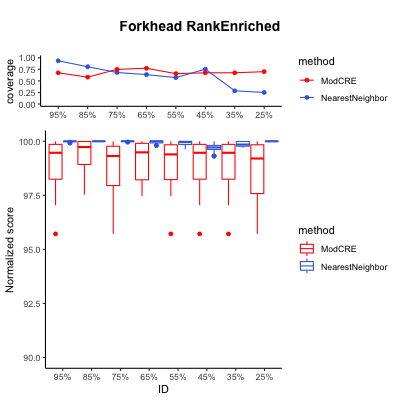

### GCM_Enriched.png

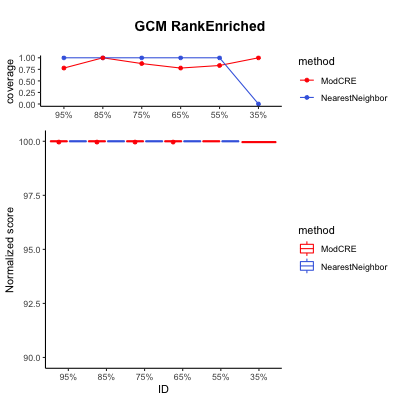

### GCR1_Enriched.png

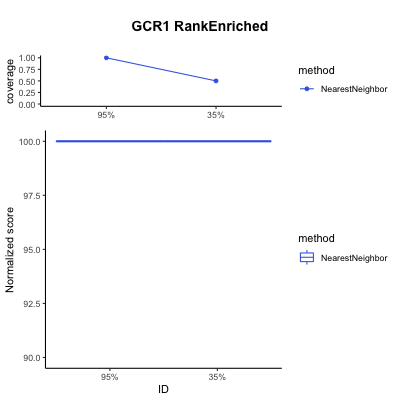

### global_Enriched.png

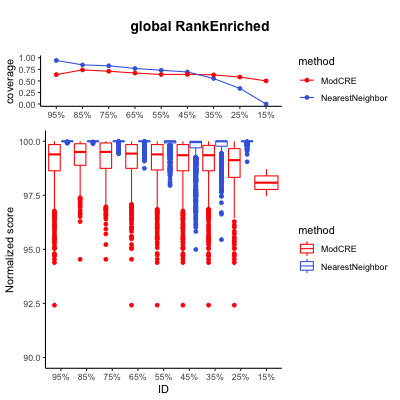

### Homeodomain_Enriched.png

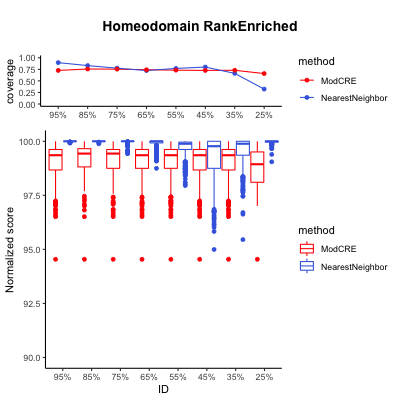

### Homeodomain_POU_Enriched.png

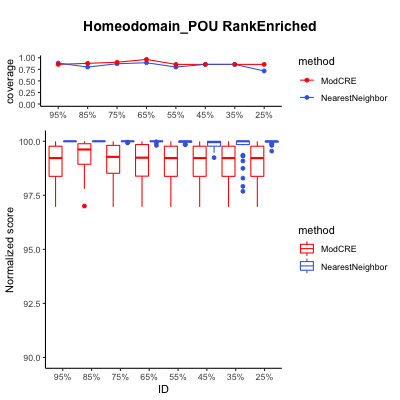

### HSF_Enriched.png

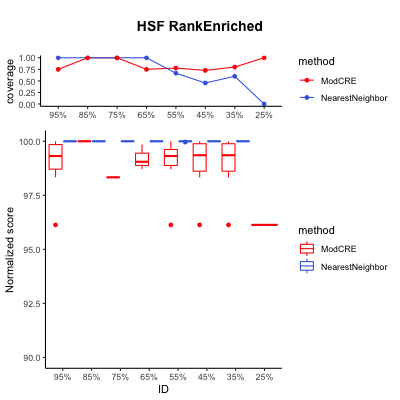

### IBD_Enriched.png

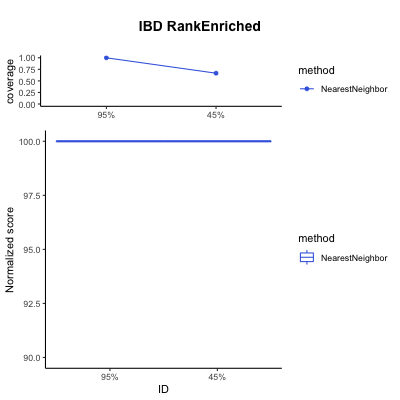

### IRF_Enriched.png

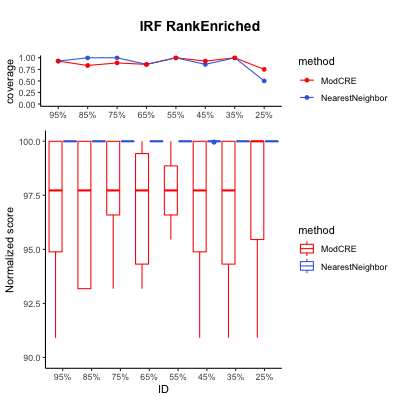

### LOB_Enriched.png

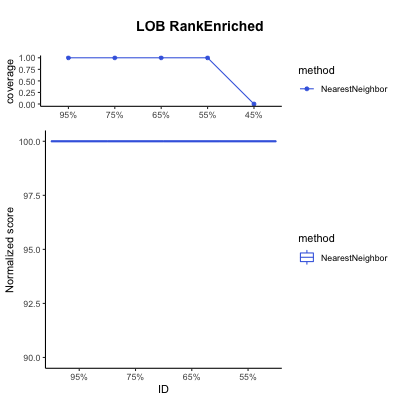

### MADS_box_Enriched.png

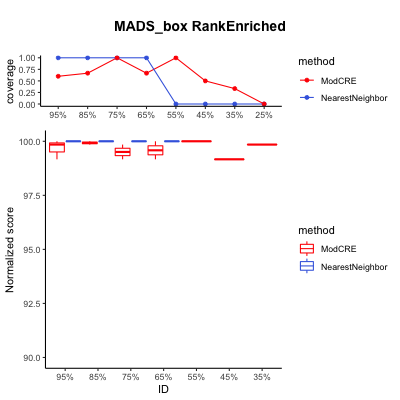

### NAC_NAM_Enriched.png

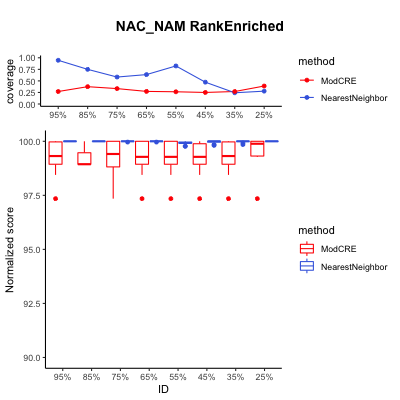

### Nuclear_receptor_Enriched.png

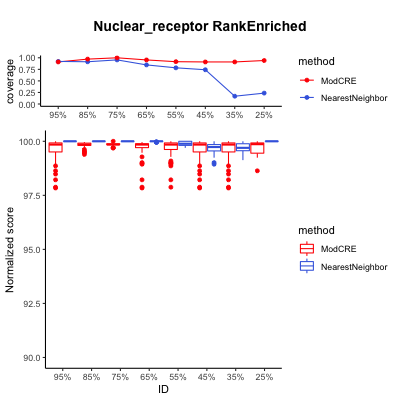

### POU_Enriched.png

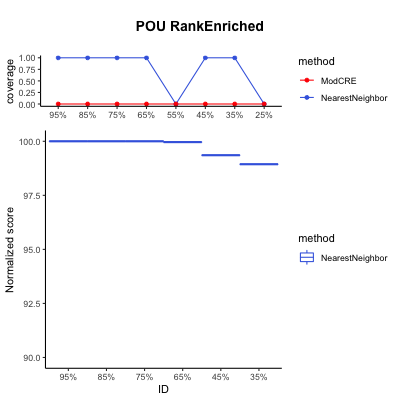

### Rel_Enriched.png

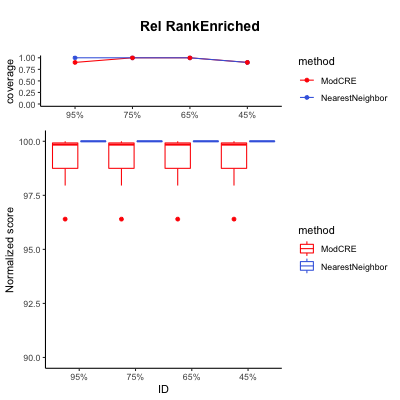

### RFX_Enriched.png

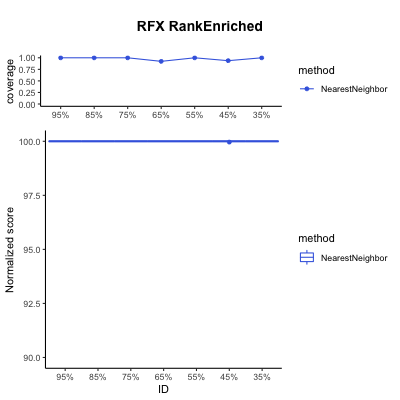

### SMAD_Enriched.png

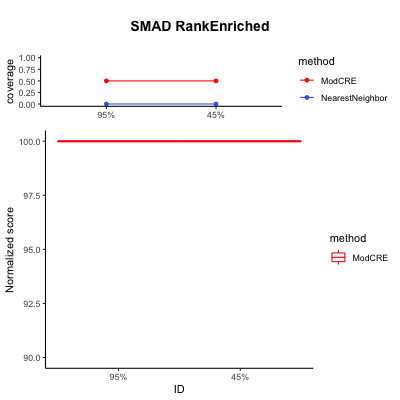

### Sox_Enriched.png

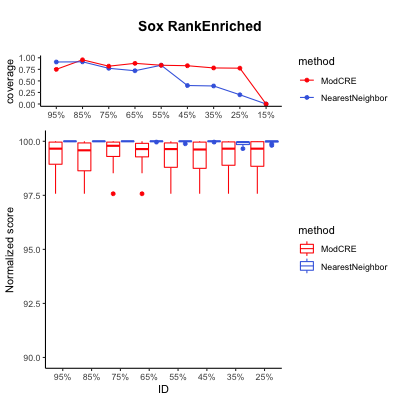

### TCP_Enriched.png

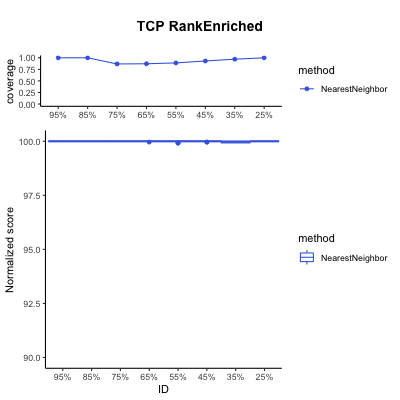

### TCR_CxC_Enriched.png

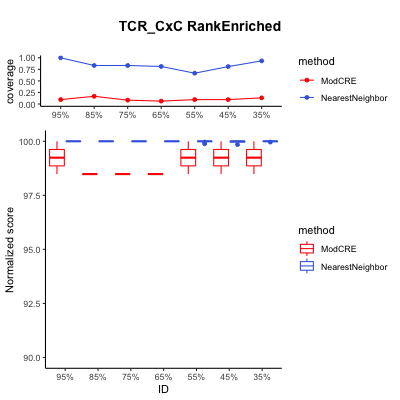

### TEA_Enriched.png

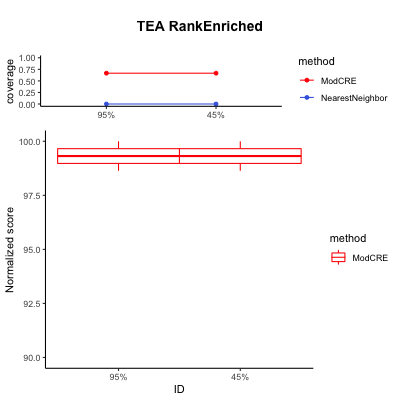

### WRC_Enriched.png

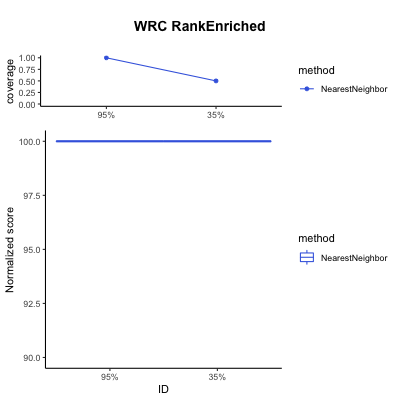
