## Supplementary figures and images for "Structure-based learning to model complex protein-DNA interactions and transcription-factor co-operativity in *cis*-regulatory elements"

### ABF1_Rank.png

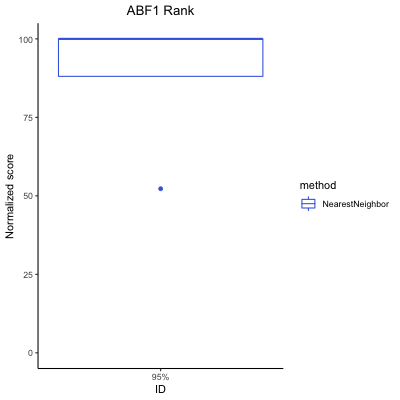

### AFT_Enriched.png

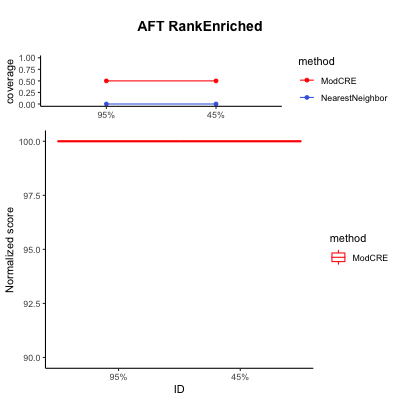

### AFT_Rank.png

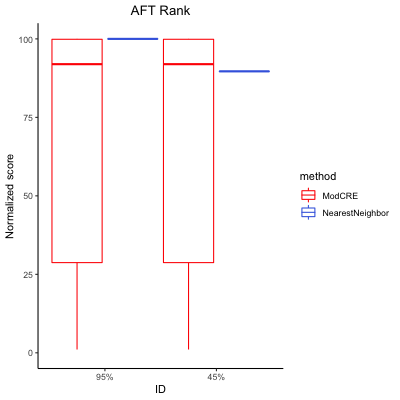

### AP2_B3_Enriched.png

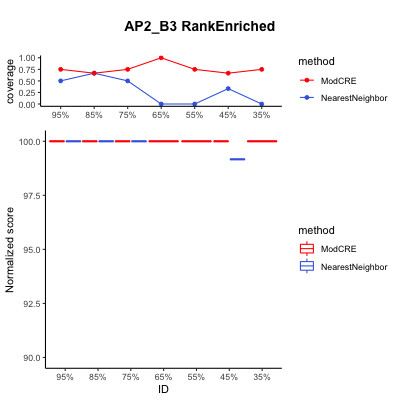

### AP2_B3_Rank.png

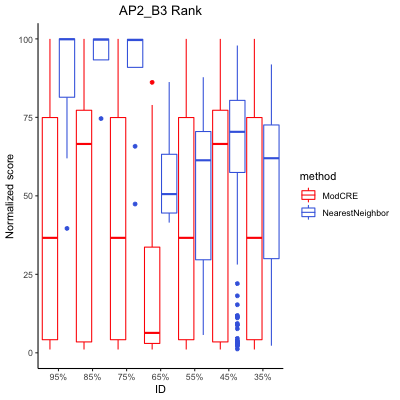

### AP2_Rank.png

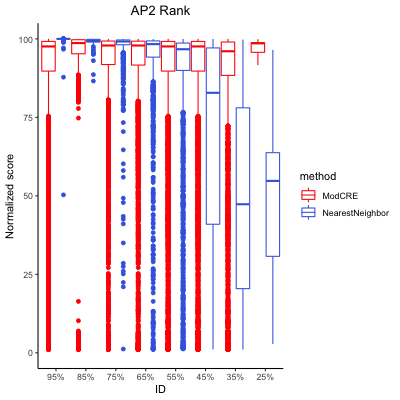

### AP-2_Rank.png

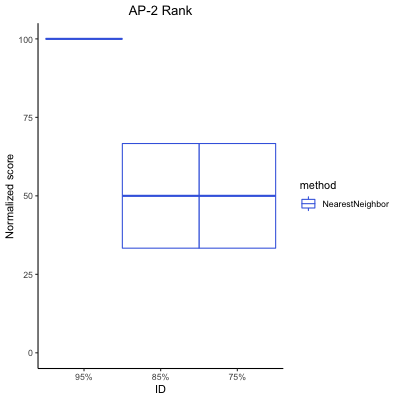

### APSES_Rank.png

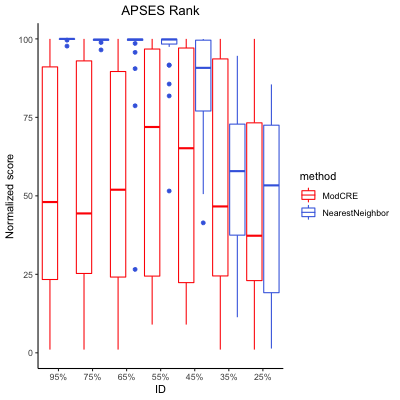

### ARID_BRIGHT_Rank.png

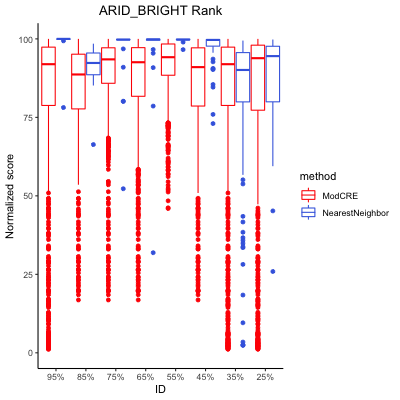

### AT_hook_Enriched.png

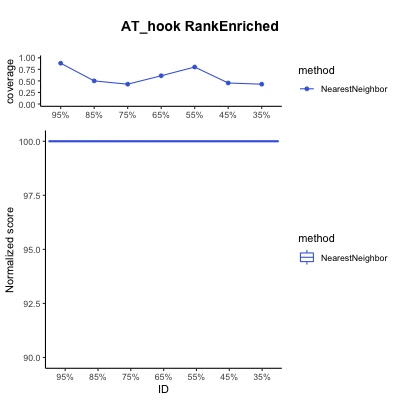

### AT_hook_Rank.png

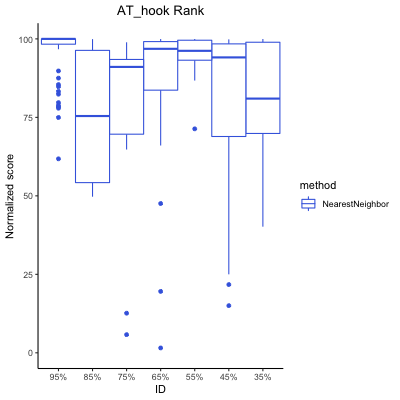

### B3_Rank.png

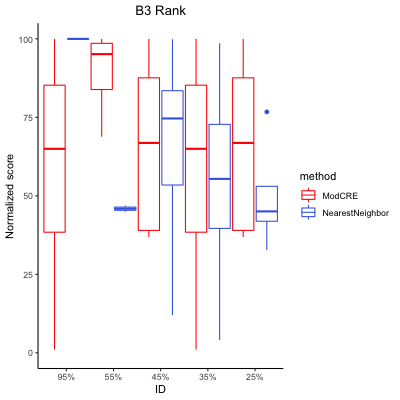

### bHLH_Enriched.png

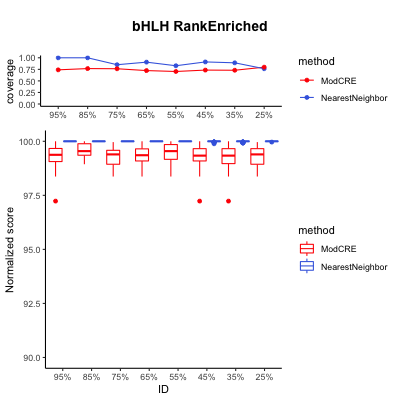

### bHLH_Rank.png

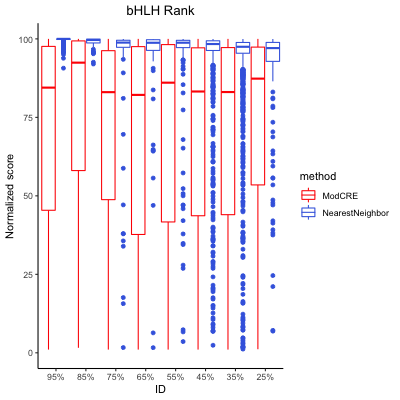

### Brinker_Rank.png

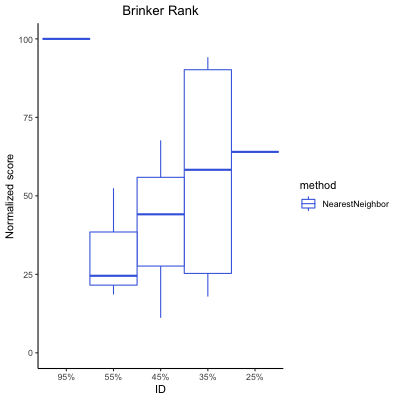

### bZIP_Enriched.png

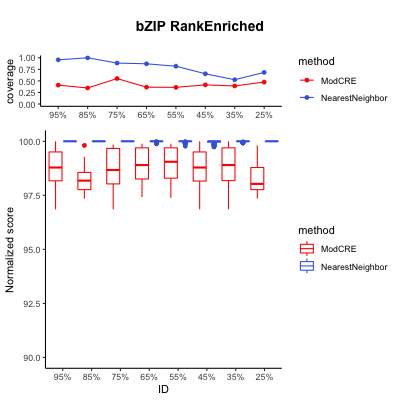

### bZIP_Rank.png

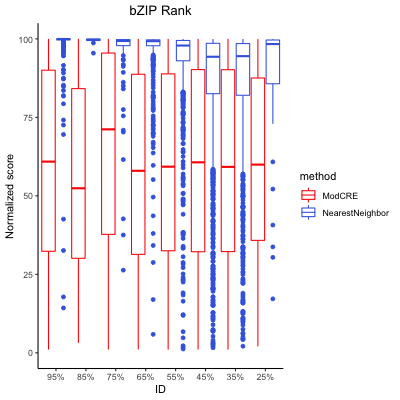

### C2H2_ZF_Homeodomain_Rank.png

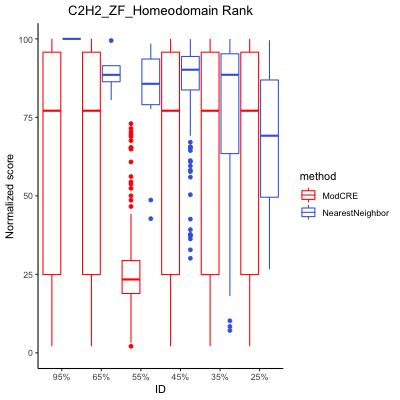

### C2H2_ZF_MADF_Rank.png

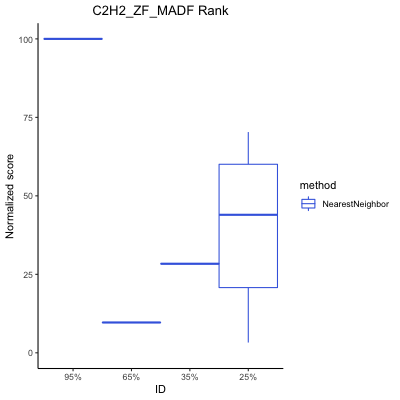

### C2H2_ZF_Myb_SANT_Enriched.png

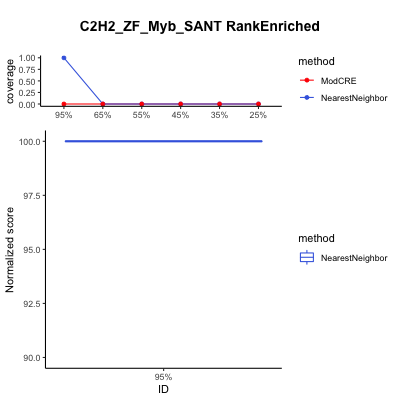

### C2H2_ZF_Myb_SANT_Rank.png

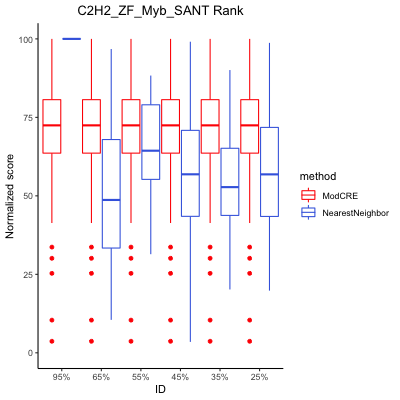

### C2H2_ZF_Rank.png

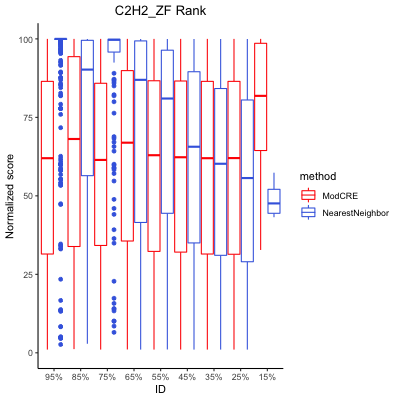

### C2H2_ZF_Zinc_cluster_Rank.png

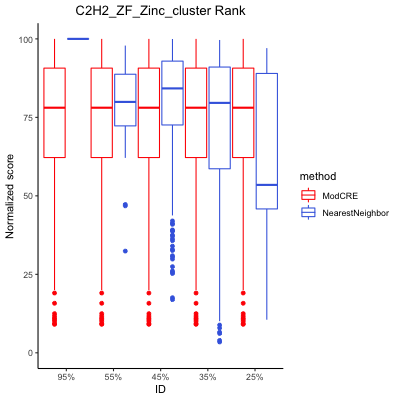

### C2HC_ZF_Rank.png

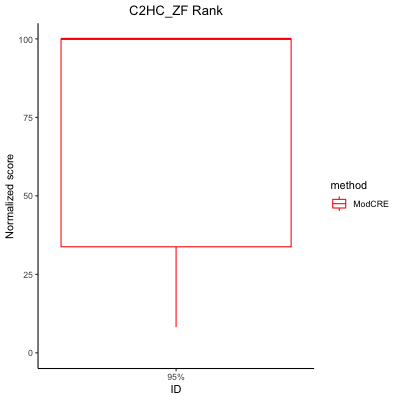

### CCCH_ZF_Rank.png

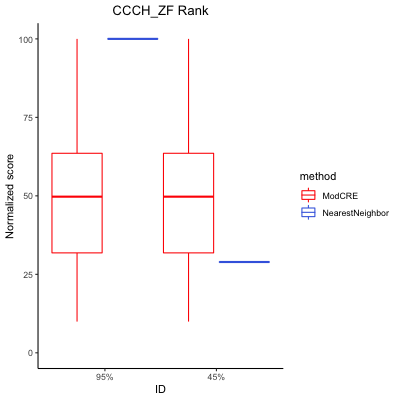

### CENPB_Enriched.png

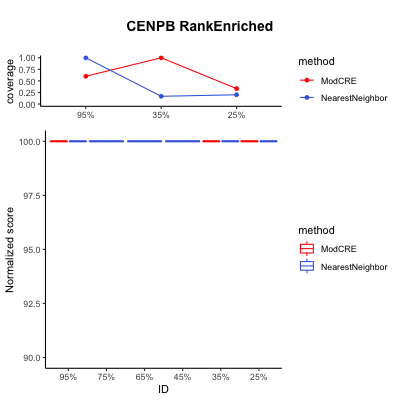

### CENPB_Rank.png

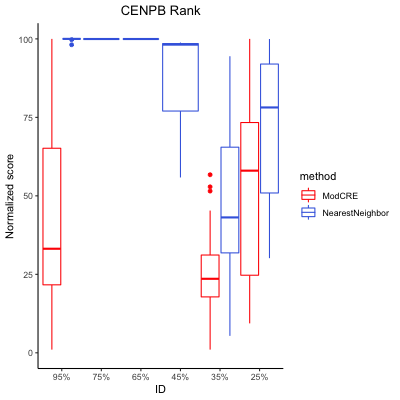

### CG-1_Rank.png

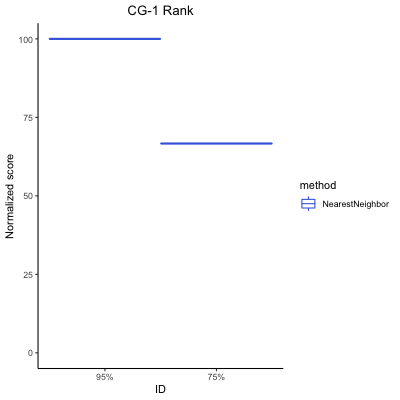

### CSD_Rank.png

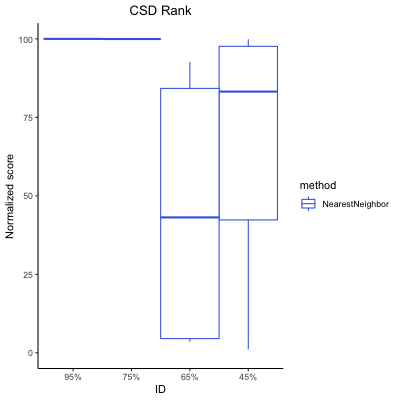

### CSL_Enriched.png

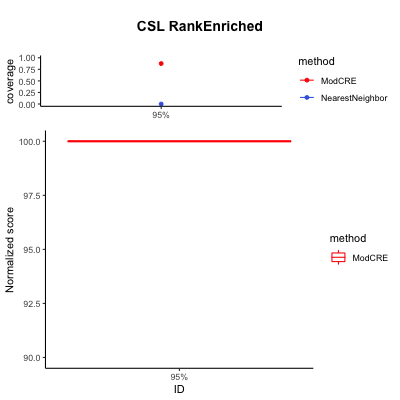

### CSL_Rank.png

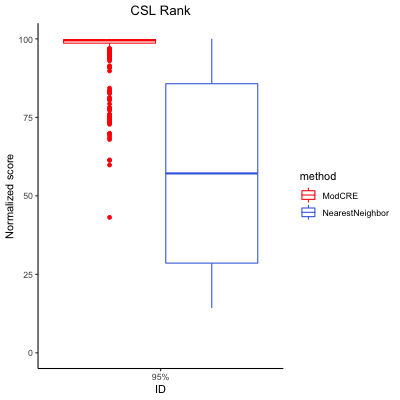

### CUT_Homeodomain_Rank.png

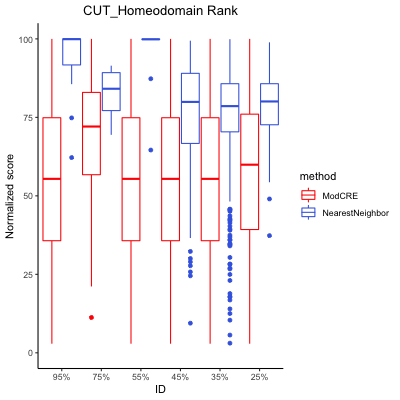

### CUT_Rank.png

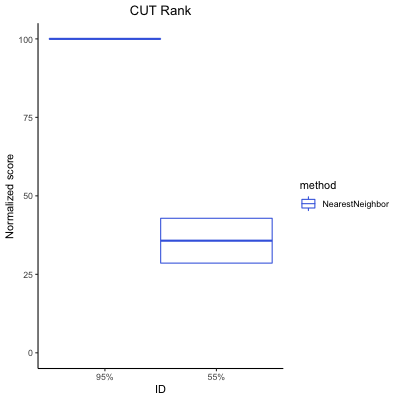

### CxxC_Rank.png

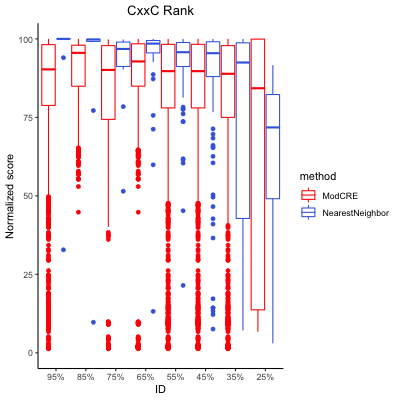

### DM_Rank.png

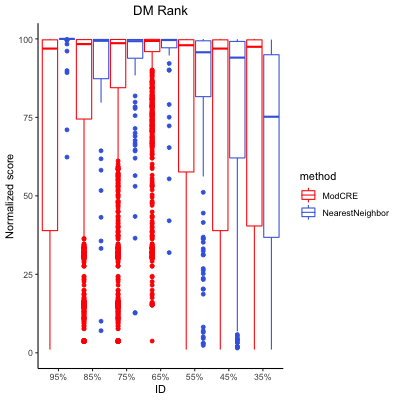

### Dof_Rank.png

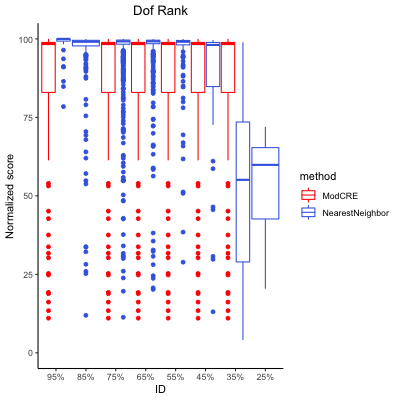

### E2F_Rank.png

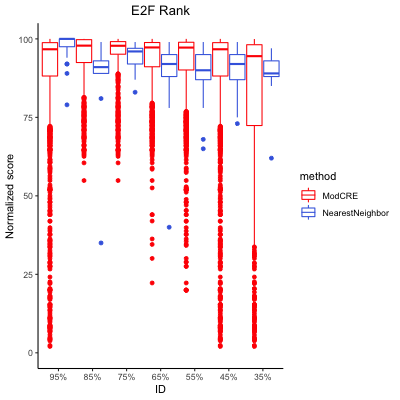

### EIN3_Rank.png

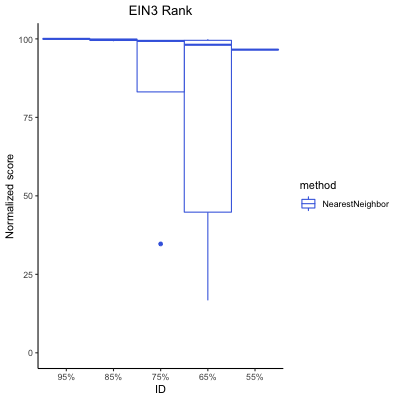

### Ets_Enriched.png

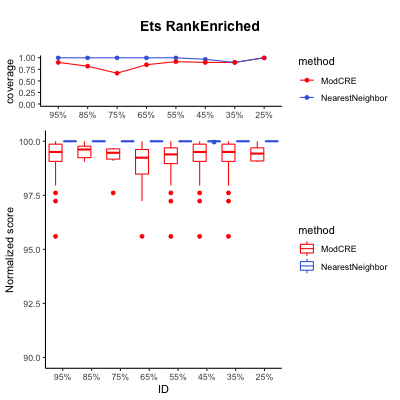

### Ets_Rank.png

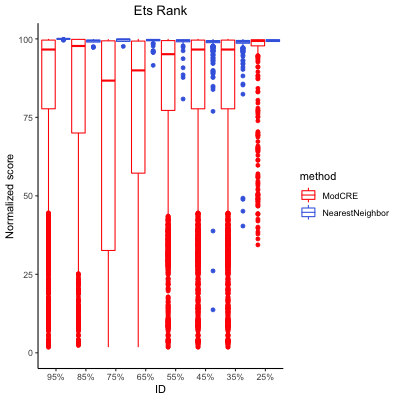

### Forkhead_Rank.png

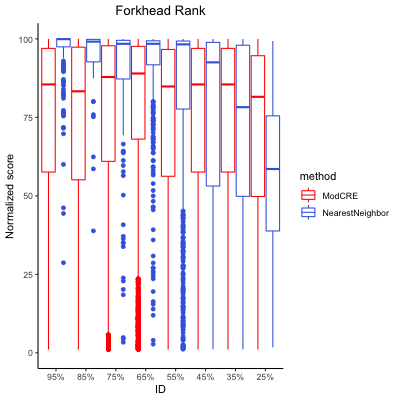

### GATA_Enriched.png

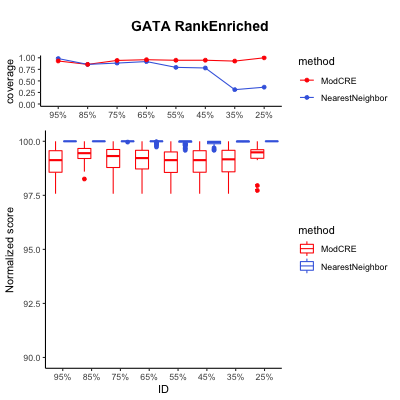

### GATA_Rank.png

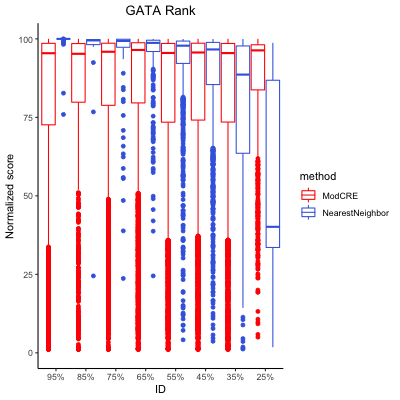

### GCM_Rank.png

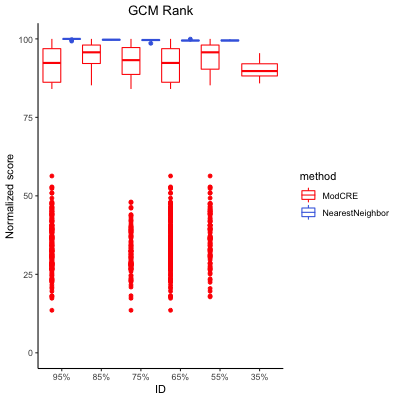

### GCR1_Rank.png

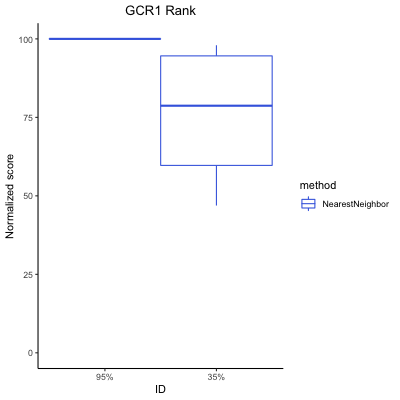

### global_Rank.png

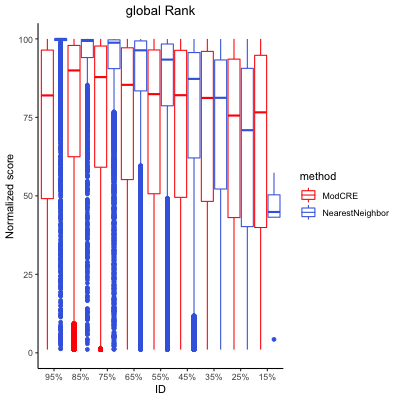

### Grainyhead_Rank.png

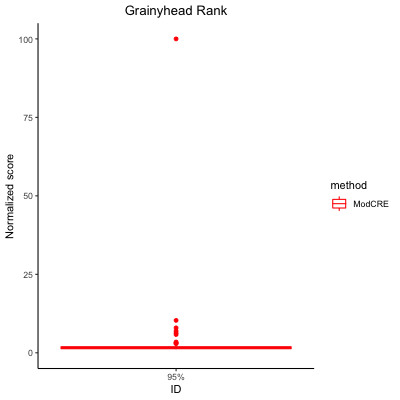

### Homeodomain_Paired_box_Enriched.png

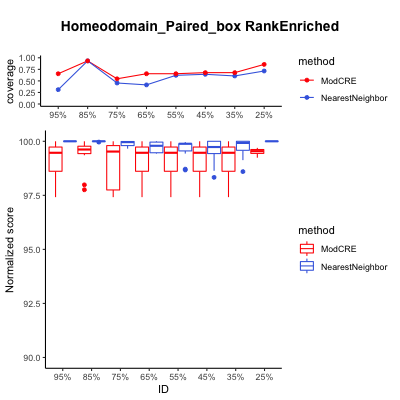

### Homeodomain_Paired_box_Rank.png

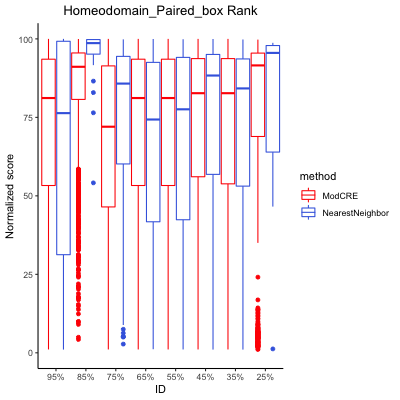

### Homeodomain_POU_Rank.png

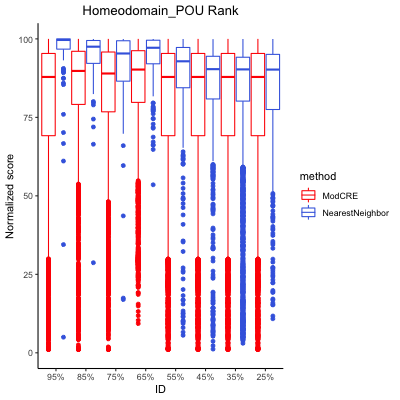

### Homeodomain_Rank.png

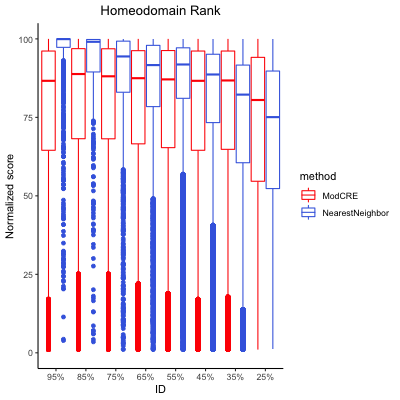

### HSF_Rank.png

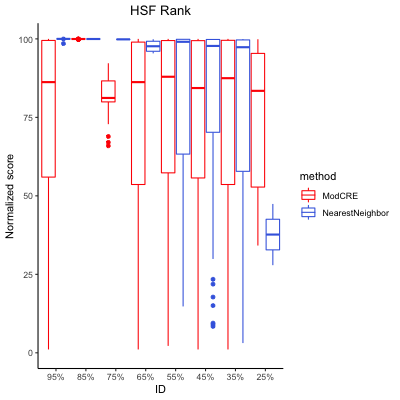

### IBD_Rank.png

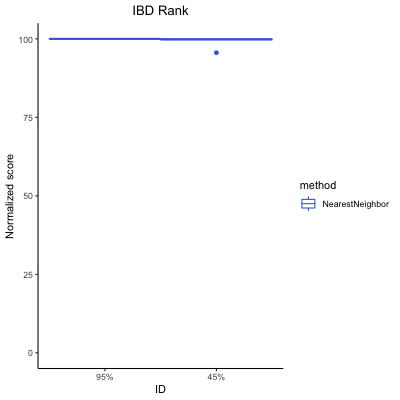

### IRF_Rank.png

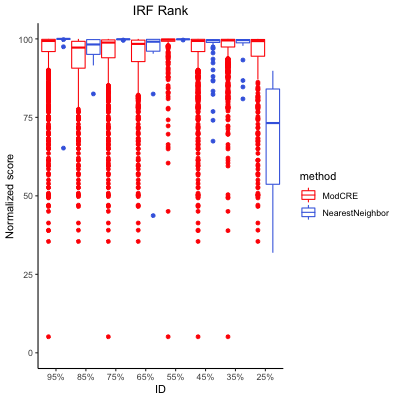

### LOB_Rank.png

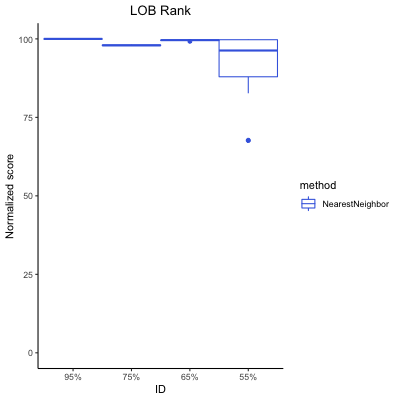

### MADF_Enriched.png

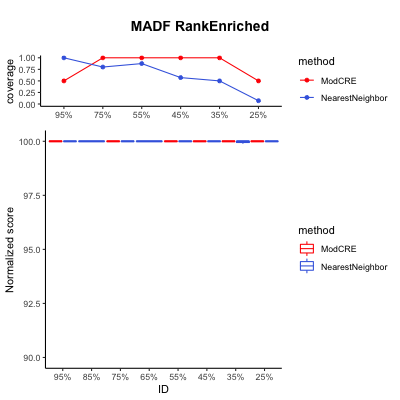

### MADF_Rank.png

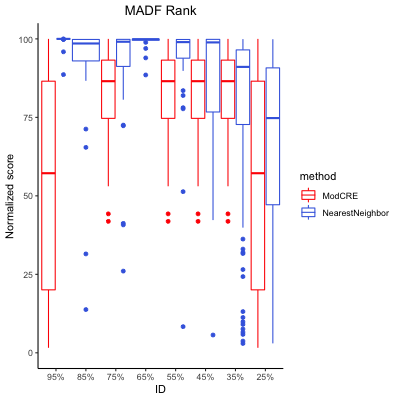

### MADS_box_Rank.png

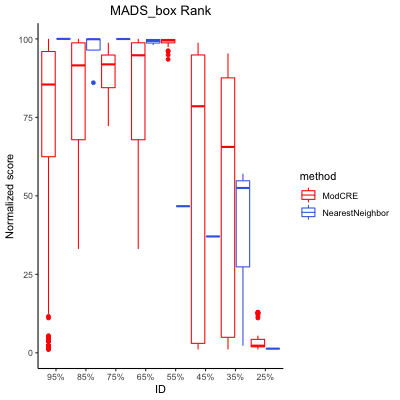

### MBD_Rank.png

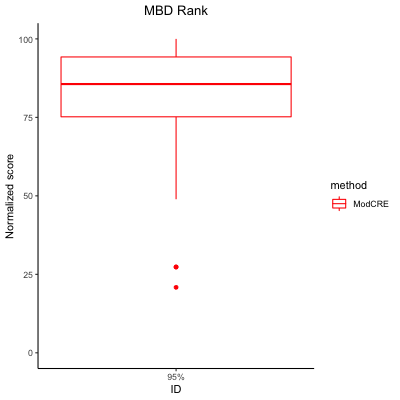

### Myb_SANT_Enriched.png

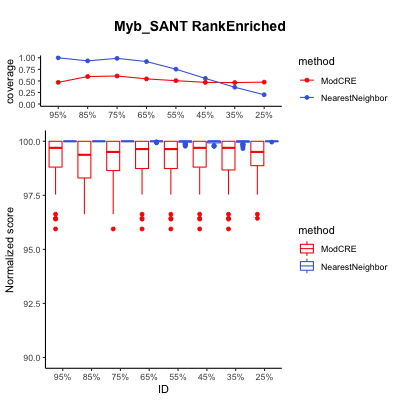

### Myb_SANT_Rank.png

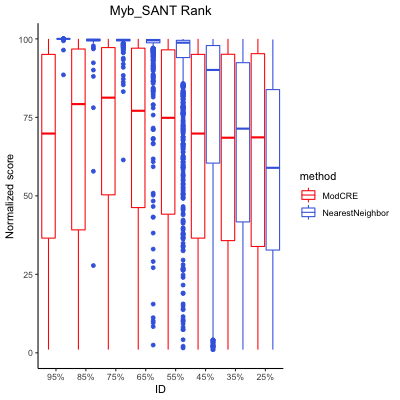

### NAC_NAM_Rank.png

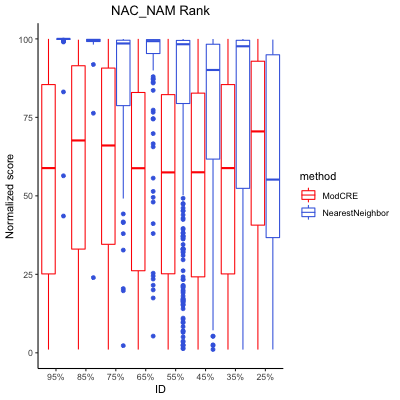

### Ndt80_PhoG_Enriched.png

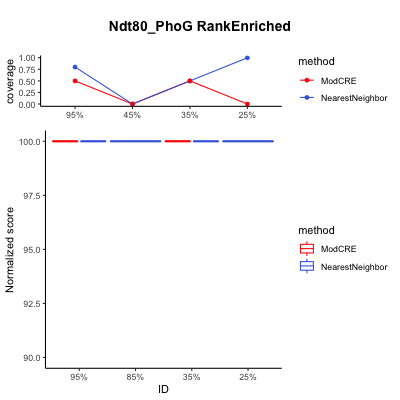

### Ndt80_PhoG_Rank.png

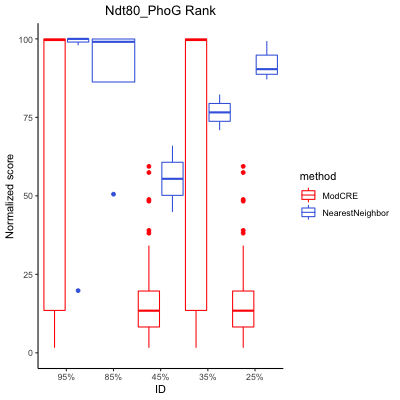

### Nuclear_receptor_Rank.png

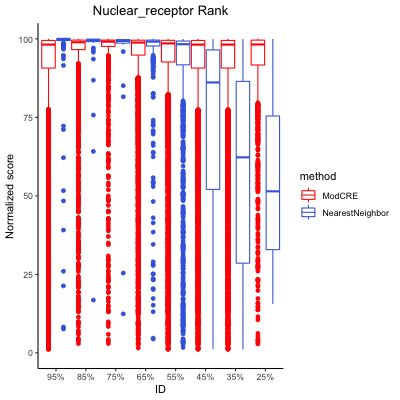

### Paired_box_Enriched.png

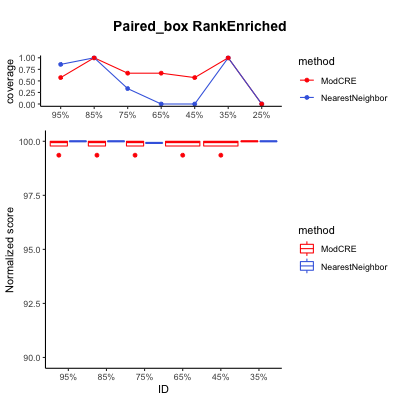

### Paired_box_Rank.png

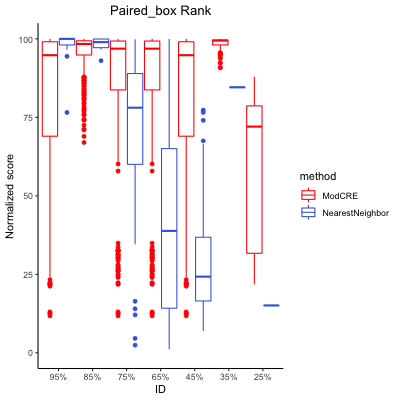

### Pipsqueak_Enriched.png

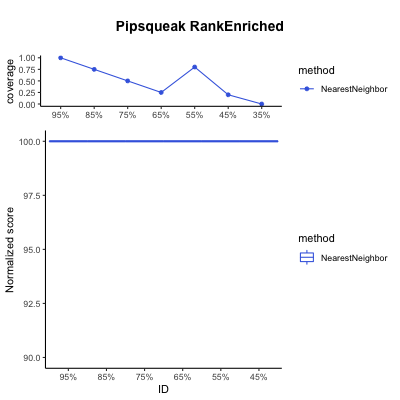

### Pipsqueak_Rank.png

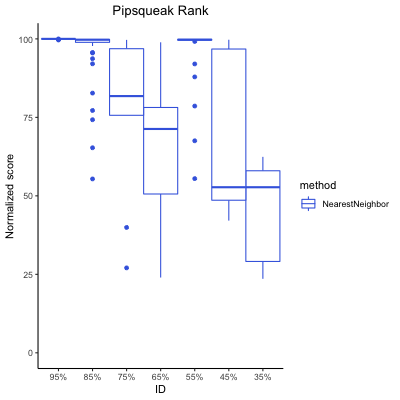

### POU_Rank.png

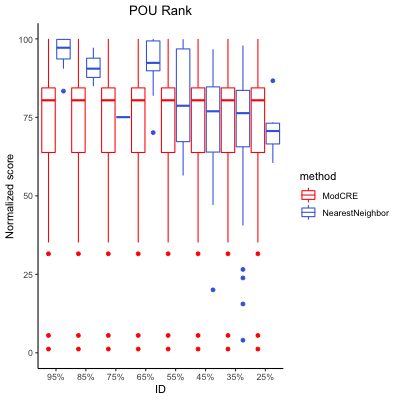

### Prospero_Rank.png

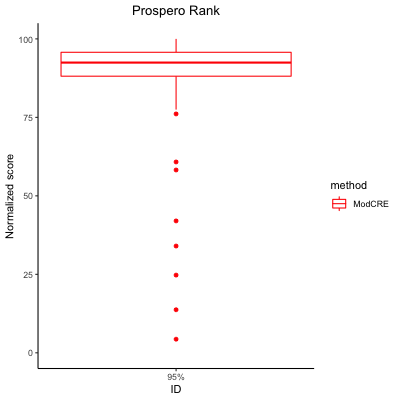

### Rap1_Enriched.png

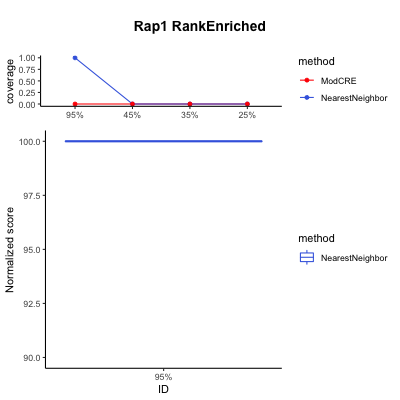

### Rap1_Rank.png

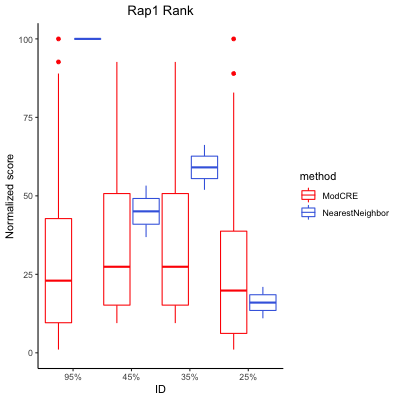

### Rel_Rank.png

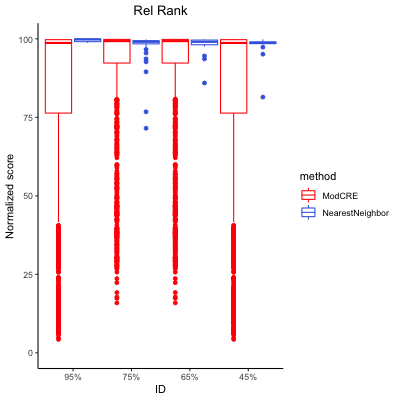

### RFX_Rank.png

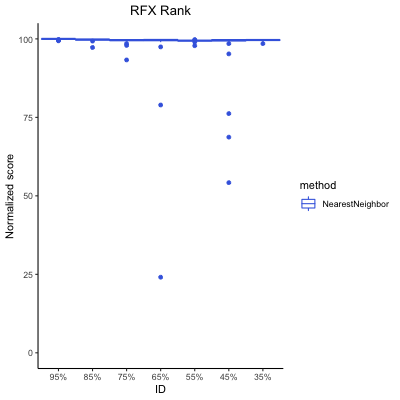

### Runt_Enriched.png

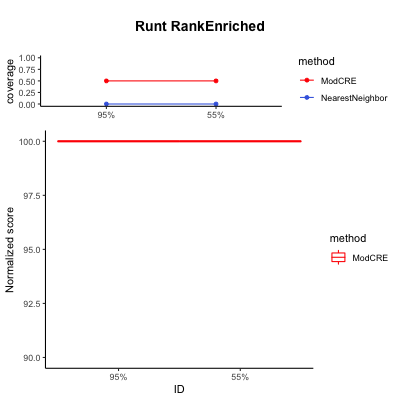

### Runt_Rank.png

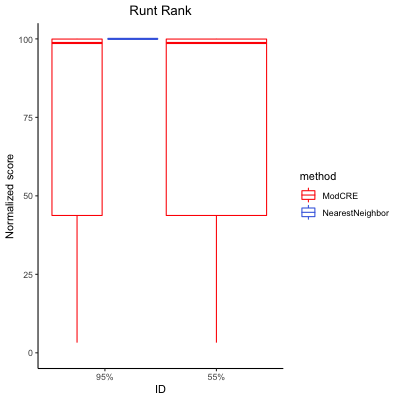

### SAND_Enriched.png

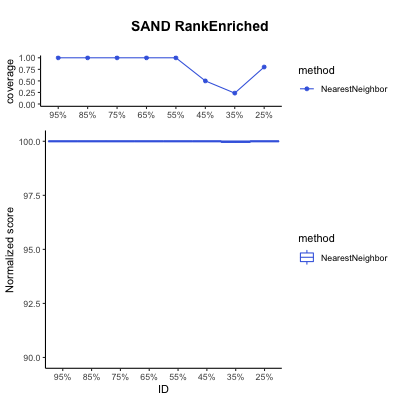

### SAND_Rank.png

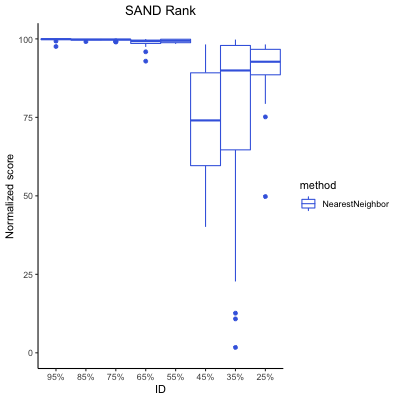

### SBP_Enriched.png

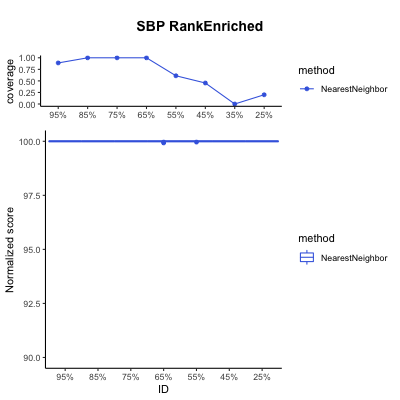

### SBP_Rank.png

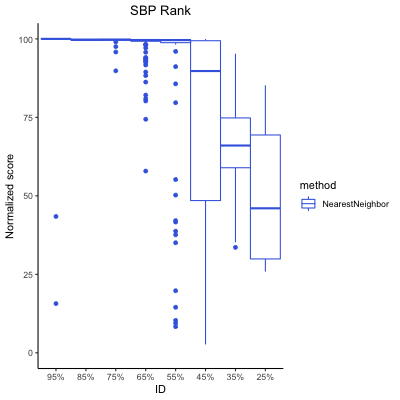

### SMAD_Rank.png

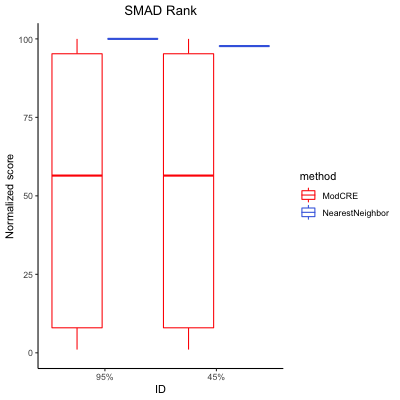

### Sox_Rank.png

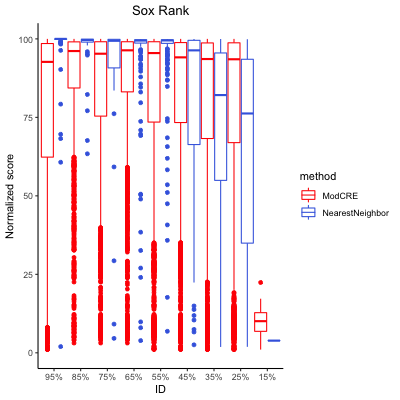

### STE_Rank.png

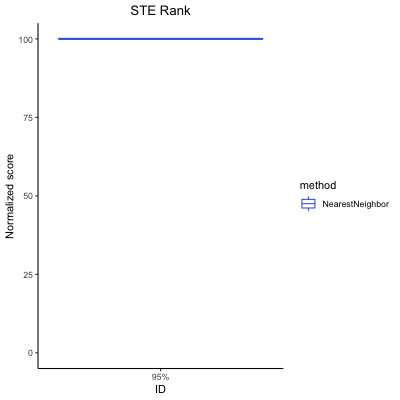

### Storekeeper_Enriched.png

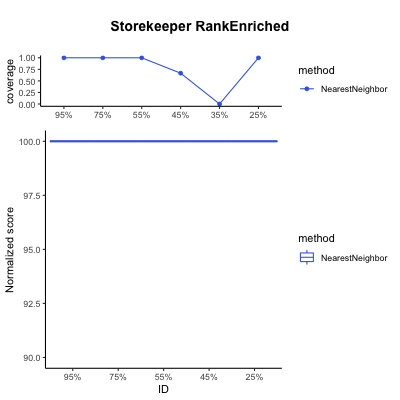

### Storekeeper_Rank.png

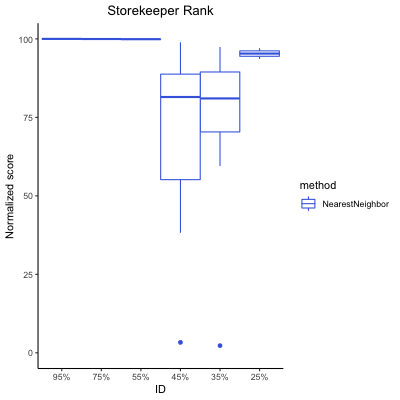

### T-box_Enriched.png

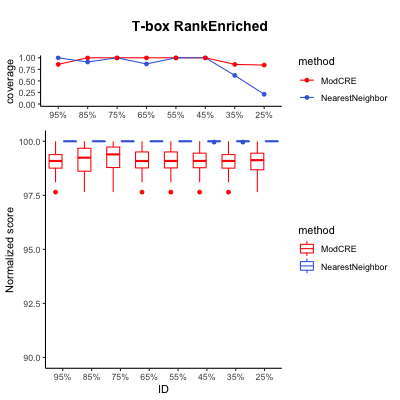

### T-box_Rank.png

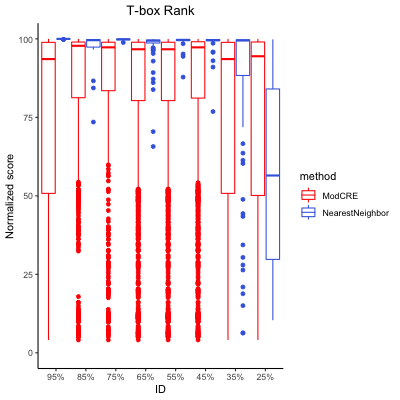

### TBP_Enriched.png

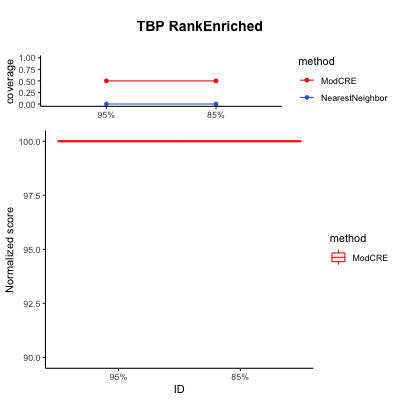

### TBP_Rank.png

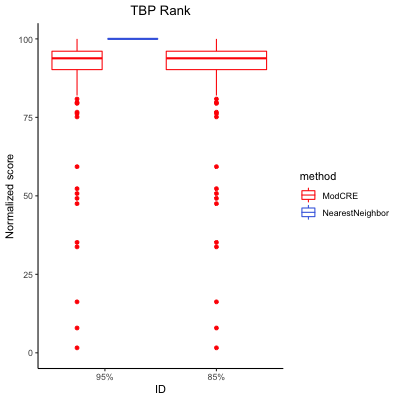

### TCP_Rank.png

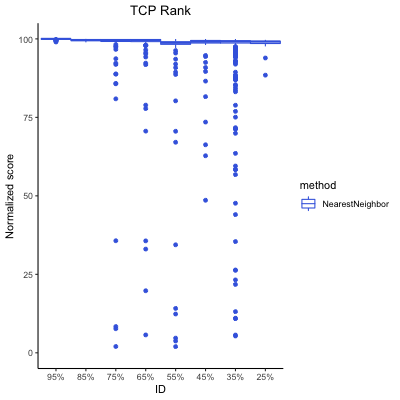

### TCR_CxC_Rank.png

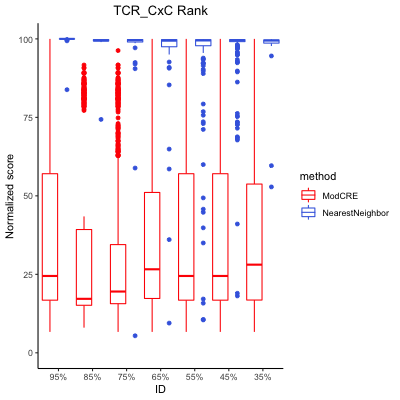

### TEA_Rank.png

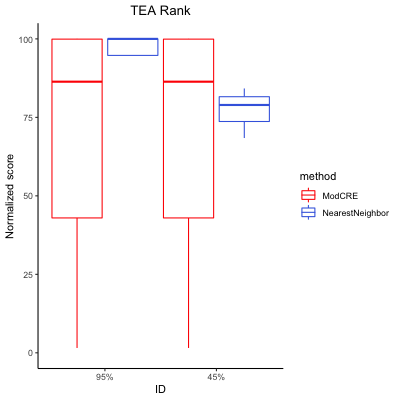

### THAP_finger_Enriched.png

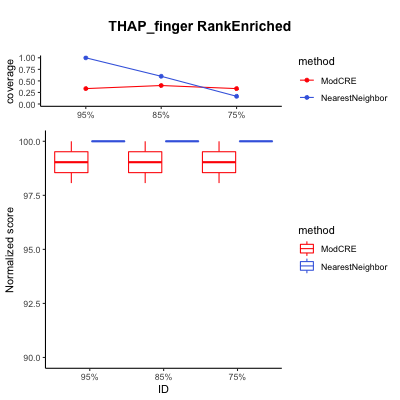

### THAP_finger_Rank.png

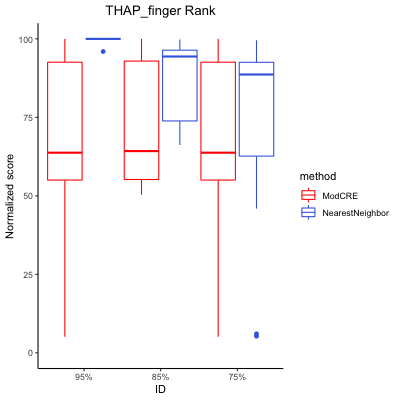

### WRC_Rank.png

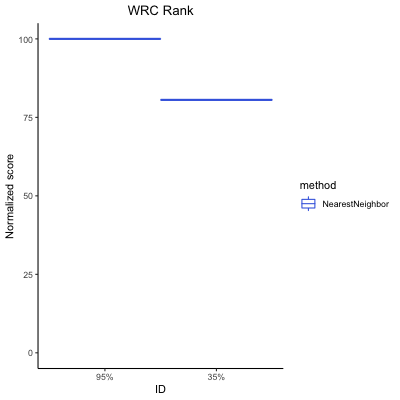

### WRKY_Enriched.png

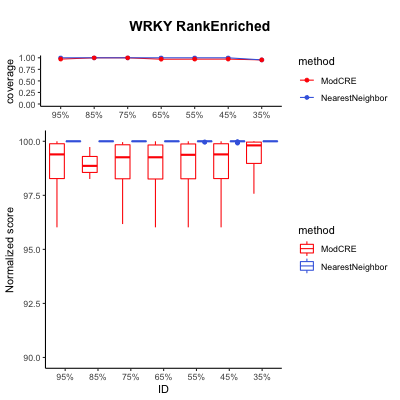

### WRKY_Rank.png

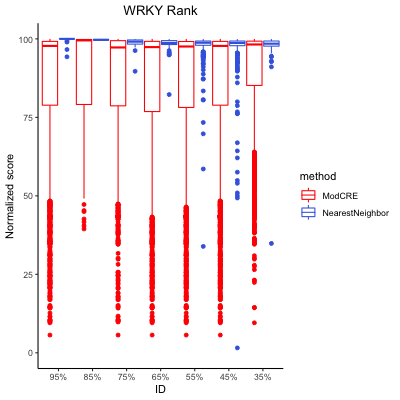

### Zinc_cluster_Enriched.png

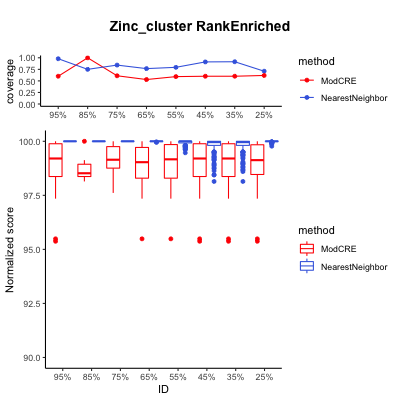

### Zinc_cluster_Rank.png

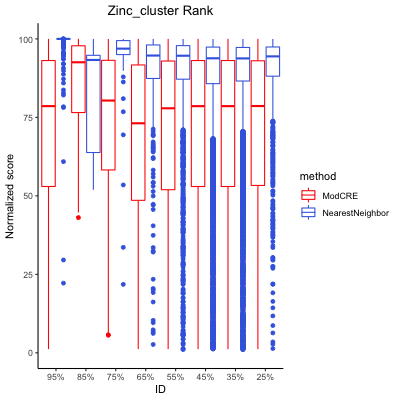
